# The fitness cost of spurious phosphorylation

**DOI:** 10.1101/2023.10.08.561337

**Authors:** David Bradley, Alexander Hogrebe, Rohan Dandage, Alexandre K Dubé, Mario Leutert, Ugo Dionne, Alexis Chang, Judit Villén, Christian R Landry

## Abstract

The fidelity of signal transduction requires the binding of regulatory molecules to their cognate targets. However, the crowded cell interior risks off-target interactions between proteins that are functionally unrelated. How such off-target interactions impact fitness is not generally known, but quantifying this is required to understand the constraints faced by cell systems as they evolve. Here, we use the model organism *S. cerevisiae* to inducibly express tyrosine kinases. Because yeast lacks *bona fide* tyrosine kinases, most of the resulting tyrosine phosphorylation is spurious. This provides a suitable system to measure the impact of artificial protein interactions on fitness. We engineered 44 yeast strains each expressing a tyrosine kinase, and quantitatively analysed their phosphoproteomes. This analysis resulted in ∼30,000 phosphosites mapping to ∼3,500 proteins. Examination of the fitness costs in each strain revealed a strong correlation between the number of spurious pY sites and decreased growth. Moreover, the analysis of pY effects on protein structure and on protein function revealed over 1000 pY events that we predict to be deleterious. However, we also find that a large number of the spurious pY sites have a negligible effect on fitness, possibly because of their low stoichiometry. This result is consistent with our evolutionary analyses demonstrating a lack of phosphotyrosine counter-selection in species with *bona fide* tyrosine kinases. Taken together, our results suggest that, alongside the risk for toxicity, the cell can tolerate a large degree of non-functional crosstalk as interaction networks evolve.

## Introduction

Signalling pathways are often represented as perfectly specific systems, with a linear chain of directed interactions linking the primary signal to an effector enzyme or transcription factor. However, a number of biophysical and evolutionary considerations imply the existence of non-functional interactions in a dense protein network (Levy *et al*, 2009). Firstly, signalling interactions are often mediated by short linear motifs (SLiMs) that are degenerate in sequence and therefore may be created or destroyed by a small number of substitutions (Davey *et al*, 2015; Kliche *et al*, 2023). Secondly, members of a protein family may share protein folds and binding interfaces that risk illicit interactions between proteins homologous to the functional binding partners (Nocedal & Laub, 2022; McClune & Laub, 2020). This risk is exacerbated when one considers the thousands of proteins that may interact in the cytoplasm (Levy *et al*, 2009), and the crowded cellular environment through which signalling occurs (Delarue *et al*, 2018; Nussinov *et al*, 2021; Li *et al*, 2021). While several mechanisms (co-expression, co-localisation, scaffolding, PTMs) can enhance functional specificity (Pawson, 2004; Scott & Pawson, 2009; Good *et al*, 2011; Miller & Turk, 2018), there is also evidence from PTM data for spurious interactions that contribute biological ‘noise’ to signalling networks (Levy *et al*, 2012; Landry *et al*, 2013; Hornbeck *et al*, 2015; James *et al*, 2018a, 2018b).

The optimum that a cell system can achieve is also influenced by evolutionary constraints at the level of the population. That is, whether a deleterious mutation can be removed by purifying selection and a beneficial one can reach fixation. The efficacy of selection is thus linked to the size of the population (higher in large populations), the strength of the selection pressure, and in the context of signalling the nature of the bias for a spontaneous mutation to either gain or lose a physical interaction (Lynch & Hagner, 2015; Lynch *et al*, 2016). However, understanding how these constraints shape the evolution of signalling networks requires the fitness effect of spurious interactions to first be quantified (Levy *et al*, 2009).

In this context we consider phosphotyrosine signalling in the budding yeast *Saccharomyces cerevisiae*. Yeast lacks classical tyrosine kinases, SH2 domains, or PTB domains, and the small amount of phosphotyrosine in the proteome is thought to derive from dual-specificity kinases (Manning *et al*, 2002; Pincus *et al*, 2008; Lim & Pawson, 2010; Kaneko *et al*, 2012; Leutert *et al*, 2023). Therefore, the heterologous expression of tyrosine kinases in this species is expected to generate many phosphotyrosine (pY) sites that are spurious by definition, some of which could have deleterious consequences. Indeed, expression of the hyperactive viral kinase v-SRC leads to toxicity in yeast (Brugge *et al*, 1987; Kornbluth *et al*, 1987) – an established result that has been replicated several times (Xu & Lindquist, 1993; Boschelli *et al*, 1993; Florio *et al*, 1994; Trager & Martin, 1997) and demonstrated in the fission yeast *Schizosaccharomyces pombe* for chicken c-Src (Superti-Furga *et al*, 1993). In more recent years, this relationship between kinase activity and toxicity has been used as the basis for fitness-based deep mutational scanning (DMS) assays of the tyrosine kinase domain (Ahler *et al*, 2019; Chakraborty *et al*, 2023).

The advantage of this system for the study of spurious interactions is that protein phosphorylation across the proteome can be assessed and quantified using mass spectrometry-based phosphoproteomics (Olsen *et al*, 2006; Villén *et al*, 2007; Rikova *et al*, 2007; Leutert *et al*, 2019). This phosphoproteomic approach has been used for the modelling of human tyrosine kinase specificity amid a minimal background of native pY signalling in *S. cerevisiae (Corwin et al, 2017)*. More recently, a large panel of human kinases has been tested for fitness in yeast across several conditions and used to assay for phosphorylation-dependent interactions between human proteins (Jehle *et al*, 2022). Here we express several human tyrosine kinases in yeast alongside their kinase-dead counterparts to specifically focus on the effect of kinase activity. In each case, we assess the differences in fitness and phosphorylation between the WT tyrosine kinase and kinase-dead strains. This comparison allows us to relate spurious protein-protein-interactions (PPIs) to fitness. For this purpose we also take advantage of Alphafold2-derived protein structure models and recent bioinformatic advances to perform proteome-wide variant effect prediction (VEP) for tyrosine phosphorylation on the basis of protein structure and protein conservation. Finally, with this phosphoproteomic data as a baseline of what could be potentially phosphorylated by these kinases in the absence of selection for function, we perform a deep evolutionary analysis of the pY sites and determine whether spurious Y phosphosites were selected against in species with *bona fide* tyrosine kinases.

## Results

### Expression of human kinases in yeast and their activity

We expressed 24 Y kinases and 7 S/T kinases in yeast by inserting the coding sequence into a landing pad in the yeast genome where the coding sequence is regulated by an inducible promoter **(Figure 1a, Table S1).** The set of tyrosine kinases included non-receptor tyrosine kinases that had been previously studied by (Corwin *et al*, 2017) and shown to be active (ABL2, SRC, SYK, BMX, FES, FRK, FYN, LYN, SRMS, and TNK1), three additional non-receptor tyrosine kinases (ABL1, LCK, TEC), and several variants of vSRC (n=13) that span a range of kinase activities (Ahler *et al*, 2019). We also included several kinase domains of receptor tyrosine kinases (RTK) (EPHA1, EPHA2, EPHA3, EPHB3, EPHB1, EPHB4, FGFR2, FGFR3, MERTK, and MET) and S/T kinases (IRAK4, NEK6, NEK7, RAF1, TBK1, TLK2, VRK1). The S/T kinases were selected because they had no orthologs in yeast, their peptide recognition profiles overlapped minimally with the yeast phosphoproteome, clones were readily available, and their activity can be increased by autophosphorylation (Beenstock *et al*, 2016). All oligonucleotides used for the construction of these strains are given in **Table S2.**

**Figure 1.**
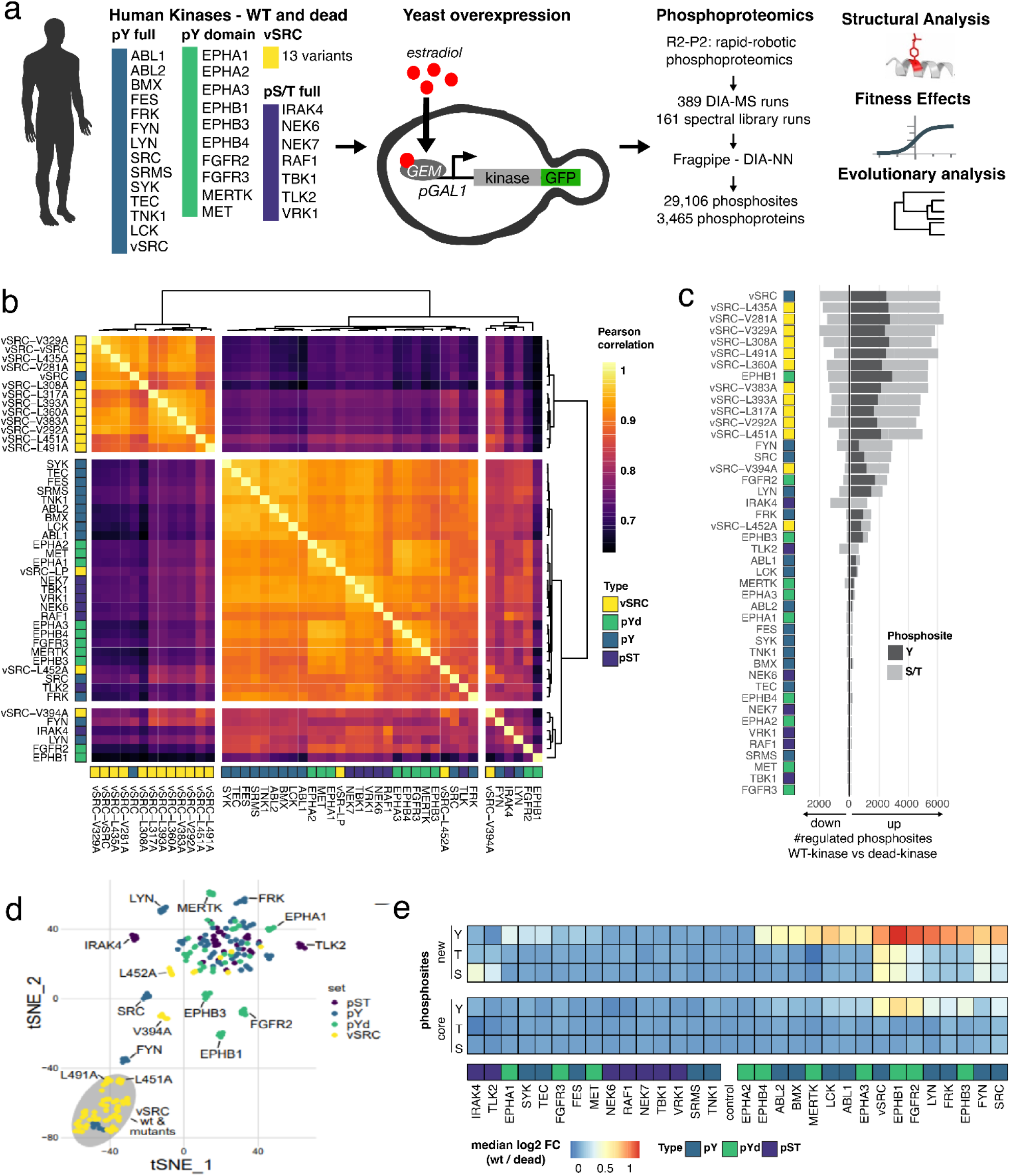
Expression of human tyrosine kinases in yeast and detection of their substrates using mass spectrometry. **a)** Inducible expression of human tyrosine kinases from a genomic landing pad in *S. cerevisiae*, followed by data-independent acquisition (DIA) mass spectrometry. After phosphoproteomics, the impact of phosphorylation on protein structure, on fitness, and evolution is analysed. pY: full-length tyrosine kinase, pYd: tyrosine kinase domain, vSRC: WT VSRC and its mutants, pS/pT: full-length serine and threonine kinases. **b)** Correlated phosphorylation profiles (Pearson’s correlation coefficient) between all kinases tested, based upon the median phosphosite intensity (pS/pT/pY) across technical replicates. **c)** Number of up- and downregulated pY (dark grey) and pS/pT (light grey) sites per kinase. Up- and down-regulation for each WT kinase is with respect to the kinase-dead mutant. **d)** Separation of phosphorylation profiles (per kinase) in two dimensions using the tSNE dimensionality-reduction method **e)** Relative phosphosite abundance log2 (WT/dead) for each kinase, with respect to the phosphoacceptor identity (S, T, Y) and whether or not the phosphosite is a member of the core phosphoproteome in *S. cerevisiae* that is found to be phosphorylated in many conditions (Leutert *et al*, 2023).

The expression and activity of each tyrosine kinase or kinase domain (tagged with GFP) was examined by Western blot, using the protein extracts of yeast cells induced or not with estradiol **(Figure S1-S2)**. Detectable expression (anti-GFP) was found for most kinases tested (14/24). The activity of some of the enzymes was validated by monitoring the overall tyrosine phosphorylation of proteins (4G10; pan anti-phosphotyrosine), using kinase-dead (KD) point mutants as controls (10/24). In addition, we analysed the expression and activity of the different vSRC mutants used in this study. We found wild-type vSRC to be well expressed and active in yeast cells. Of the 13 mutants tested, 11 had observable phosphotyrosine signal and 9 were sufficiently expressed to detect GFP bands at the correct molecular weights. Since most kinases were detected by Western blotting, we considered all kinases for downstream experiments even if their expression could be low. Almost all (∼95%) of the tested kinases were detected in their phosphorylated form by mass spectrometry (section below), thus providing additional evidence of their expression in this system.

### Effects of human kinase expression on the yeast phosphoproteome

Phosphorylation in all strains was assessed across 5 biological replicates using data-independent acquisition mass spectrometry (DIA-MS) **(Figure 1a).** Yeast proteins were extracted via bead-beating, reduced and alkylated, and digested into peptides by trypsin. Phosphopeptides were enriched by immobilised metal ion affinity chromatography (IMAC) using the automated R2-P2 method (Leutert *et al*, 2019) and MS-based measurement was performed using an Orbitrap Exploris 480. Across the 389 samples we identified a total of 42,653 unique phosphopeptides comprising 29,106 unique phosphosites **(Figure S3)**. Data quality was high, with a median phosphopeptide enrichment efficiency of 97.3% and a median number of 9,585 identified phosphopeptides per sample. We observed good quantitative reproducibility between each of the 5 biological replicates per condition with a median Pearson correlation of 0.94 and a median coefficient of variation of 25.3%. The phosphoproteome of each WT kinase strain was compared with that of its kinase-dead mutant strain to determine significantly up- and down-regulated phosphosites. In total, we found 4,082 upregulated pY sites mapping to 1,970 proteins and 9,014 up- and down-regulated pS/T sites mapping to 2,361 proteins. Phosphorylation motifs of the WT kinases generally recapitulate motifs established in the literature and thus validate the kinase activities **(Figure S4)**. Phosphorylation profiles were highly variable between kinases **(Figure 1b and 1c)** but overall were consistent with the kinase activity assays in **Figure S1** and **Figure S2**. Highly active kinases such as vSRC and EPHB1 generated over 2,000 phosphosites while others were indistinguishable from the dead mutant, suggesting that they failed to become active when expressed in yeast (Corwin *et al*, 2017; Jehle *et al*, 2022). The different kinase groups (full-length pY, kinase domain pYd, vSRC mutants, and pST) can generally be distinguished using dimensionality reduction methods such as tSNE and kinases with a similar number of targets (e.g. WT vSRC and EPHB1) showed distinct phosphorylation profiles **(Figure 1d)**. The majority of induced pY sites have never been observed before, but there is also some up-regulation of endogenous pY sites with established functions **(Figure 1e)**, for example pY192 on the activation loop of the MAPK Slt2. Likewise, the regulated pS/pT sites that we observe upon kinase expression are distributed between those mapping to the ‘core’ pS/pT phosphoproteome and those that are condition-specific (Leutert *et al*, 2023) (**Figure 1e**). Fold-changes between Y kinases and their dead mutants are generally higher for Y sites than S/T sites, confirming the expectation that direct substrates are more strongly regulated than indirect downstream targets **(Figure 1e)** (Kanshin *et al*, 2017).

### Structural profile of the spurious pY phosphoproteome

The recent availability of AlphaFold2 enables the structural analysis of mutations and PTMs on a proteome-wide scale (Jumper *et al*, 2021; Varadi *et al*, 2022). We take advantage of these models to perform a structural analysis across all of our pY protein substrates (n=1970 proteins), with the goal of understanding how spurious phosphorylation may perturb the proteome.

We first calculate the relative solvent accessibility (RSA) for all pY sites directly from the AF2 structural models (**Figure 2a**). This analysis reveals 35.09% of the pY sites to be buried (**Figure 2b**), which is similar to the fraction of buried residues (35.86%) we calculate for a set of spurious pY sites identified previously (Corwin *et al*, 2017) **(Fig S5a-5b)**. We then predict the order/disorder content of our pY sites, using the AF2 structural models and the mean RSA in a +/- 12 amino acid window surrounding the pY position as a proxy for protein disorder (Akdel *et al*, 2022). We find that the average accessibility in this window tends to be low and therefore predict the majority of our pY sites (83.3%) to map to ordered regions (**Figure 2c**). Similar conclusions are found (73.3% ordered pY) when using sequence-based predictors of protein disorder **(Fig S5c-d)**, in agreement with the analysis of spurious pY order/disorder content performed by Corwin et al (Corwin *et al*, 2017). To give context to our findings, we conducted this analysis on a recent reference pS/pT phosphoproteome in yeast (Leutert *et al*, 2023), revealing that our spurious pY sites are significantly more likely to be buried and ordered than endogenous pS/pT sites (Kolomogorov-Smirnov p < 2 x 10^-16^, Fisher p < 2×10^-16^, **Figure 2b-c)**. Interestingly, endogeneous pY sites lie between the extremes of spurious pY and endogenous pS/pT in terms of their accessibility and order (see section below ‘Limited overlap between the spurious and native phosphoproteomes’).

**Figure 2.**
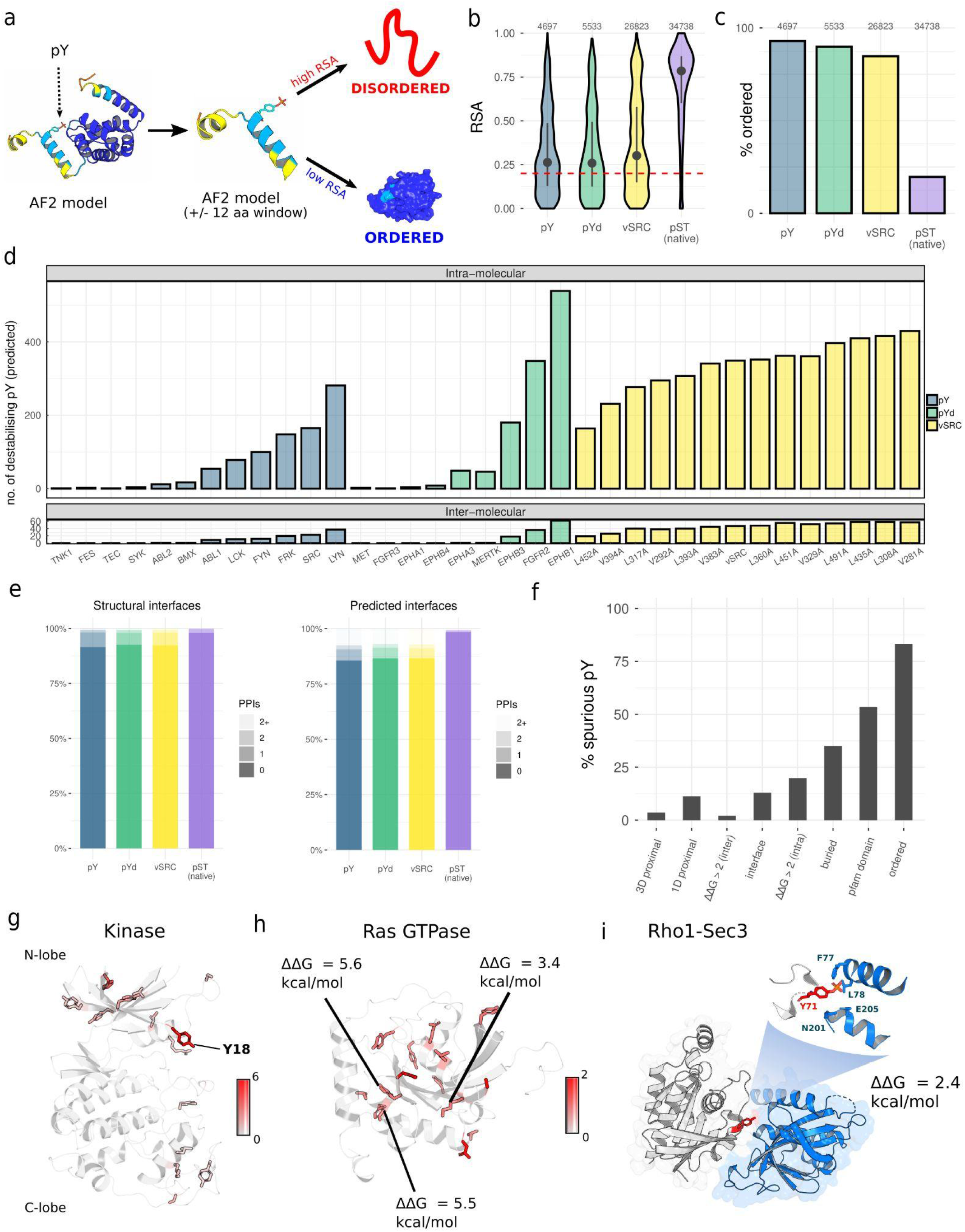
Structural overview of spurious phosphorylation across the proteome. **a)** Use of AlphaFold2 structural models to calculate the relative solvent accessibility (RSA) and disorder of spurious phosphosites. **b)** The relative solvent accessibility (RSA) of spurious phosphosites generated by full-length tyrosine kinases (pY), tyrosine kinase domains (pYd), WT vSRC and vSRC mutants (vSRC), and the endogenous yeast pS/pT sites reported in (Leutert *et al*, 2023) (pST). The red dashed line corresponds to the cut-off for buried residues set at an RSA of 0.2 **c)** The percentage of phosphosites that map to ordered regions for the pY, pYd, vSRC, and pST groups. **d)** For each of the tested kinases, the number of spurious pY sites (WT-dead) predicted to destabilise the protein fold (intramolecular) or at least one protein-protein interface (intermolecular), on the basis of a ΔΔG threshold of 2 kcal/mol. **e)** For the pY, pYd, vSRC, and pST groups, the number of unique interfaces (per phosphosite) found in structural models (PDB, homology, or AF2) (left), or predicted from machine learning (right) (Meyer *et al*, 2018). **f)** Structural profile of all unique pY phosphosites detected in this study. **g)** The total number of unique pY sites that map to the protein kinase domain, represented by the AF2 model of the cyclin-dependent kinase Pho85. **h)** The total number of unique pY sites that map to the Ras GTPase domain (including predicted destabilising pY), represented by the AF2 model of Gsp1. **i)** The spurious phosphosite Rho1 pY71 is predicted to destabilise the Rho1-Sec3 interaction with a ΔΔG of 2.4 kcal/mol (PDB: 3a58).

The presence of so many spurious pY sites mapping to buried and structured regions suggests that a substantial fraction of the spurious phosphoproteome may destabilise protein folding and thus have functional consequences. We check this systematically by using FoldX to predict the change in free energy of folding (ΔΔG for Y -> pY) across all unique pY sites mapping to an AF2 structural model. We validate this approach by comparing ΔΔG predicted on a sample of AF2 models with ΔΔG from the corresponding experimental structures (n=231), revealing a strong correlation of 0.91 **(Fig S5e)**. We also find that, as expected, phosphorylation becomes more destabilising as pY sites become less accessible (**Fig S5f**) and that around 20% of all spurious pY are predicted to be destabilising using a standard ΔΔG threshold of 2 kcal/mol (Nishi *et al*, 2011; Wagih *et al*, 2018; Høie *et al*, 2022). This results in a large number of predicted destabilising pY across the proteome for highly active kinases such as vSRC (n=349) and EPHB1 (n=539), with an average number of 178 destabilising pY sites for all non-redundant kinases that we consider to be active (**Figure 2d**). Spurious phosphorylation therefore has the potential for pervasive deleterious effects on protein structure.

We next map the spurious pY sites to protein-protein interfaces and again use FoldX to predict phosphosites that destabilise protein interactions. We use Interactome3D for the structural annotation of protein-protein interfaces in *S. cerevisiae* (Mosca *et al,* 2013), in addition to a recent AF2-based screen that was performed proteome-wide for the prediction of protein complexes and their structures (Humphreys *et al*, 2021). We supplement this data with high-confidence machine learning predictions from InteractomeInsider (Meyer *et al*, 2018), which predicts residues at the protein-protein interface (**Figure S5g**). Only a small fraction of pY sites (7.2%) map to at least one structural interface although this percentage increases to 13.0% when also considering interfaces predicted by machine learning **(Figure 2e)**. The total number of interfaces (per pY) predicted by machine learning is generally much higher than what is observed from the structural models **(Figure S5h-i)**, indicating missing structural data in the *S. cerevisiae* proteome. However, a significant number of interactions are still predicted to be destabilised (ΔΔG > 2 kcal/mol) on the basis of these protein structures and particularly for highly active kinases such as vSRC (n=47) and EPHB1 (n=62) (**Figure 2d**). All structural parameters are given in **Table S3** (pY), **Table S4** (pYd), **Table S5** (vSRC), and **Table S6** (pS/pT).

Spurious pYs may also impact function by directly affecting endogenous signalling on protein surfaces. To assess this, we map spurious pY phosphosites in relation to native pS/pT sites in *S. cerevisiae*. We first search simply for pY sites proximal to endogenous pS/pT sites in 1D space, using a +/-4 amino acid window. We find 441 such spurious pY sites in our data, which are 11.2% of the number of total unique sites. We next repeat the analysis in 3-dimensions, given that two phosphosites may be separated in 1D space but not in a fully-folded protein structure. We use AF2 models and a distance threshold of 8 Angstroms for this purpose (Humphreys *et al*, 2021). We find a significant number of such proximal 3D sites although they are a small fraction of the spurious pY phosphoproteome (3.6%). Notably, the average pST-pST (native-native) distances from the global pS/pT phosphoproteome in *S. cerevisiae* are significantly lower than corresponding pY-pS/T (spurious-native) distances **(Fig S5j)**, providing orthogonal support for the recent finding that native pS/pT sites tend to cluster in 3D space (Bludau *et al*, 2022).

In **Figure 2f**, we present a structural profile of spurious pY phosphorylation covering protein order/disorder, destabilisation (folding and interactions), and spurious-native proximity (1D and 3D). These data can be examined further to find examples of functionally significant proteins that may be perturbed by phosphorylation. For example, we find 27 unique spurious pY that map to the kinase domain of native *S. cerevisiae* kinases, reflecting the intrinsic specificity of kinases for other kinases as substrates (Invergo & Beltrao, 2018). Of these, 20 map to the smaller N-terminal lobe that is important for ATP binding and the regulation of kinase function (Pellicena & Kuriyan, 2006; Taylor & Kornev, 2011; McClendon *et al*, 2014), including a position in the glycine-rich loop that is frequently phosphorylated here and known to have an inhibitory effect in some kinases when natively phosphorylated (Steinberg, 2018) **(Figure 2g)**. The Ras GTPase family (including Ras/Rab/Ran/Ypt1 sub-families) is also critical for signal transduction and is spuriously phosphorylated at 19 unique sites in our data. These pY sites are more equally distributed in the 3D structure, but we predict 6 to destabilise the protein fold (Ras2 Y115, YPT32 Y101, YPT52 Y194, Rho5 Y216, Rho5 Y177, GSP1 Y148), and 4 (Rho1 Y71, YPT1 Y109, Gsp1 Y149, Gsp1 Y157) to map to interfaces covering 13 protein-protein interactions of known or predicted structure **(Figure 2h)**. In 9 cases the spurious pY is predicted to destabilise the physical interaction; in **Figure 2i** we show as an example the destabilisation of the Rho1:Sec3 interface (PDB: 3a58, ΔΔG = +2.433 kcal/mol) required for targeting of the secretory protein Sec3 to the plasma membrane (Yamashita *et al*, 2010). Finally, we give an example of spurious-native phosphorylation proximity in Tma19, a protein that stabilises microtubules and has apoptosis-related functions (Rinnerthaler *et al*, 2006). The spurious phosphosite pY18 is in close proximity (5.4 Angstroms) to a native phosphosite (pS15) (Holt *et al*, 2009) that is known to be functional because it gives a deleterious growth phenotype in methotrexate and ethanol conditions when mutated to alanine (S15A) **(Figure S5k)** (Viéitez *et al*, 2022). In cases like this, the spurious phosphorylation may be either affecting the recognition of the native phosphosite or functionally mimicking it, as has been suggested for rapidly evolving protein regions (Holt *et al*, 2009).

In summary, our proteome-wide structural analysis of spurious pY predicts the widespread destabilisation of proteins and their PPIs.

### Effects of kinase expression on fitness

We test the relationship between the activity of the kinases and their impact on fitness by measuring the growth of all strains expressing these kinases. The WT kinases are compared with their corresponding kinase-dead mutants to specifically determine the fitness effect of pY phosphorylation and not of heterologous protein expression. In total, the fitness effects of 44 kinases (13 Y kinases, 10 Y kinase domains, 7 S/T kinases, and 14 vSRC variants) were tested across 41 conditions known to induce various stresses **(Table S7)** and used previously in (Dionne *et al*, 2021).

We take the size of yeast colonies over time as a proxy for fitness and use an automated plate reader to measure changes in the colony size across 50 different time points (16 replicates per measurement). The area under the growth curve (AUC) was calculated for the WT kinase and inactive mutant, and then the difference in AUC between the two (WT-dead) was used to determine the fitness cost of pY phosphorylation for that kinase **(Figure 3a)**. A summary of the results are given in **Figure 3b-c**, **Figure S6,** and **Table S8.** We find that 5/13 Y kinases, 3/10 Y kinases domains, 3/7 S/T kinases, and 14/14 vSRC variants are significantly more deleterious (FDR-adjusted) for fitness in at least one condition when comparing the WT kinase with the corresponding kinase-dead mutant. The kinase-dead strains have generally similar colony sizes **(Figure S7)**, demonstrating a comparable impact upon fitness even if some were poorly expressed in the Western blots **(Figure S1 and S2)**. Direct competition assays were performed for the pY and vSRC strains against an empty landing-pad control and used to determine the relative growth of the kinase **(Figure 3d, Table S9, see methods),** giving results that correlate significantly with the AUC_min_ values derived from the colony sizes (r_s_=0.70, p=9.85×10^-5^) and thus validate these assays.

**Figure 3.**
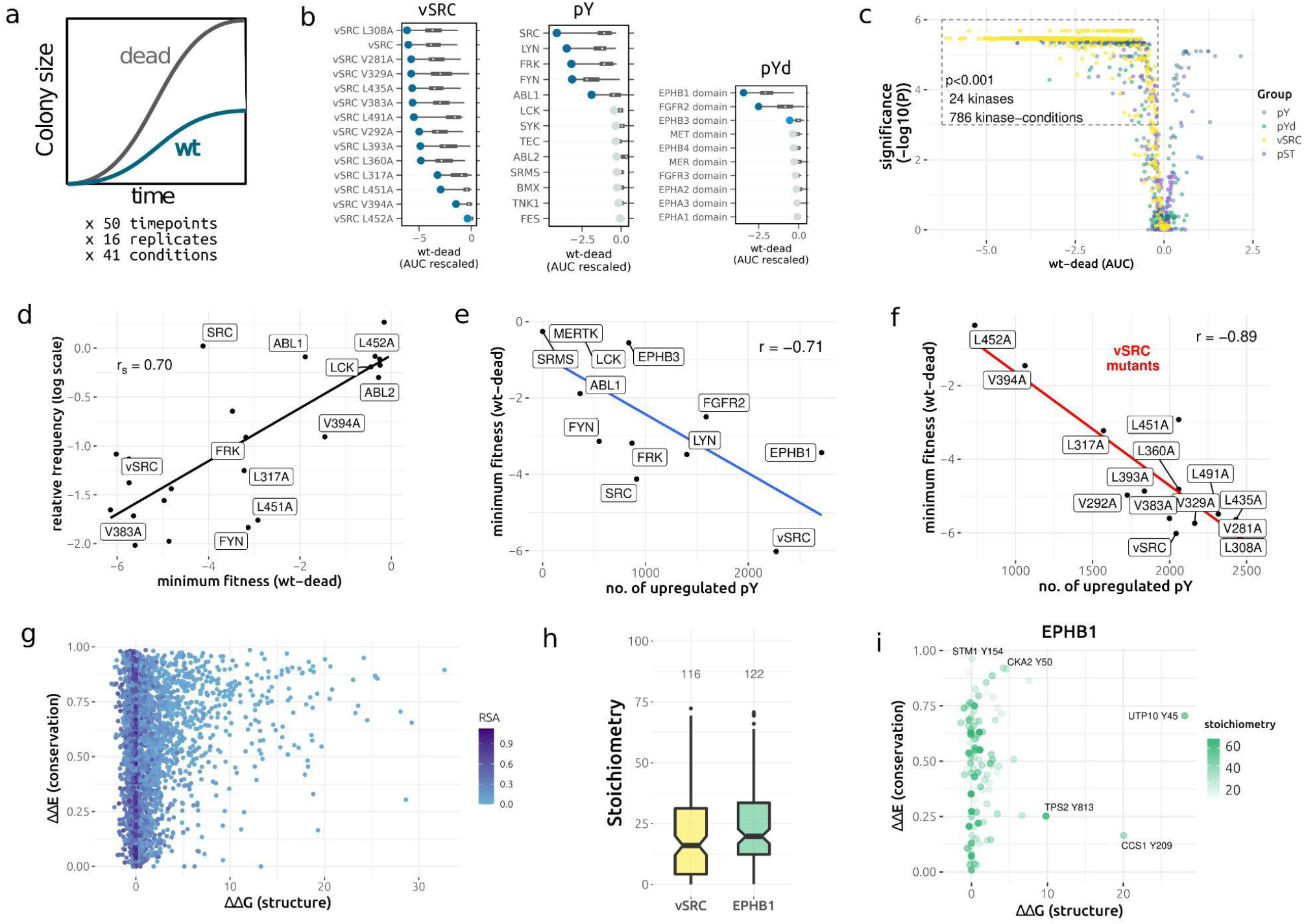
Measuring the fitness effect of human kinase expression (WT-dead) in yeast. **a)** Colony size is used as a proxy for fitness and the difference in area under the growth curve (AUC) between the WT kinase and the kinase-dead mutant is used to infer the fitness effect of spurious phosphorylation. **b)** The fitness for each kinase tested, represented by the WT-dead fitness defect across 41 conditions. More negative AUC values (WT-dead) indicate greater toxicity. The colours of the dots indicate the statistical significance of the fitness score: dark blue indicates P<0.001, light blue indicates P<0.05, and grey indicates non-significance **c)** volcano plot of WT-dead AUC (x-axis) against the FDR-adjusted p value for the significance of the difference between the WT and kinase-dead mutant. pY: full-length tyrosine kinase, pYd: tyrosine kinase domain, vSRC: vSRC and mutants, pST: serine/threonine kinases. **d)** correlation between WT-dead fitness inferred from colony sizes (x-axis), and relative growth between the kinase and empty landing-pad control (y-axis). The relative frequencies were log-scaled and taken at the final time point (72 hours) of the competition assay (see Methods). Strains that grow worse than the control have negative relative frequencies. **e)** correlation between the number of spurious pY (per kinase) and the minimum fitness (WT-dead) across conditions for all non-redundant kinases tested here **f)** correlation between the number of spurious pY (per kinase) and the minimum fitness (WT-dead) across conditions for WT vSRC and the vSRC mutants tested here **g)** scatter plot for the predicted effect of spurious pY on protein structure (ΔΔG) and predicted effect on the basis of sequence conservation (ΔΔE). More destabilising pY have higher ΔΔGs and and more conserved Y positions have values closer to 1. All unique spurious pY are included. **h)** distribution of stoichiometries for vSRC pY substrates (n=116) and EPHB1 pY substrates (n=122), where significant phosphosite regulation was inferred (q < 0.01, see Methods). **i)** For EPHB1, the predicted effect of spurious pY on protein structure (ΔΔG) and predicted effect on the basis of sequence conservation (ΔΔE) for all sites with inferred stoichiometries. Higher inferred stoichiometries are given in dark green and lower inferred stoichiometries are given in light green.

We find that the toxicity of spurious phosphorylation can be sensitive to the conditions of growth **(Figure S8a)**, as shown in (Jehle *et al*, 2022), as some treatments (e.g. phosphatase inhibitor cocktail, rapamycin 50 nM, and SDS 0.02%) more strongly impact fitness than the least harmful condition (Caffeine 10mM) (p=1.1×10^-2^, 4.3×10^-4^, 4.7×10^-3,^ respectively, Mann-Whitney one-sided). We also find that different conditions are highly correlated in terms of their fitness effect across kinases **(Figure S8b)** and likewise that kinases with similar activity levels correlate strongly across conditions **(Figure S8c)**, suggesting few kinase-condition interactions overall. However, strong interactions can be observed for a small number of kinase-condition pairs **(Figure S8d)**. For example, the vSRC mutation L491A has a minimum WT-dead fitness in the SDS 0.02% condition that is 3.5 times lower than the median fitness effect across all conditions **(Figure S8d)**. Differential fitness effects between conditions **(Figure S8e)** could be explained by loss-of-function (LOF) phosphorylation effects mapping to conditionally-sensitive genes (Bosch-Guiteras & van Leeuwen, 2022). However, cross-referencing the data here with a systematic conditional KO screen in *S. cerevisiae (*Viéitez *et al,* 2022) provides little support for this hypothesis **(Figure S8f)**. The mechanism underlying the conditional fitness defects therefore remains unclear.

We next relate the phosphoproteomic data presented above **(Figure 1)** with the fitness data for each kinase. We find that the minimum fitness defect per kinase correlates strongly with the number of upregulated pY sites for that kinase, both across kinases (**Figure 3e)** and between the vSRC mutants (**Figure 3f)**. Similar results were found when using the median fitness defect across conditions (**Figure S9a and Figure S9b**).

It is unclear whether this strong correlation represents a general phosphorylation burden distributed across many sites or, at the other extreme, a very small number of deleterious pY that increase in stoichiometry as the kinase activity also increases. To support the first hypothesis we try to determine if the fitness-phosphorylation correlation can be improved by considering only those pY that we predicted to destabilise the protein fold **(Figure 2)**. However, the resulting correlation (r =-0.64) is lower than that calculated from all pY sites (r =-0.71). This approach is limited by the possibility that a phosphosite can be deleterious for protein function in a way that is not easily predictable from the protein structure (Gerasimavicius *et al*, 2022; Høie *et al*, 2022). To consider this possibility, we perform variant effect prediction (VEP) for all upregulated pY sites (∼4000 sites on ∼2000 proteins) by retrieving substrate homologs across the protein universe and predicting the phosphorylation effect on the basis of sequence conservation **(Figure 3g, Figure S10)**. The effect of phosphorylation cannot be inferred directly but we derive it here by calculating the effect of tyrosine substitution for all 20 AAs and then taking a weighted average for pY on the basis of its biophysical similarity to the 20 proteinogenic amino acids (**Figure S9c, Methods**). We combine this data with the structural ΔΔG calculations above to predict any pY with a deleterious effect, either on the basis of structure or sequence conservation. However, incorporating this data on predicted deleterious effects offers no improvement to the fitness-phosphorylation correlation across a range of structural and conservation thresholds (**Figure S9d)**. Conservation (ΔΔE) and structural parameters (ΔΔG) for all pY phosphosites are presented in **Figure 3g, Figure S10,** and listed in **Table S10** alongside their surface accessibilities.

Assuming a general loss-of-function (LOF) paradigm for toxicity would also suggest that focusing on essential proteins would offer more explanatory power for fitness than considering spurious substrates proteome-wide, which would include substrates with no measurable phenotype upon knock-out (KO). However, even in combination with the structural and conservation parameters outlined above, no significant improvement in correlation was found for the subset of essential substrates (**Figure S9e)**. Next, we attempted a simple weighting of pY phosphosites on the basis of the number of predicted interfaces – to account for spurious protein-protein interactions – and found some strengthening of the phosphosite-fitness correlation for the unique Y kinases (−0.761 vs. −0.71) but not for the vSRC mutant (−0.873 vs. −0.89) (**Figure S9f & S9g**). Finally, we consider that the Y kinase domain EPHB3 has ∼800 upregulated pY sites, of which 191 are predicted to be destabilising and 125 are predicted to be deleterious on the basis of sequence conservation **(Figure S10, Figure S11).** However, expression of this kinase has no significant fitness effect in most conditions **(Table S8, Figure S9a)** and only a small effect for the condition of minimum fitness **(Figure 3e).** The results taken together suggest that a significant fraction of spurious pY have a negligible fitness effect in spite of the *in silico* predictions.

One possibility is that many of the pY detected by mass spectrometry are either transient, of low stoichiometry, and/or rapidly removed by phosphatases during protein synthesis and folding (Hunter, 2009). We probe this point further by performing a general measurement of phosphorylation stoichiometry across substrates. As we use a regulation-based approach to infer stoichiometry (see Methods), we choose the two most strongly active kinases in our dataset: EPHB1 and vSRC **(Figure S10, Figure S11**). The relatively high abundance of phosphorylated peptides in these conditions ensures high regulation between wt and kinase-dead, which results in more accurate stoichiometry estimations (**Figure S12, Figure S13**). For the phosphosites that are sufficiently regulated between the WT and dead conditions, we infer a median stoichiometry of 16.0% for vSRC and 19.8% for EPHB1 **(Figure 3h)**, suggesting that the effect of phosphorylation would be much weaker than if the equivalent positions were mutated in the genome (i.e at 100% effective stoichiometry). Indeed, haploinsufficiency screens in yeast suggest that even the removal of one gene copy (∼50% protein reduction) does not have a measurable fitness effect for the vast majority of genes (Deutschbauer *et al*, 2005). While some predicted deleterious pY are still found at intermediate-high stoichiometries and are highlighted in **Figure 3i** (EPHB1) and **Figure S14** (vSRC), we do not generally observe a decrease in abundance for the substrates of such sites (**Figure S15, Figure S16)**. Stoichiometry values are given in **Table S11.**

### Limited overlap between the spurious and native phosphoproteomes

All kinases used in this study (except vSRC) have native substrates in the human proteome that act as functional effectors of the kinase. For spurious phosphorylation, it is an open question whether the non-native kinase will phosphorylate preferentially the homologs of its native substrates or indiscriminately across the proteome. We address this question first by mapping all yeast-human orthologs where possible for the identified spurious pY substrates. For preferential targeting, we would expect spurious pY substrates to have more human orthologs than expected by chance. We find that, of 1347 spurious pY substrates, 781 (58.0%) have at least one human homolog. The corresponding percentage for detected proteins not spuriously phosphorylated is 47.9%, indicating that spurious substrates are more likely to have human orthologs than random *S. cerevisiae* proteins (**Figure 4a**, p=9.8×10^-9^, odds ratio = 1.5, Fisher test). Conversely, native pY substrates are not significantly more likely to have at least one human ortholog (**Figure S17a**, 49.0%, p=0.86, odds ratio = 1.04, Fisher test).

**Figure 4.**
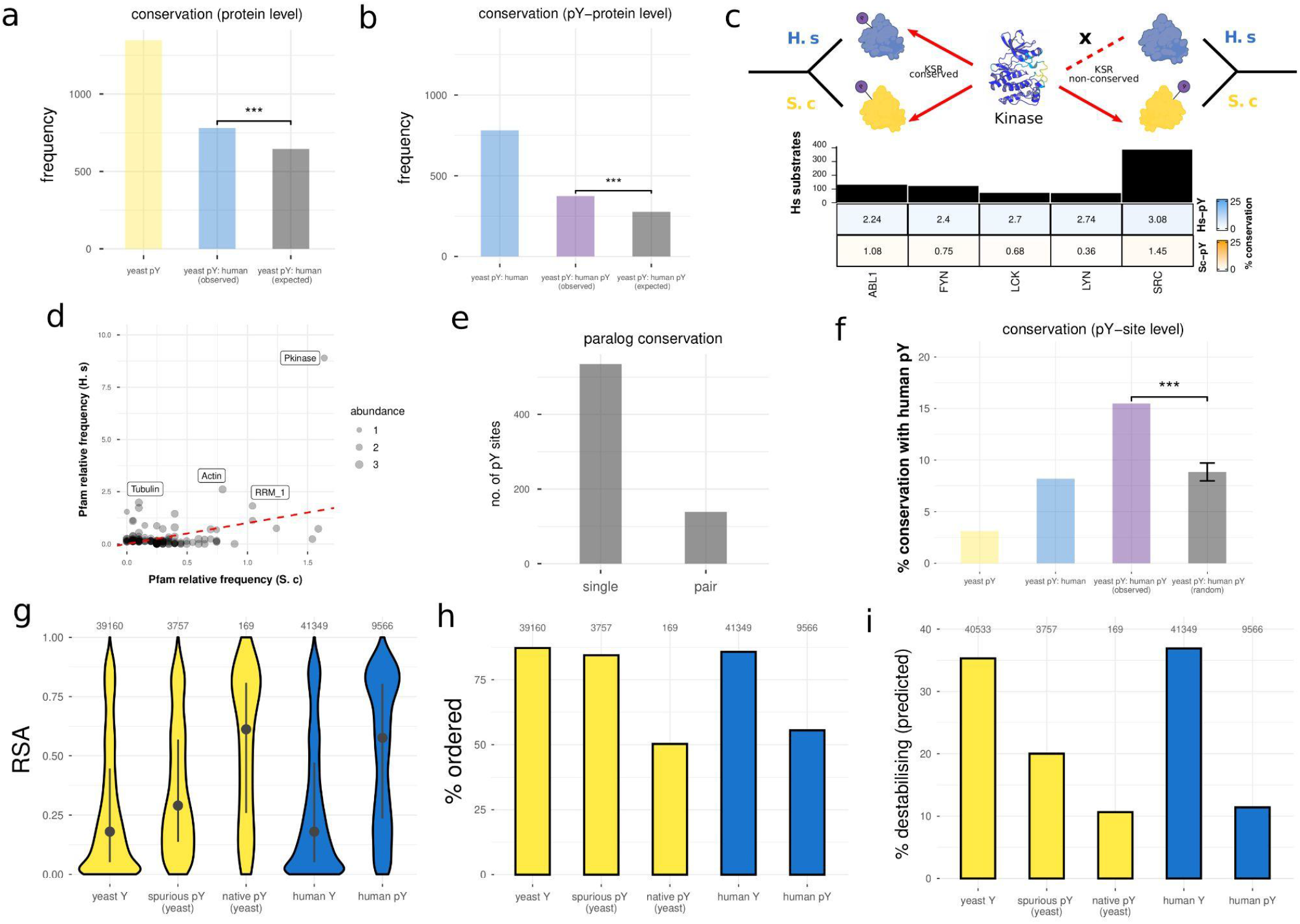
Conservation between spurious pY in yeast and native pY in human. **a)** Human-yeast conservation at the whole protein level. ‘Yeast pY’ – number of unique spurious pY proteins in yeast, excluding any native pY proteins. ‘Yeast pY: human (observed)’ – number of *observed* unique spurious pY proteins in yeast with at least one ortholog in human. ‘Yeast pY: human (expected)’ – number of *expected* unique spurious pY proteins in yeast with at least one ortholog in human **b)** Human-yeast conservation at the level of whole protein pY phosphorylation. ‘Yeast pY: human’ – number of *observed* unique spurious pY proteins in yeast with at least one ortholog in human. ‘Yeast pY: human pY (observed)’ – number of *observed* unique spurious pY proteins in yeast with at least one ortholog in human that is Y-phosphorylated. ‘Yeast pY: human pY (expected)’ – number of *expected* unique spurious pY proteins in yeast with at least one ortholog in human that is Y-phosphorylated. **c)** conservation of kinase-substrate relationships (KSRs) between human and yeast. Top row (blue) is the proportion conserved relative to the sample size of known KSRs in humans. Bottom row (yellow) is the proportion conserved relative to the sample size of KSRs found in this study for yeast. **d)** Relative frequency (summing to 1) of Pfam domain phosphorylation in yeast (x-axis) and human (y-axis). Abundance is given in the units of log10(parts per million). Scatter plot includes only domains supported by at least 5 unique pY sites in either human or yeast. **e)** conservation of spurious pY phosphorylation between paralog pairs for paralog-specific peptides (see Methods). Single: phosphorylated on one paralog-specific peptide but not the homologous position. Pair: phosphorylated on both paralog-specific peptides at homologous Y positions. **f)** site-specific (i.e. alignment-based) conservation between spurious pY and human native pY. As a percentage of all unique yeast spurious pY (yeast pY, yellow), all unique yeast spurious pY with at least one human ortholog (yeast pY: human, blue), all unique yeast spurious pY with at least one pY-phosphorylated human ortholog (yeast pY: human pY (observed), purple), and all unique yeast spurious pY with at least one pY-phosphorylated human ortholog with x100 randomisations of the human pY positions. **g-i)** For yeast non-pY Y residues, spurious pY residues, native pY residues, human non-pY Y residues, and human pY residues, distribution of surface accessibility (RSA), percentage mapping to predicted ordered regions, and the percentage predicted to be destabilising for the protein fold (using a threshold of ΔΔG > 2 kcal/mol).

We next test for the conservation of pY phosphorylation between human (native) and yeast (spurious) at the level of the whole protein. Considering yeast proteins that have a human ortholog, we found pY sites in the human protein for 47.9% of the spurious pY yeast proteins compared to 35.4% for the non-pY yeast proteins. This indicates that conservation of the pY phosphorylation state between yeast and human is greater than the null expectation (**Figure 4b**, p=1.3×10^-7^, odds ratio = 1.68, Fisher test). For endogenous pY substrates in yeast this enrichment is even greater given that in 76.4% of cases at least one human ortholog is tyrosine phosphorylated (**Figure S17b**, 1.3×10^-11^, odds ratio = 5.90, Fisher test). Therefore, while endogenous pY substrates are no more likely than random proteins to have a human ortholog, if one exists then it is much more likely to be tyrosine-phosphorylated in human than for random *S. cerevisiae* proteins or spurious pY substrates.

For cases where the pY state is conserved between human and yeast orthologs, we use known kinase substrate relationships to determine if the sites are phosphorylated by the same kinase in both species. The completeness of KSR annotations is limiting for this analysis (Invergo & Beltrao, 2018; Needham *et al*, 2019), as only five of the active tyrosine kinases used here have more than 20 known human substrates. Examination of this data reveals that the extent of conservation between human and yeast KSRs is very low across all kinases, either when considering the conserved KSRs as a fraction of the total number of yeast (spurious) substrates or of human (native) substrates **(Figure 4c)**. This analysis can be broadened by examining the conservation of protein domain preference among the pY sites in human and yeast **(Figure 4d)**. This approach is agnostic to known kinase-substrate annotations, orthology mappings, and can include anciently diverged homologs that share a protein domain. This analysis reveals conserved phosphorylation on a small number of domains including the kinase, actin, and tubulin domains. However, 32% of all spuriously phosphorylated domains in yeast are not observed to be tyrosine-phosphorylated in humans, revealing the recurrent spurious phosphorylation of non-native domains. Finally, we show that when paralog pairs in yeast are spuriously phosphorylated, the phosphosite position is conserved in 20.6% of cases (**Figure 4e**, see Methods), which is higher than the corresponding conservation (∼13%) found for native pS/pT sites in yeast (Leutert *et al*, 2023).

While the phosphorylation state can be conserved at the level of the whole protein or protein domain, as described, the effect of phosphorylation on protein function may differ depending on the position of the phosphosite in the protein sequence. We therefore check the extent to which yeast pY (spurious) and human pY (native) align at the site-specific level. The overall level of site-specific conservation across all spurious pY is very low (3.10%). As expected, the percentage conservation increases (to 8.20%) when considering only spurious pY with at least one human ortholog, and more so to 15.49% when considering only spurious pY with at least one human ortholog that is also phosphorylated in human (i.e. pY conservation at the protein level) **(Figure 4f)**. Through 100 random permutations of the phosphosite positions we show that this percentage is significantly higher than the chance expectation **(Figure 4f)**. The observed extent of site-specific conservation is modestly improved when considering proximal pY in an alignment window (-/+0, -/+3, -/+5, -/+ 7, -/+9 alignment positions), which may reflect a slight under-estimation of site-conservation caused by alignment errors, phosphosite mislocalisation, or selectively neutral shifts in phosphosite position (Landry *et al*, 2014) **(Figure S17c)**.

For the structural analysis of spurious phosphorylation (above, **Figure 2**), we described the tendency of non-native pY sites to map to buried and ordered protein regions. We now repeat this analysis for native human pY sites to determine if shifts in phosphosite structural preference can explain the low level of pY conservation at the site-specific level. For context, we also perform this analysis for yeast non-phosphorylated Y, human non-phosphorylated Y, and endogenous pY in yeast. The results reveal that, compared to yeast spurious pY, human native pY is significantly more likely to be accessible **(Figure 4g)** and disordered **(Figure 4h)**, and therefore less likely to destabilise the protein structure, confirmed using ΔΔG predictions *in silico* (**Figure 4i**). In turn, yeast spurious pY is more accessible, more disordered, and less destabilising than the expectation for non-phosphorylated Ys in the proteome, given that we predict *∼*20% of our spurious pY to be destabilising compared to a rate of *∼*35% for Ys not modified according to mass spectrometry. The data therefore demonstrate significant structural differences between the spurious and native phosphoproteomes in yeast and human, and also between spurious phosphorylation and the null expectation for random (non-modified) Ys, revealing some structural constraint upon spurious phosphorylation **(Figure 4g-i, ‘yeast Y’)**. However, order-disorder discordance of pYs between human-yeast ortholog pairs poorly explains the absence of site conservation as pY sites that do not align have consistent order/disorder predictions 78.9% of the time **(Figure S17d)**.

Taken together, this analysis reveals that spurious phosphorylation is weakly biased towards homologs of native kinases substrates but there is also significant phosphorylation of completely new (i.e. non-homologous) substrates. Spurious-native pY conservation is low at the level of the whole proteins and even more so at the site-specific level, although it is still higher than the expectation for completely random Y phosphorylation.

### Tyrosine counter-selection in metazoan proteomes

Tyrosine kinases are absent from fungal species and the small level of detectable Y phosphorylation on fungal proteins likely arises from dual-specificity kinases (Lim & Pawson, 2010; Corwin *et al*, 2017; Leutert *et al*, 2023). It has been proposed that the emergence of *bona fide* Y kinases in animals resulted in the selective loss of Y content in their proteomes to avoid spurious phosphorylation (Tan *et al*, 2009, 2011), a claim that has been challenged by later studies (Su *et al*, 2011; Pandya *et al*, 2015; Kritzer *et al*, 2018). Having at hand a set of sites that could be phosphorylated if such kinases were present in fungi, we can examine further this model. We explicitly test this hypothesis at the level of the proteome **(Figure 5a, 5b, Figure S18)**, proteins **(Figure 5c, 5d, Figure S19)**, linear motifs **(Figure S20)**, and individual sites **(Figure 5e-g, Figure S21)** using the spurious pY data (i.e. *∼*4,000 unique upregulated pY sites total for all expressed kinases) generated in this study.

**Figure 5.**
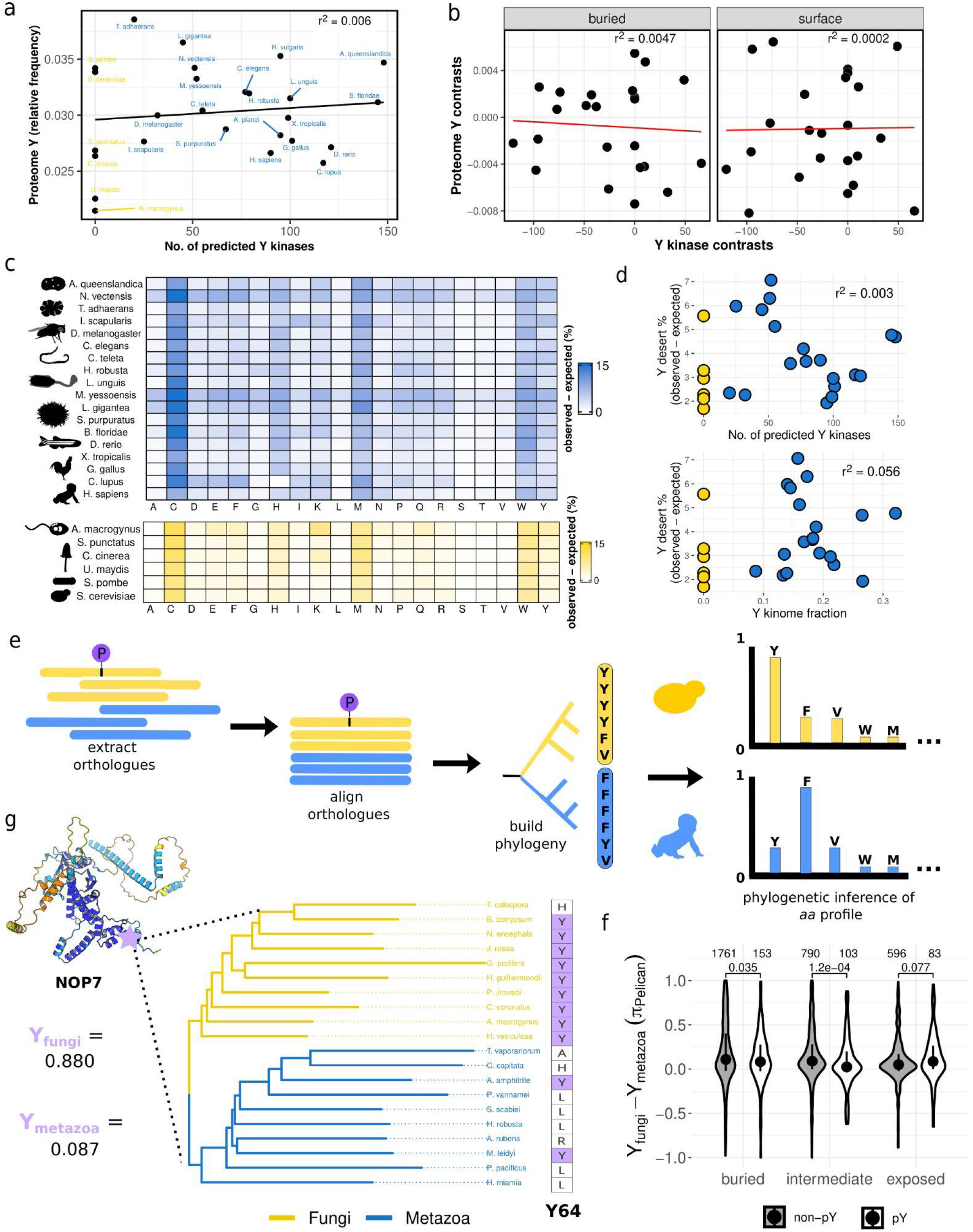
Testing for counter-selection against spurious pY residues in animal species. **a)** Correlation between the number of predicted tyrosine kinases and proteomic Y content (relative frequency) for 20 animal species (blue) and 6 fungal species (yellow). **b)** Correlation between the number of predicted tyrosine kinases and proteomic Y content (relative frequency) after applying a phylogenetic correction (phylogenetically independent contrasts, (Felsenstein, 1985)). The left panel represents buried tyrosine (RSA < 0.2) proteome-wide and the right panel surface residues (RSA > 0.4) proteome-wide. The species and kinases analysed are the same as in panel **a**. **c)** Percentage of amino acid deserts (observed - expected) for all 20 canonical amino acids. An amino acid desert is defined as a protein where more than 50% of the protein length is missing the amino acid. The percentage observed is compared to the percentage calculated for 100 simulated proteomes (see *Methods*). Animal species in blue, fungal species in yellow. **d)** Top: correlation between the number of predicted tyrosine kinases (per species) and percentage of tyrosine deserts (observed - expected). Bottom: correlation between the number of predicted tyrosine kinases as a fraction of the total number of kinases (per species) and percentage of tyrosine deserts (observed - expected). **e)** A schematic for testing Y counter-selection at the level of individual sites in multiple sequence alignments (MSAs). For each unique spurious pY, orthologs are extracted, the ortholog sequences are aligned, a phylogenetic tree is constructed, and a profile of amino acid preference is inferred on the basis of the MSA and phylogenetic tree. Animal species in blue, fungal species in yellow. **f)** Difference between the inferred preference for Y in fungal species and metazoa species, calculated as their difference in equilibrium frequencies (***π***) by the Pelican software (Duchemin *et al*, 2023). The results are given for pY and non-pY tyrosines and separated according to their solvent accessibility (buried: RSA > 0.2, intermediate: 0.2 < RSA < 0.4, exposed: RSA > 0.4). Results only given for sites with significant changes in amino acid profile (between animals and fungi) at an adjusted p-value < 0.05. Distributions (pY and non-pY) compared using a Kolmogorov-Smirnov two-tailed test. **g)** Example of a protein (NOP7) with predicted counter-selection against Y (Y64) in animal species. Inferred Y preferences (0-1) are represented by Y_fungi_and Y_metazoa_. A small sample of representative species are shown for the fungal and metazoan clades.

We first demonstrate that we are able to recapitulate the previously observed negative correlation between the number of predicted tyrosine kinases and proteomic Y content (r^2^ = 0.69 compared to r^2^ = 0.66 in (Tan *et al*, 2009), **Figure S18a)**, using available reference proteomes and a state-of-the-art method for kinase prediction and classification (see *Methods*). We then take advantage of the increased availability of genome sequences in recent years to repeat this analysis across a larger set of species that is more phylogenetically balanced and representative of major phyla across the metazoa and fungi (**Figure S18b**). Using this new data, we find no significant relationship between the number of predicted tyrosine kinases and proteome Y content (r^2^ = 0.006, **Figure 5a**). Notably, the Y content in other fungal species (*C. cinerea, U. maydis*, *S. punctatus*, and *A. macrogynus*) is significantly lower than in *S. cerevisiae* and *S. pombe* even though all lack Y kinases. We also generated a species tree to calculate the evolutionary distance between species (**Figure S18b**), which allows this analysis to be repeated while accounting for the branch lengths connecting data points. The original correlation remains statistically significant after controlling for phylogenetic non-independence (r^2^ = 0.52, p=0.002, **Figure S18c**), in agreement with (Tan *et al*, 2011), while we still find no significant relationship between the number of tyrosine kinases and proteome Y content for our more recent set of species after applying this control (r^2^ = 0.036, p=0.376, **Figure S18d)**. The analysis was repeated after stratifying the data according to ‘buried/accessible’ status of the pY sites using proteome-wide AF2 models (across all species) but again without a significant correlation (buried: r^2^ = 0.0047; surface: r^2^ = 0.0002; **Figure 5b).** Finally, we test the expectation that orthologs of the spurious substrates identified here will be subject to stronger selection against spurious phosphorylation than non-substrates, assuming some conservation of structure, abundance, and subcellular localisation between animal and fungal species. While we indeed observe a stronger correlation for pY orthologs (r= −0.25,p=0.24, **Figure S18e left-top**) compared to the rest of the proteome (r= −0.19,p=0.38, **Figure S18e left-bottom**), larger differences are found for other amino acids such as aspartate (r_ortho_ = −0.297, r_non-ortho_ = −0.015, **Figure S18e-middle, Table S12**). Taken together, these results fail to strongly support the hypothesis of proteome-wide pY counter-selection in animal species.

Evidence for pY counter-selection may instead be present at the level of individual proteins. The presence of PTM ‘deserts’ has been suggested as a mechanism for the avoidance of off-target modifications (Fredrickson *et al*, 2013; Boomsma *et al*, 2016; Sharma *et al*, 2017), prompting recent studies into lysine-depleted regions and their relation to spurious lysine ubiquitination and degradation by the proteasome (Kampmeyer *et al*, 2023; Szulc *et al*, 2023). In line with this reasoning, we investigate the possibility of analogous ‘tyrosine deserts’ to avert tyrosine phosphorylation. At the same time we examine deserts for the other 19 amino acids. Differences in the proteomic amino acid composition and disorder content across species bias the expected number of amino acid deserts under selective neutrality. We therefore randomly simulate each proteome 100 times, while accounting for the amino acid and disorder content of each species, and then compare the expected number of amino acid deserts with those observed from real sequences **(Figure 5c)**. We find on average a 3.29% increase in tyrosine deserts beyond the neutral expectation, behind lysine (3.41%), histidine (4.66%), tryptophan (6.87%), methionine (7.07%), and cysteine (10.65%) **(Table S13)**. However, tyrosine desert frequency does not correlate with the absolute or relative number of tyrosine kinases in the kinome **(Figure 5d)**. A gene ontology (GO) analysis of human tyrosine deserts reveals a strong enrichment of tyrosine deserts in proteins that regulate signalling receptors and in proteins with cytokine activity **(Figure S19)**. Therefore, while we do not observe proteome-wide tyrosine depletion that correlates with the expansion of tyrosine kinases, tyrosine depletion may be linked to other proteins involved in tyrosine signalling (O’Shea *et al*, 2011; Morris *et al*, 2018).

Additional tests for pY counter-selection can be performed at the level of sequence motifs by checking for amino acid strings that are either depleted or completely absent in the proteome or a subset of interest (Koulouras & Frith, 2021; Georgakopoulos-Soares *et al*, 2021). We search for avoided motifs in three subsets of the human proteome: human proteins not tyrosine phosphorylated, human proteins not tyrosine phosphorylated that are orthologs of spurious pY substrates in *S. cerevisiae*, and all proteins with the GO:0005125 (cytokine activity) and GO:0030545 (signalling receptor regulator activity) terms that are enriched among tyrosine deserts in human. The results of this analysis are given in **Figure S20.** Non-phosphorylated human proteins are significantly more likely than random proteins to have avoided Y motifs (12.3% vs. 11.2%, p < 0.01), but not orthologs of spurious pY substrates (p=0.79), or the GO term proteins (10.1% vs. 5.0%, p=0.054).

Finally, in the absence of support for tyrosine counter-selection at the level of the whole proteome, selection against tyrosine residues may still be evident at the level of individual phosphosites that were significantly upregulated between the WT and dead conditions. The depletion of tyrosine at any given site is itself not evidence of selection against spurious phosphorylation because the presence or absence of Y could affect the stability of protein folding or interactions (Pace *et al*, 2001; Koide & Sidhu, 2009). However, the preferential or biased counter-selection for *S. cerevisiae* sites detected as Y-phosphorylated here – compared to non-phosphorylated tyrosines and while controlling for surface accessibility – would support this hypothesis. We test this explicitly using an evolutionary model that takes as an input a multiple sequence alignment (MSA) and phylogenetic tree of spuriously phosphorylated substrates and their orthologs across animal and fungal species. This software (*Pelican, (Duchemin et al, 2023)*) aims to detect shifts in amino acid preferences between lineages by modelling each clade using a separate equilibrium frequency vector of amino acids **(Figure 5e)**. Using this approach, we determine whether an equilibrium depletion of tyrosine in animal species offers a significantly better fit to the data than using a uniform equilibrium vector across all species (animal and fungal).

From this point we refer to shifts in amino acid equilibrium frequencies inferred from the phylogeny and MSA as shifts in the amino acid ‘preference’. At a false discovery rate (FDR) of 0.05, around *∼*35% of all Y positions among the spurious substrates exhibit a significant change in amino acid profile between animal and fungal sequences **(Figure S21a).** We then test specifically for shifts in Y preference by comparing the Y preference in animals with the Y preference in fungal species (Y***_π_***_-fungi_ - Y***_π-metazoa_***). Values close to 1 correspond to cases where Y is strongly preferred in the fungi but not in the metazoa, and *vice versa* for negative values. We also stratify the data according to accessibility (buried, intermediate, exposed) given established relationships between residue exposure and the rate of evolution (Echave *et al*, 2016; Bricout *et al*, 2023), and that the molecular impact of phosphorylation will likely depend upon the accessibility of the phosphosite **(Figure S5f).** This analysis does not reveal any significant evidence for Y counter-selection for pYs relative to non-pYs (**Figure 5f, Table S14)**. It is possible that some of the ‘non-pY’ tyrosines in this analysis are susceptible to phosphorylation but were simply not targeted among any of the active Y kinases used to generate the data. We therefore repeat this analysis but only using non-pY sites with a low motif score across all human Y kinases for which we have specificity models (Sugiyama *et al*, 2019). Applying this control again does not support the hypothesis of pY counter-selection **(Figure S21b)**. Another possibility is that there is strong counter-selection against pY motif residues flanking the phosphoacceptor to prevent Y phosphorylation (Deng *et al*, 2014; Li *et al*, 2023; Sugiyama *et al*, 2019). However, we do not observe strong evidence for motif counter-selection **(Figure S21c, Figure S21d).** Finally, we check the possibility that ‘high quality’ substrates – those that are phosphorylated by many independent kinases – will be subject to stronger counter-selection than sites targeted by a small number of Y kinases. Recurrently phosphorylated Y are more strongly counter-selected than rarely phosphorylated Y on average but the difference is not significant (p=0.084, Kolmogorov-Smirnov, one-sided) **(Figure S21e)**.

Taken together, our findings do not support the hypothesis of proteome-wide pY counter selection or that this can serve as a driver of proteomic Y content. However, we do not discount the possibility of pY-counter selection for restricted phylogenetic lineages or a smaller number of sites and proteins functionally related to signalling **(Figure S19, Figure S20).** For example, Y64 of the yeast protein Nop7, which is an essential protein that is required for the synthesis of some ribosomal subunits (Adams *et al*, 2002) **(Figure 5g).** The functional consequences of Y absence/presence in such candidates requires further experimental research.

## Discussion

Here we have expressed human tyrosine kinases in *S. cerevisiae* and then measured the impact of kinase expression both on yeast fitness and the yeast phosphoproteome, each time comparing the WT kinase to its kinase-dead (KD) mutant. Proteome-wide structural analysis of these sites reveals that around 20% are predicted to be destabilising for protein folding **(Figure 2f)**. Additionally, we perform proteome-wide variant effect prediction (VEP) on the basis of phosphoacceptor (Y) conservation and predict >1000 spurious pY sites to negatively affect protein function **(Figure S10, Figure 3g)**. Paradoxically, while we observe a strong negative correlation between the number of spurious pY sites and fitness **(Figure 3e-f),** it is also apparent that many such sites predicted to be deleterious have a negligible effect on fitness. For example, expression of the kinase EPHB3 produces 274 predicted deleterious pY sites but only has a small fitness defect in one of the tested growth conditions **(Figure 3e, Figure S9-S11).**

A potential explanation for the surprisingly weak fitness defects is that pY may be of low stoichiometry. Contrary to the *in silico* methods used for VEP that assume 100% stoichiometry for DNA-encoded mutations, phosphorylation of tyrosines occurs post-translationally and is reversed by phosphatases, which actually exist in yeast (Pincus *et al*, 2008; Hunter, 2009; Lim & Pawson, 2010; Chen *et al*, 2017). In agreement, we found that a phosphatase inhibitor cocktail was one of the most harmful growth conditions among the 41 treatments we tested **(Figure S8a)**. As additional evidence, we found low estimated stoichiometries among the strongly active kinases v-SRC and EPHB1 **(Figure 3h),** which is also consistent with the generally low stoichiometry of pY phosphorylation in human tissues in the absence of specific stimulation (Sharma *et al*, 2014; Tsai *et al*, 2015, 2022). We further note that yeast lacks conventional SH2 and PTB domains for the binding of phosphotyrosine (Kaneko *et al*, 2012), which can shield the pY residue from phosphatase activity (Jadwin *et al*, 2018; Hunter, 2009). Finally, we consider that a haploinsufficiency screen in yeast, with a theoretical reduction in protein abundance of 50%, generated a measurable fitness defect for only ∼3% of the yeast genome (Deutschbauer *et al*, 2005). In principle, many proteins can therefore individually experience a reduction in abundance without a strong impact on fitness.

Our data also allows for a comparison between the spurious pY sites generated here and the native human pY proteome. Crucially, the former represents sites that have never been subject to evolutionary constraints while the latter have been exposed to natural selection. We observe that, compared to human pY sites, spurious sites are less accessible, more likely to map to ordered regions, and more likely to destabilise the protein structure **(Figure 4g-i).** This implies a structural shift towards more benign phosphorylation in systems with evolved phosphotyrosine signalling. However, it is not clear whether this represents the action of purifying selection against destabilising pY directly or the requirement for many pY sites to be accessible to SH2 domains, (PTB) domains, and tyrosine phosphatases for reversible modification. How such buried sites (∼35% spurious pY) came to be phosphorylated in the first place is another important consideration. Buried phosphorylation has been documented in a small number of cases, indicating the transient exposure of the phosphoacceptor during protein dynamics (Li *et al*, 2008; Henriques & Lindorff-Larsen, 2020; Orioli *et al*, 2022; Swadling *et al*, 2022); a related explanation is that binding of the substrate *in vivo* to other molecules promotes a different conformational state to the one predicted by AlphaFold2 as the default structure (Mayer, 2015; Del Alamo *et al*, 2022). Another possibility is that phosphorylation occurs co-translationally before the protein is fully synthesised and folded. However, it is not yet clear if these mechanisms can sufficiently explain the ∼1400 buried pY generated in this study.

Further examination of the pY sites generated here also reveals the recurrent spurious phosphorylation of targets across large swathes of the proteome. This implies that, in the context of off-target signalling, counter-selection against adventitious binding will not be restricted to native targets of the effector protein **(Figure 4)**. Such contexts may arise in nature due to: the sudden shift in protein expression levels (Faulkner *et al*, 2009; Young *et al*, 2015), the sudden shift in protein localisation (Nguyen Ba *et al*, 2014), the horizontal transfer of a PTM to a new host (Cummings *et al*, 2022), and/or the generation of a fusion protein leading to the aberrant specificity and activation of the effector (e.g. BCR-Abl kinase) (Smolnig *et al*, 2023). This result in principle is consistent with the proposal in 2009 that the animals and their unicellular relatives underwent a proteome-wide reduction in Y to avoid off-target phosphorylation by the emergent tyrosine kinases (Tan *et al*, 2009, 2011). However, we find no strong support for the hypothesis after testing it at the level of the proteome, proteins, motifs, and individual sites. Our work is therefore in closer alignment with other studies that have challenged this hypothesis (Su *et al*, 2011; Pandya *et al*, 2015; Kritzer *et al*, 2018). In particular, the 2018 finding that v-SRC toxicity can be suppressed by the overexpression of a single S/T kinase (SMK1) in spite of persistent tyrosine phosphorylation in the proteome (Kritzer *et al*, 2018). Proteome-wide selection against off-target Y phosphorylation therefore seems unlikely, although we do not exclude the possibility of Y counter-selection for a smaller subset of spurious targets or phylogenetic lineages.

In conclusion, even in spite of clear toxicity, this work suggests that many spurious interactions (i.e. phosphosites) have a minimal impact upon fitness. Consistent with this, we do not observe any excess counter-selection of such spurious sites relative to sites not at risk of promiscuous binding. The findings indicate off-target protein interactions can be tolerated to a certain extent during evolution, possibly providing a means by which functional systems may evolve from non-functional ancestors. It is a remaining challenge to understand the conditions under which toxicity may arise and the fraction of spurious interactions that can give rise to fitness defects.

## Supplementary figures

**Supplementary figure 1:**
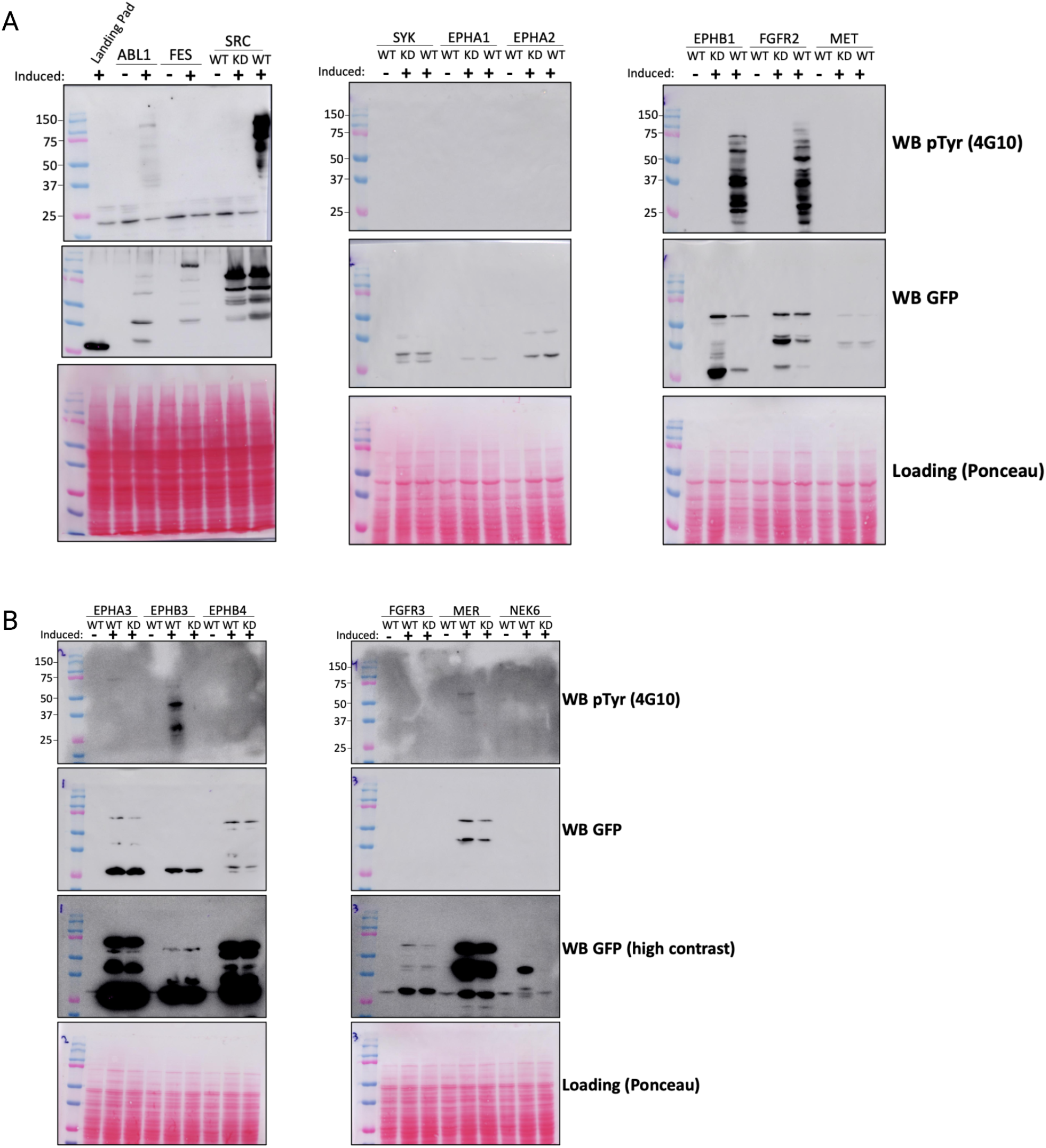
p-Tyr activity of expressed kinase in yeast. Yeast cells were grown with (+) or without (-) induction to check for p-Tyr activity. Western blots to detect p-Tyr (pTyr (4G10)) and GFP (kinase expression) were done. **a)** ABL1, FES, SRC, SYK, EPHA1, EPHA2, EPHB1, FGFR2 and MET. In the left-most panel, a small sample of kinases (ABL1, FES, and SRC) are compared to an empty landing pad (first lane) as a control. **b)** EPHA3, EPHB3, EPHB4, FGFR3, MER and NEK6. WT: wild-type kinase. KD: kinase-dead mutant.

**Supplementary figure 2:**
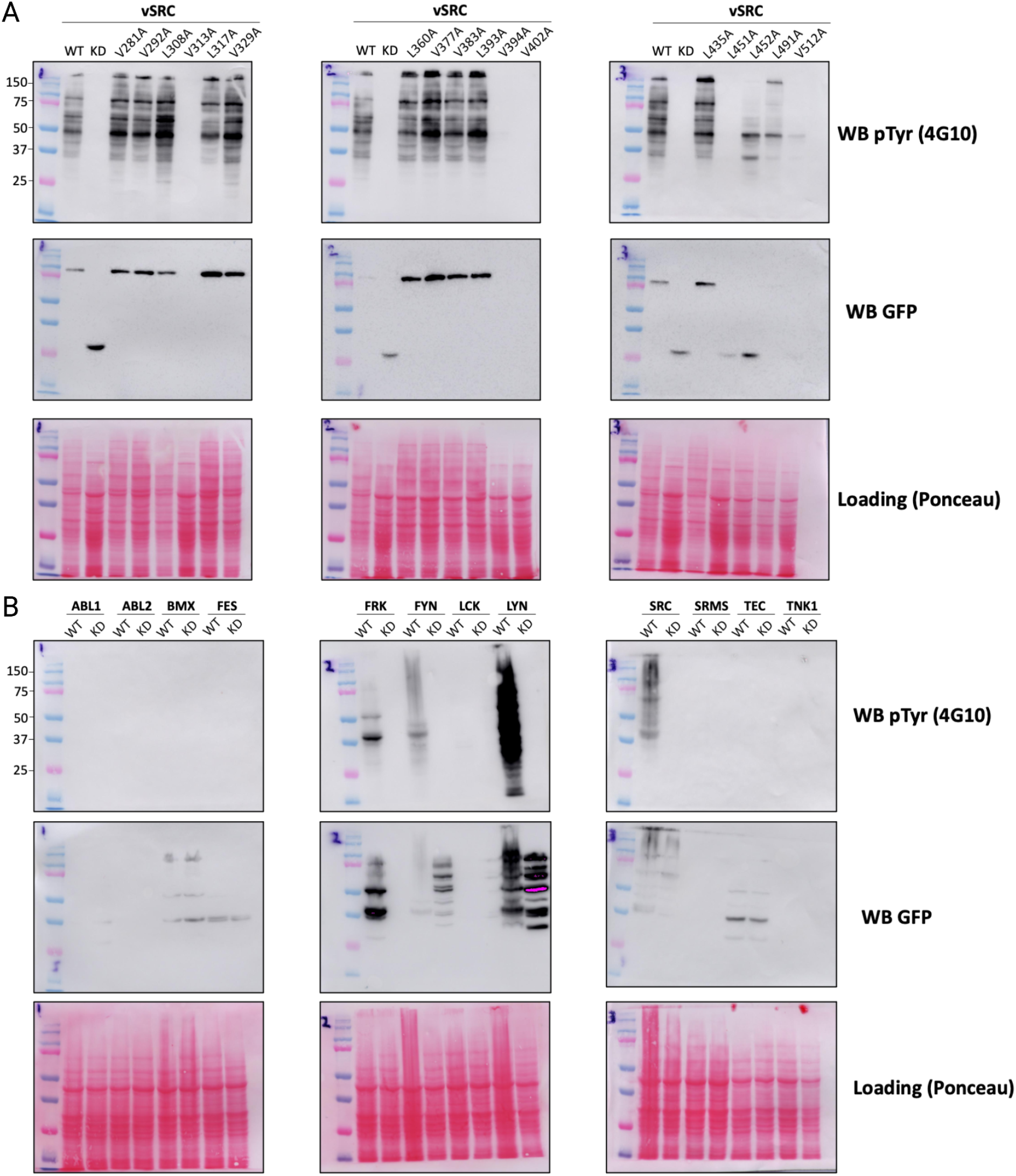
p-Tyr activity of expressed kinase in yeast. Yeast cells were grown with induction to check for p-Tyr activity. Western blots to detect p-Tyr (pTyr (4G10)) and GFP (kinase expression) were done. **a)** vSRC and its mutants **b)** ABL1, ABL2, BMX, FES, FRK, FYN, LCK, LYN, SRC, SRMS, TEC and TNK1. WT: wild-type kinase. KD: kinase-dead mutant. The vSRC mutants V313A, V402A, V512A, and V377A were later found to contain non-synonymous mutations in the coding sequence (via sequencing) and so were excluded from any further analysis.

**Supplementary Figure 3:**
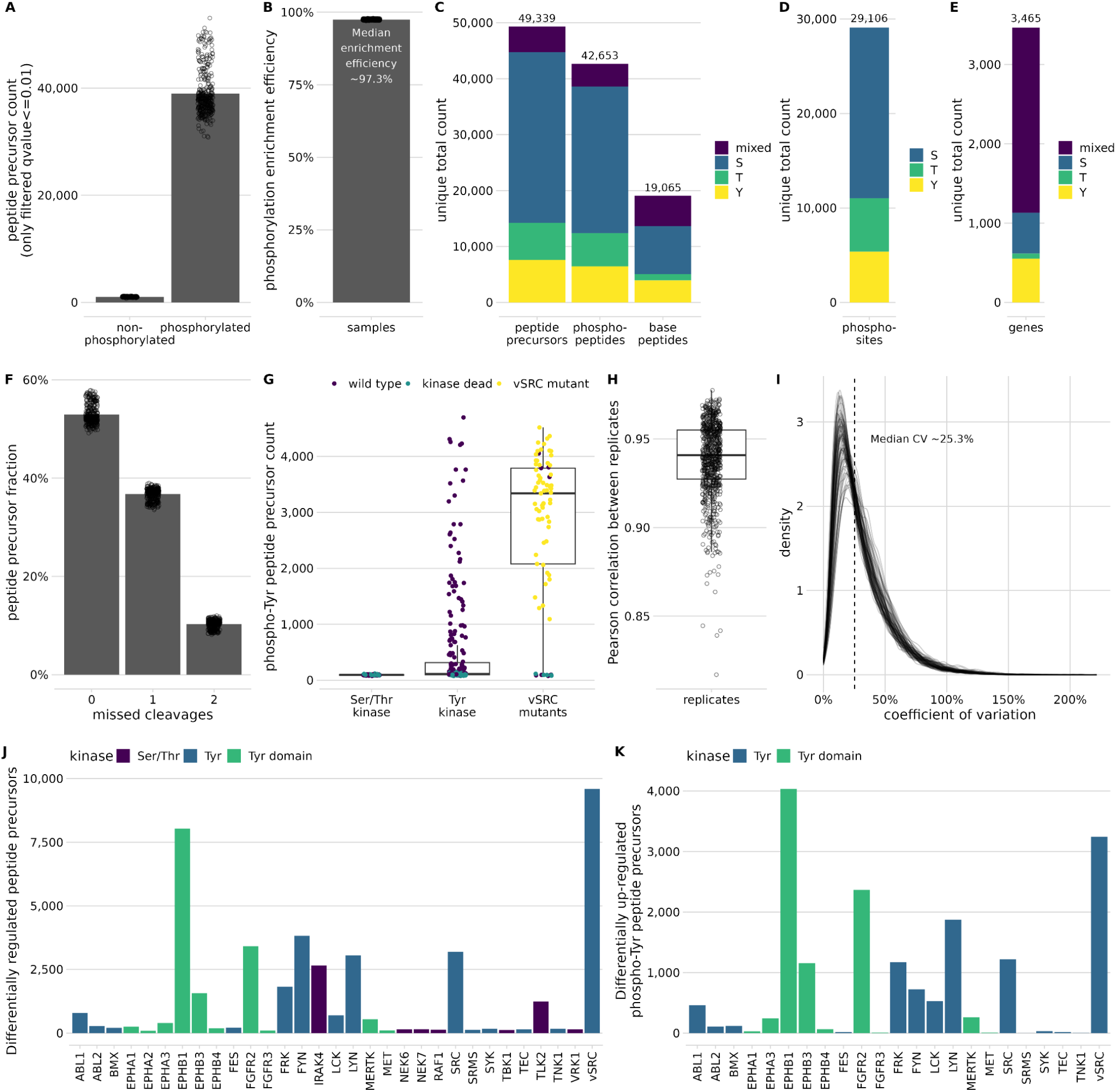
Phospho-proteomics quality control. Phospho-enriched samples were measured by mass spectrometry and processed by FragPipe/DIA-NN. **a-b)** The vast majority of identified peptide precursors are phosphorylated, yielding a median enrichment efficiency of 97.3%. For these two plots, the data was only filtered using local q-value ≤ 0.01. **c-e)** Total unique identifications on peptide precursor (PEPT(ph)IDEK_2), phospho-peptide (PEPT(ph)IDEK), base peptide (PEPTIDEK), phospho-site (Ser(ph)-123 on gene 1) and gene level. **f)** Digestion efficiency as shown by the number of missed cleavages per peptide precursor. **g)** Phospho-tyrosine containing peptide precursors are identified almost exclusively in tyrosine kinase wild type and vSRC wild type / mutant conditions. **h)** Pearson correlation is shown between biological replicates of the same conditions. **i)** Density graph showing the coefficients of variation for the 5 biological replicates per condition. **j-k)** SAM statistical significance testing shows **j)** the up- and down-regulated peptide precursors for all serine/threonine and tyrosine kinases, as well as **k)** the up-regulated peptide precursors containing phospho-tyrosine for the tyrosine kinases.

**Supplementary figure 4:**
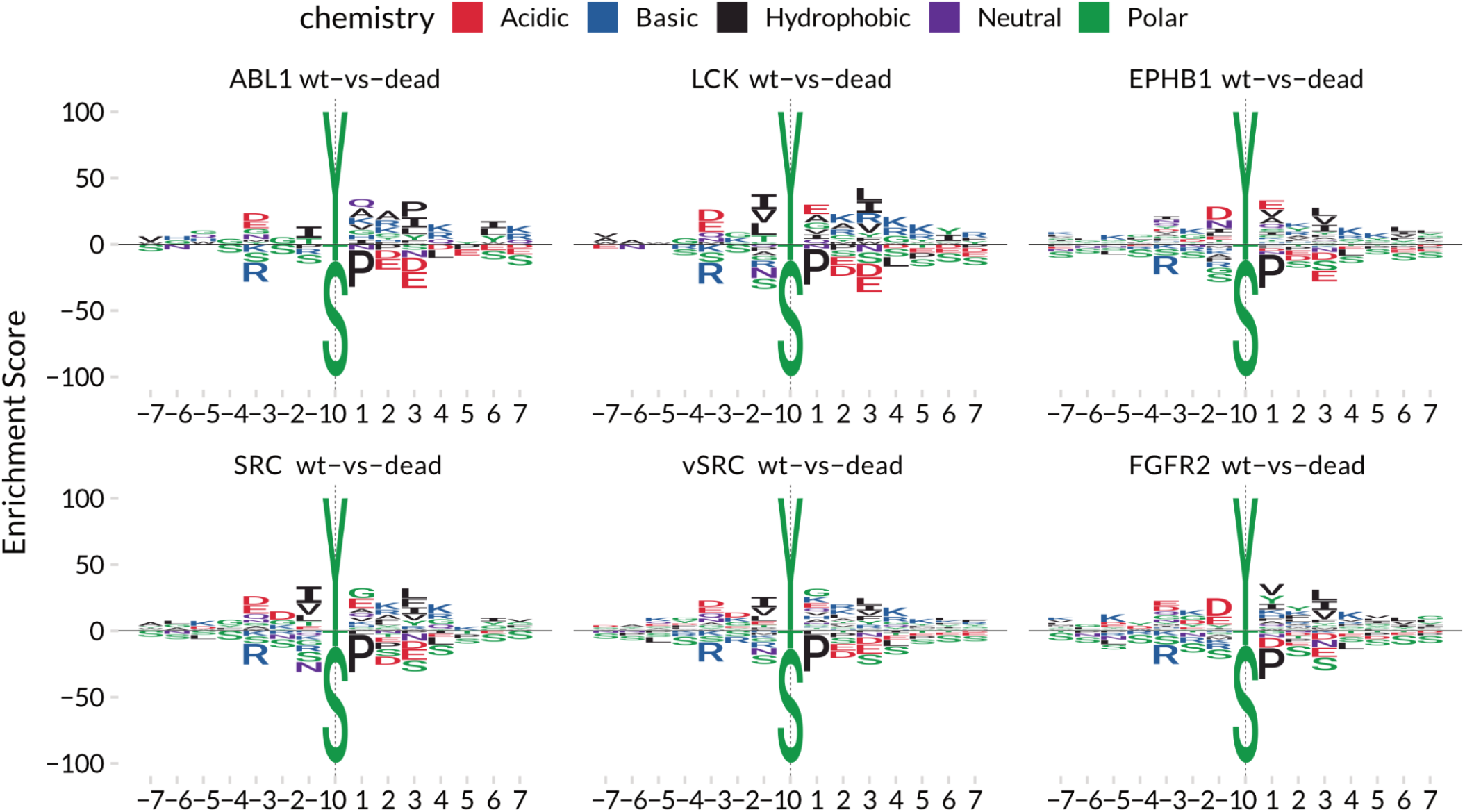
Kinase substrate motif enrichment analysis of selected phospho-tyrosine kinases. To test for kinase activity, kinase substrate motif enrichment analysis was performed on phospho-tyrosine containing peptides vs all other phospho-peptides using the R package dagLogo (Ou *et al*, 2020). By using only phosphorylated tyrosine peptides as the foreground, it could be assured that they must have been phosphorylated by the respective kinase. Amino acid positions were filtered for significance using a p-value threshold ≤ 0.05. The generated kinase substrate motifs recapitulate known kinase substrate preferences as for ABL1 (Colicelli, 2010), LCK and SRC (Shah *et al*, 2018).

**Supplementary figure 5:**
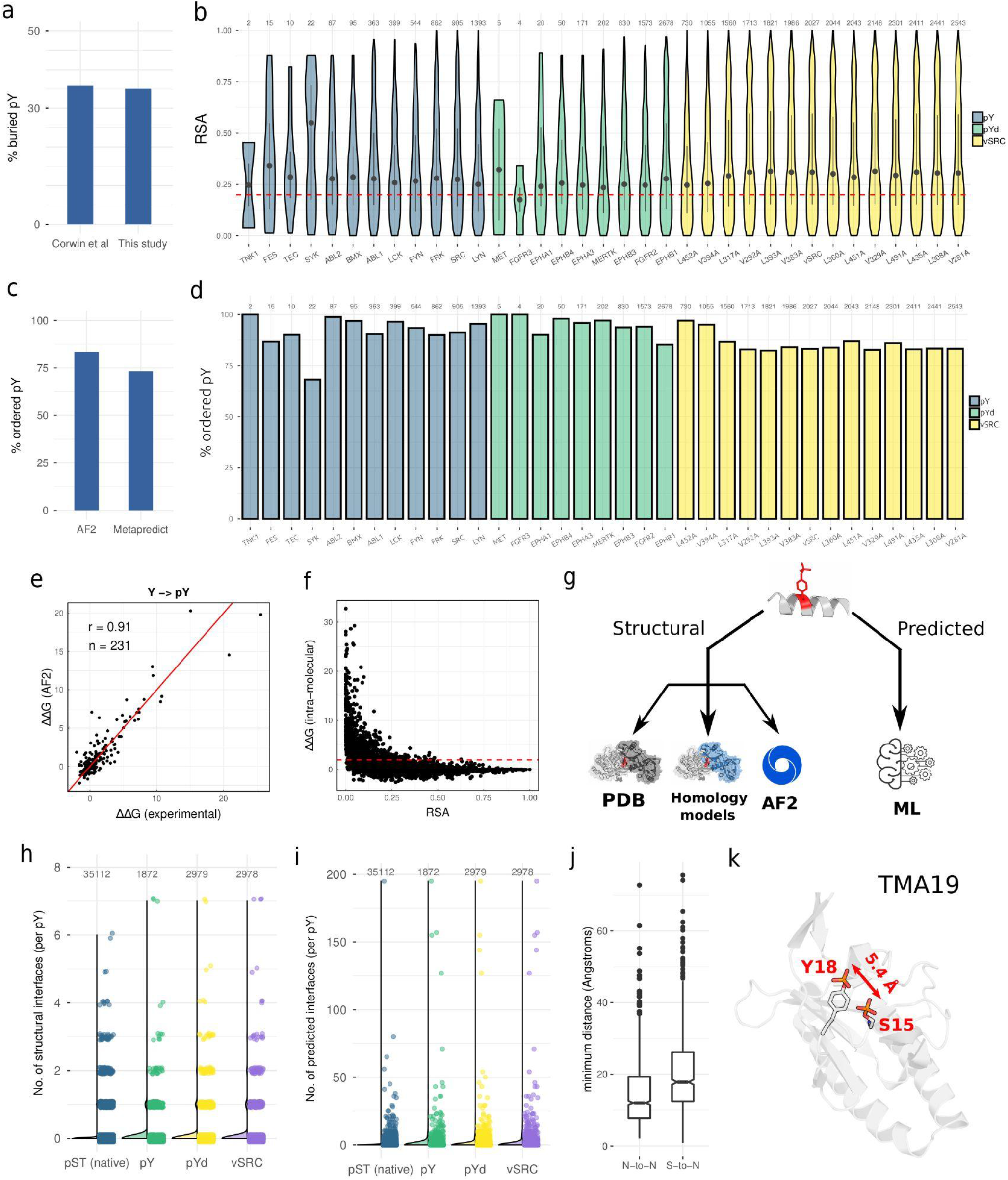
Structural overview of spurious phosphorylation across the proteome. **a)** For all unique spurious pY reported in (Corwin *et al*, 2017) and this study, the percentage of sites predicted to be buried from AF2 structures based upon a relative solvent accessibility (RSA) threshold of 0.2 **b)** The RSA of upregulated pY sites (WT-dead) for each kinase, divided by group. pY: full-length tyrosine kinases, pYd: tyrosine kinase domains, vSRC: WT vSRC and vSRC mutants. The red dashed line corresponds to the cut-off for buried residues set at an RSA of 0.2 **c)** For all unique spurious pY phosphosites reported in this study, the percentage mapping to predicted ordered regions based upon the AF2 structures (left) and the sequence-based predictor Metapredict (right) (Emenecker *et al*, 2021). **d)** The percentage of spurious pY sites mapping to ordered regions for each kinase, divided by group. Disorder/order predictions made upon the basis of the AF2 models (Akdel *et al*, 2022; Piovesan *et al*, 2022). **e)** Correlation of the FoldX ΔΔG (Y to pY) predictions between PDB structures and their corresponding AF2 models for spurious pY substrates. A resolution cut-off of 3Å was applied to the PDB models. **f)** For each unique spurious pY, relationship between the RSA and the predicted ΔΔG of phosphorylation (Y to pY). The red dashed line corresponds to a ΔΔG threshold of 2 kcal/mol for destabilising pY **g)** Spurious pY sites were mapped to structural interfaces corresponding to PDB structures (wwPDB consortium, 2019), homology models (Mosca *et al*, 2013), and AF2-based models of protein interactions (Humphreys *et al*, 2021). Interfaces of protein-protein interactions were also predicted via machine learning (Meyer *et al*, 2018). Panel created with the help of BioRender.com. **h)** The number of unique interfaces (per pY) found in structural models (PDB, homology, or AF2). **i)** The number of unique interfaces (per pY) predicted via machine learning (interactomeINSIDER) (Meyer *et al*, 2018). **j)** The minimum native-to-native (N-to-N) and spurious-to-native (S-to-N) 3D distances per protein. Native sites (pS/pT) were sourced from (Leutert *et al*, 2023). Spurious pY sites were detected in this study (Figure 1a). **k)** 3D distance between the native phosphosite pS15 and the spurious phosphosite pY18 in the AF2 model of TMA19.

**Supplementary figure 6:**
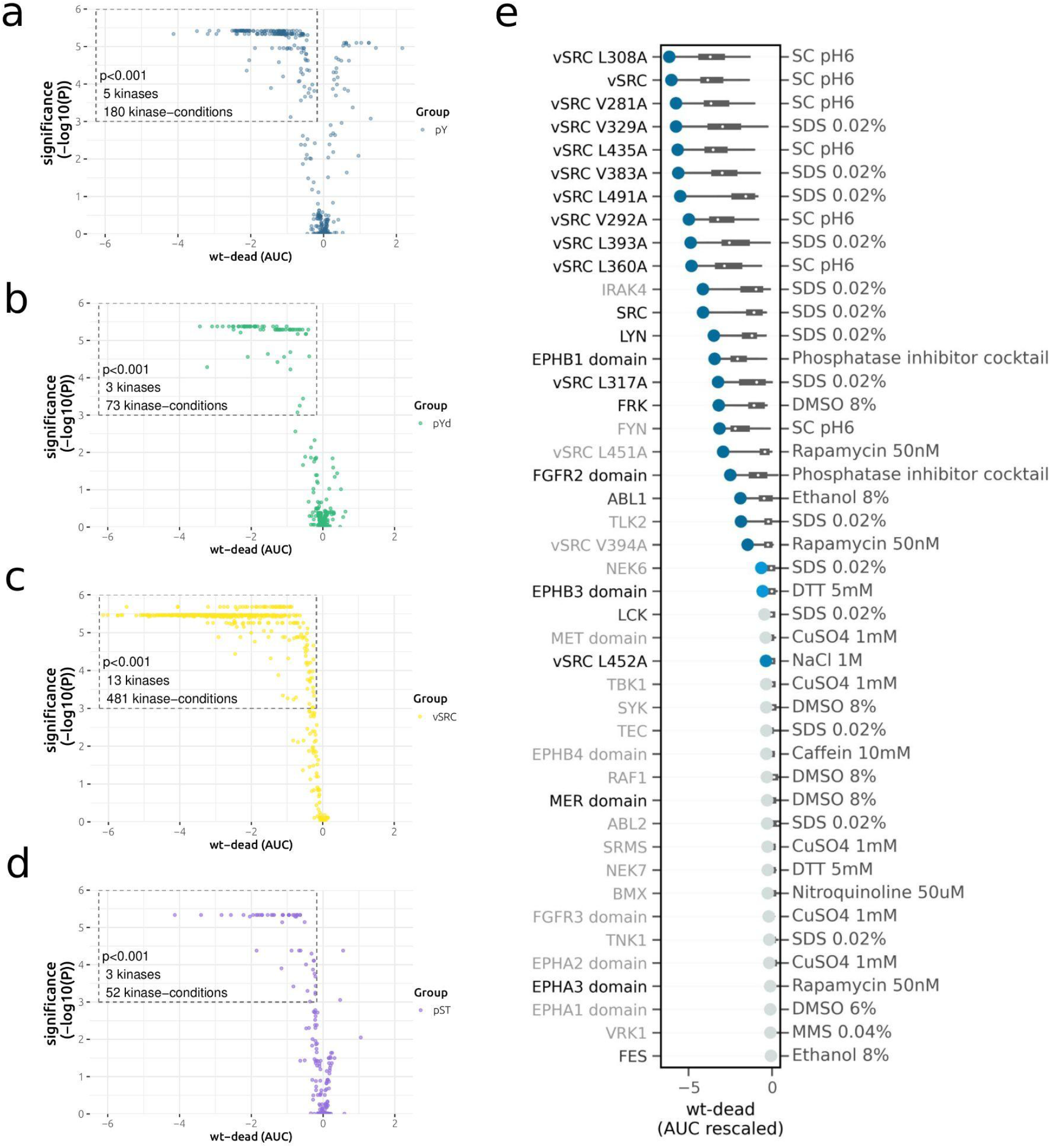
The effect of kinase expression (WT-dead) on fitness. Volcano plots for the fitness effect of different kinases separated by group: **a)** full-length tyrosine kinases (pY), **b)** tyrosine kinase domains (pYd), **c)** v-SRC and its mutants (vSRC), and **d)** human serine/threonine kinases (pST). The x-axis represents the difference in area under the curve (AUC) between growth curves for the WT kinase and corresponding kinase-dead mutant. The y-axis represents the FDR-adjusted p-value for the significance of the AUC difference between the WT and dead kinase. All kinase-condition pairs are represented in each volcano plot. **e)** The fitness scores (x-axis) of all the kinases (y-axis) measured from the fitness screen are shown. The condition with maximum deleteriousness, marked by a circular dot is indicated on the right side of the plot. The colours of the dots indicate the statistical significance of the fitness score: dark blue indicates P<0.001, light blue indicates 0.001<P<0.05 and grey indicates non-significance. The distribution of the fitness scores in all conditions is shown by the horizontal grey box plots. The colours of the kinases indicate their activity level from Western blotting analysis **(Figure S1 and S2)**: shown in black are kinases that were found to be active, grey indicates weakly active, and light grey indicates kinases that showed no activity.

**Supplementary figure 7:**
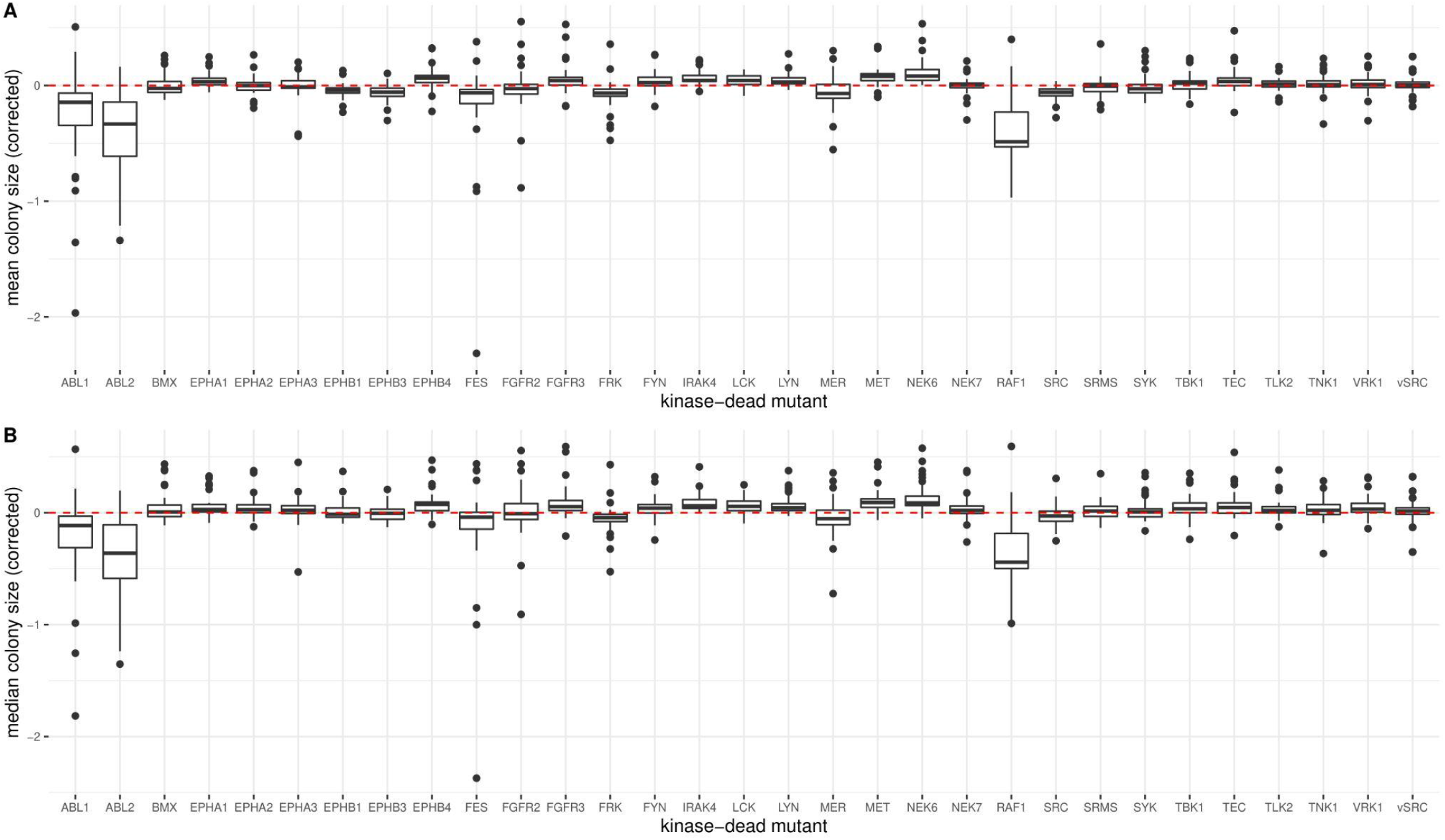
The effect of expression of the kinase-dead mutant on growth (colony size). The mean (top) and median (bottom) colony size for the kinase-dead mutant strains. Each colony size has been scaled with respect to a reference strain not expressing the kinase coding sequence (see Methods).

**Supplementary figure 8:**
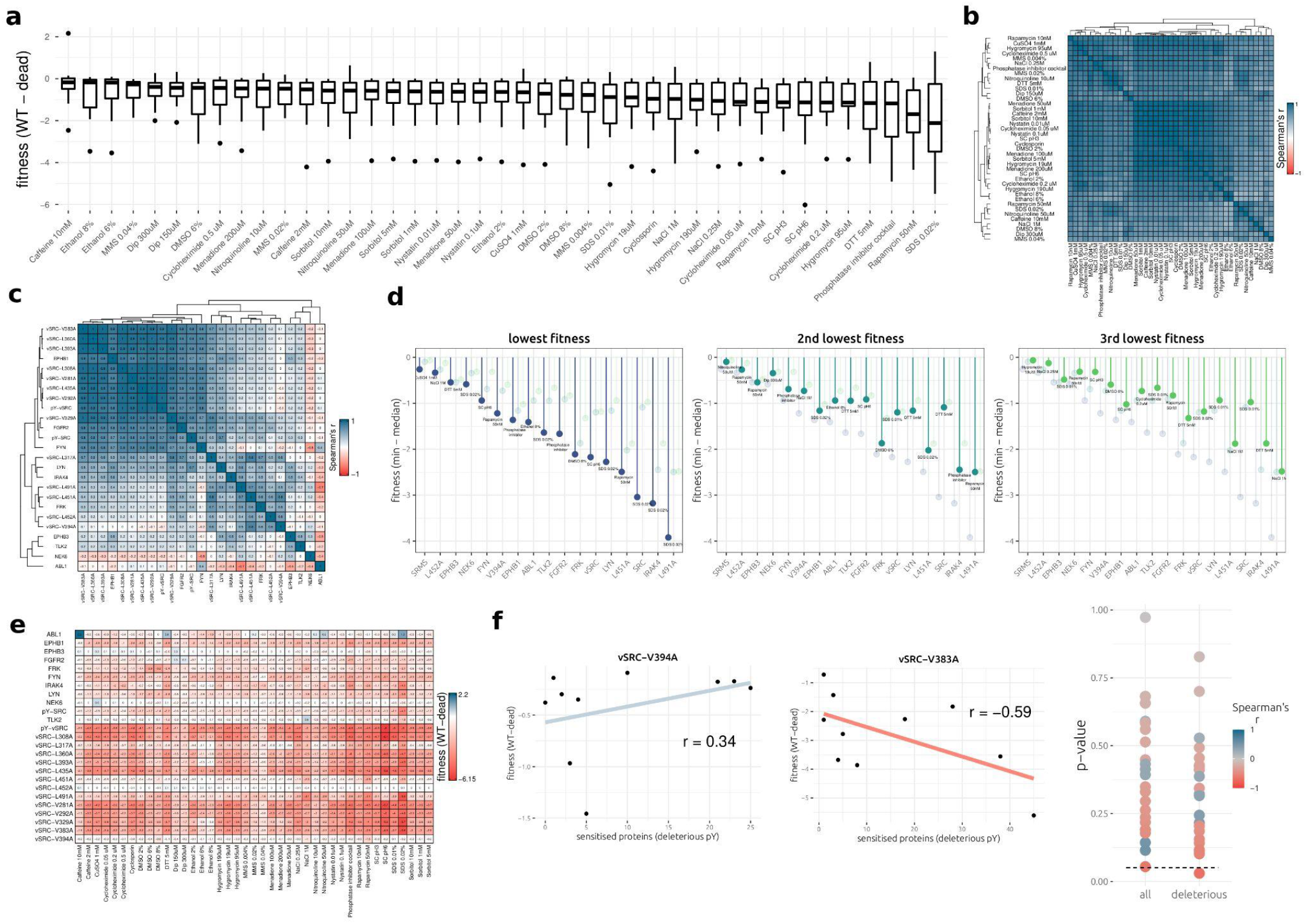
The difference in fitness effects between conditions. **a)** The fitness effects per condition across different kinases. To minimise redundancy, data for non-redundant vSRC mutants (WT, V491A, L451A, L452A, and V394A) and only one inactive kinase (SRMS) were used to generate the boxplots, alongside all active kinases. **b)** Correlation of fitness profiles (across kinases) between different conditions. Only kinases with at least one significant fitness defect were used to generate the correlation matrix. **c)** Correlation of fitness profiles (across conditions) between different kinases. Only kinases with at least one significant fitness defect were used to generate the correlation matrix. **d)** For each non-redundant kinase in the dataset, the difference between the minimum fitness and the median fitness across conditions. The second and third lowest fitnesses are represented in the middle and right panels, respectively. **e)** Heatmap of fitness scores (WT-dead) across all active kinases (rows) and tested conditions (columns). **f)** Cross-examination between kinase phosphosites detected in this study and the conditionally sensitive KO genes identified in (Viéitez *et al*, 2022). For the 10 conditions that overlap between this study and (Viéitez *et al*, 2022), the WT-dead fitness (y-axis) is taken and compared with the number of (predicted) deleterious pY mapping to conditionally sensitive proteins for each condition. (left) for the kinase vSRC V394A, with each point representing one condition. (middle) for the kinase vSRC V383A, with each point representing one condition. (right) The p-value and correlation coefficient for each regression, with each point representing one kinase. This is given for a correlation across ‘all’ pY that map to conditionally sensitive genes and for predicted ‘deleterious’ pY only. Significant negative correlations are expected if the differential fitness effects can be explained by loss-of-function (LOF) pY mapping to conditionally sensitive proteins.

**Supplementary figure 9:**
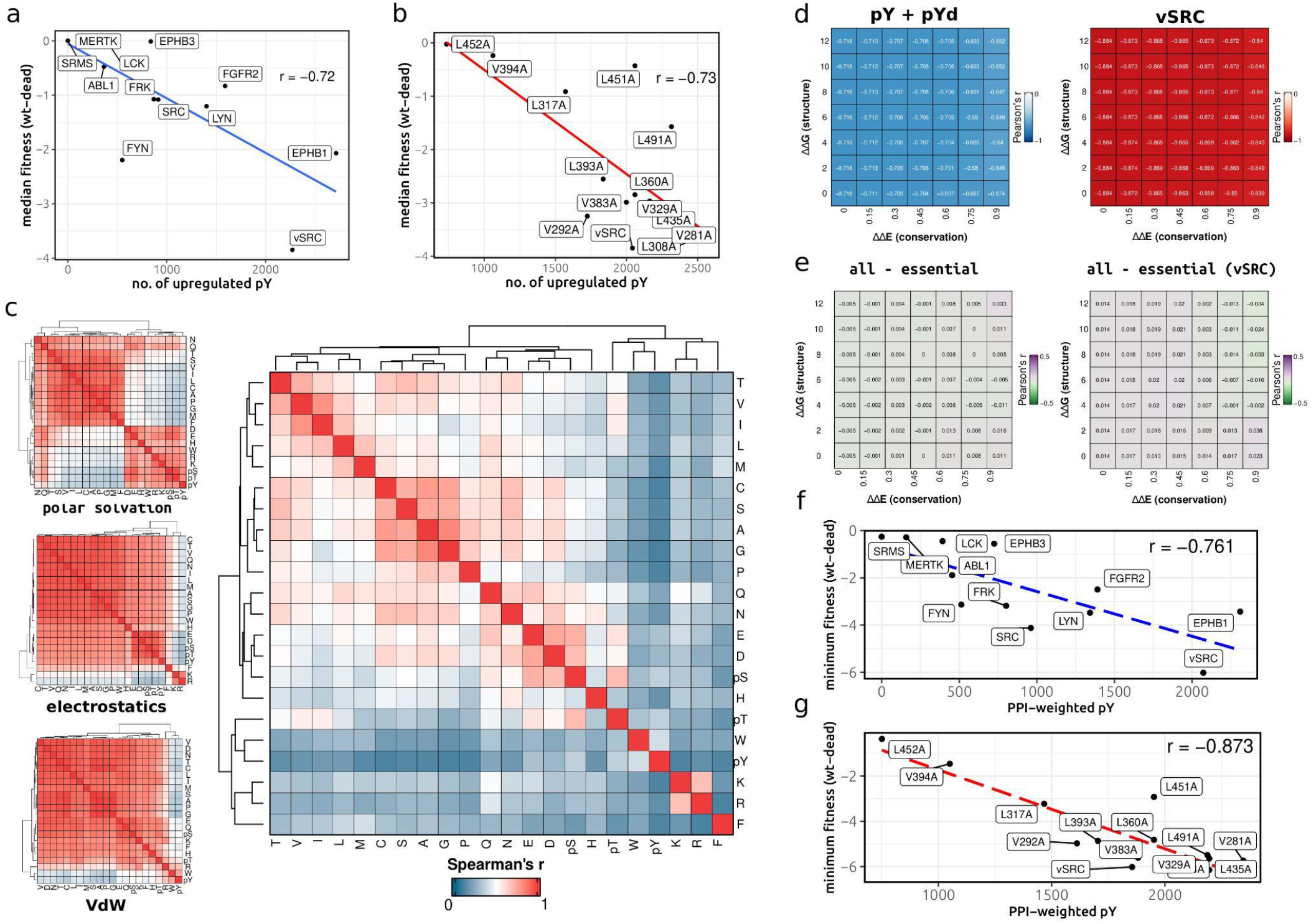
Relationship between kinase toxicity and phosphosite properties. **a)** Correlation between median fitness (across conditions, WT-dead) and the number of significantly upregulated pY sites (WT-dead) for each non-redundant kinase in the dataset. **b)** Correlation between median fitness (across conditions, WT-dead) and the number of significantly upregulated pY sites (WT-dead) for WT vSRC and the mutants generated in this study. **c)** For a large sample of spurious pY sites (n = ∼2300), the modified Y residue was mutated *in silico* to pY, pS, pT and the 19 other amino acids. Energy terms for the modification (Y to mutant/phosphosite) were calculated across all tested pY sites including polar solvation (top), electrostatic (middle), and Van der Waals (lower) energy terms. 22×22 correlation matrices were generated for each energy term to measure the similarity in effects of mutation and phosphorylation across various biophysical parameters. The mean correlation coefficient between energy terms was then taken to give an estimate of the average similarity between the 19 amino acids and pS, pT, and pY (see *Methods*). **d)** Correlation between minimum fitness (across conditions, WT-dead) and the number of significantly upregulated pY sites (WT-dead) for each non-redundant kinase in the dataset. Each value in the grid gives the Spearman’s correlation coefficient after filtering for pY sites on the basis of their destabilising effect (ΔΔG) or on the conservation of the modified residue (ΔΔE). Left: for non-redundant pY and pYd kinases. Right: for vSRC and its mutants. **e)** The same protocol is applied as in panel **d,** but this time the heatmap gives the *difference* in correlation coefficient using all spurious pY substrates against using essential proteins only (all - essential). **f)** Correlation between minimum fitness (across conditions, WT-dead) and the number of significantly upregulated pY sites (WT-dead) for each non-redundant kinase in the dataset. Each spurious pY is weighted so that pY predicted to map to many interfaces have a higher weight than non-interacting pY or those mapping to a small number of interfaces. **g)** The same correlation as in panel *f* but for vSRC and its mutants.

**Supplementary figure 10:**
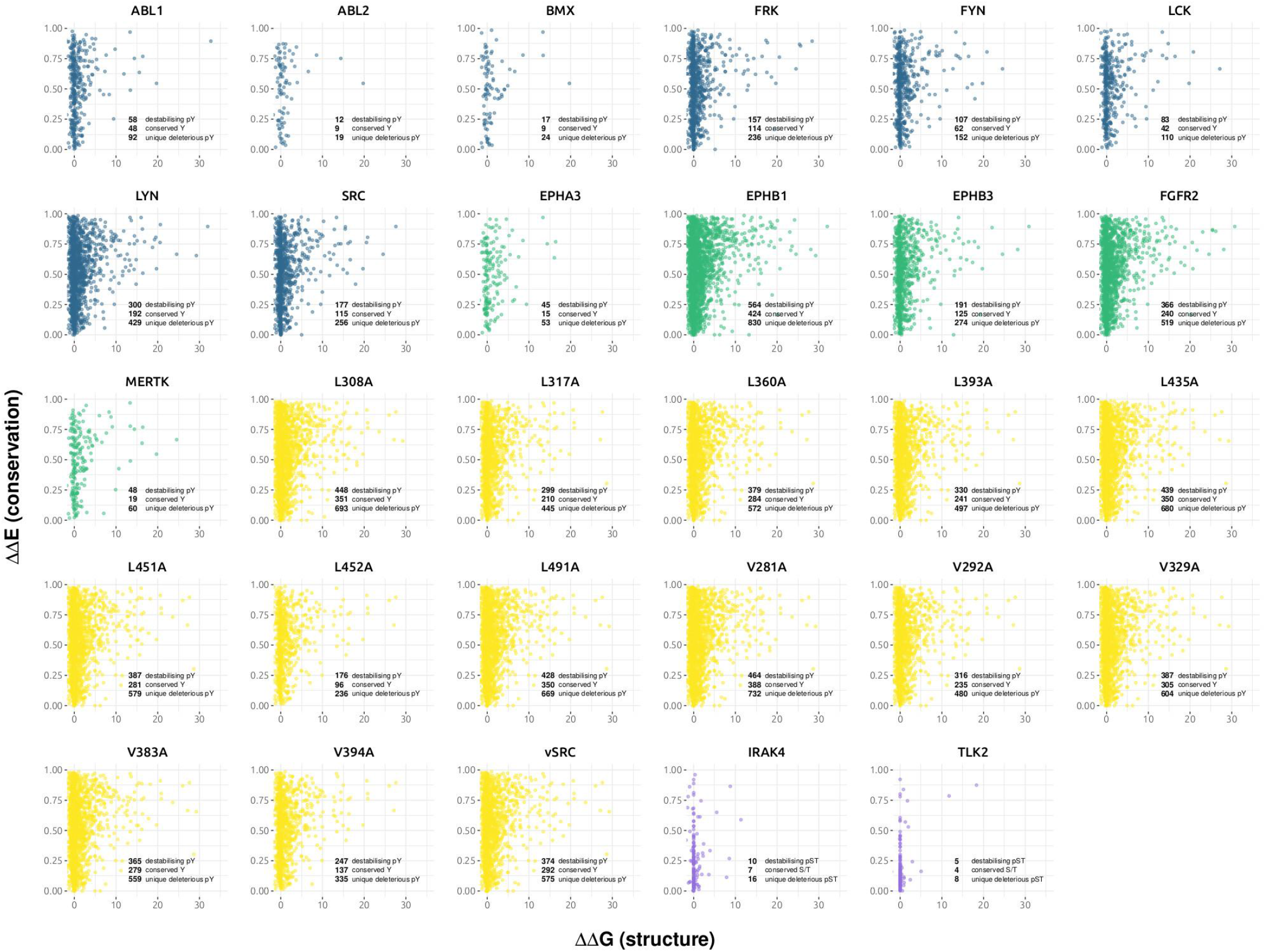
Variant effect prediction (VEP) of spurious phosphorylation with respect to protein stability (x-axis) and sequence conservation (y-axis). Higher ΔΔG values correspond to more destabilising pY (protein-level) whereas higher ΔΔE values correspond to pY mapping to more conserved Y positions. ΔΔG >2 and ΔΔE > 0.8 were the thresholds used to determine deleterious pY via structure and conservation, respectively. Kinases are coloured by their groups: blue (full-length tyrosine kinases), green (tyrosine kinase domains), yellow (vSRC and its mutants), and purple (pS/T kinases).

**Supplementary figure 11:**
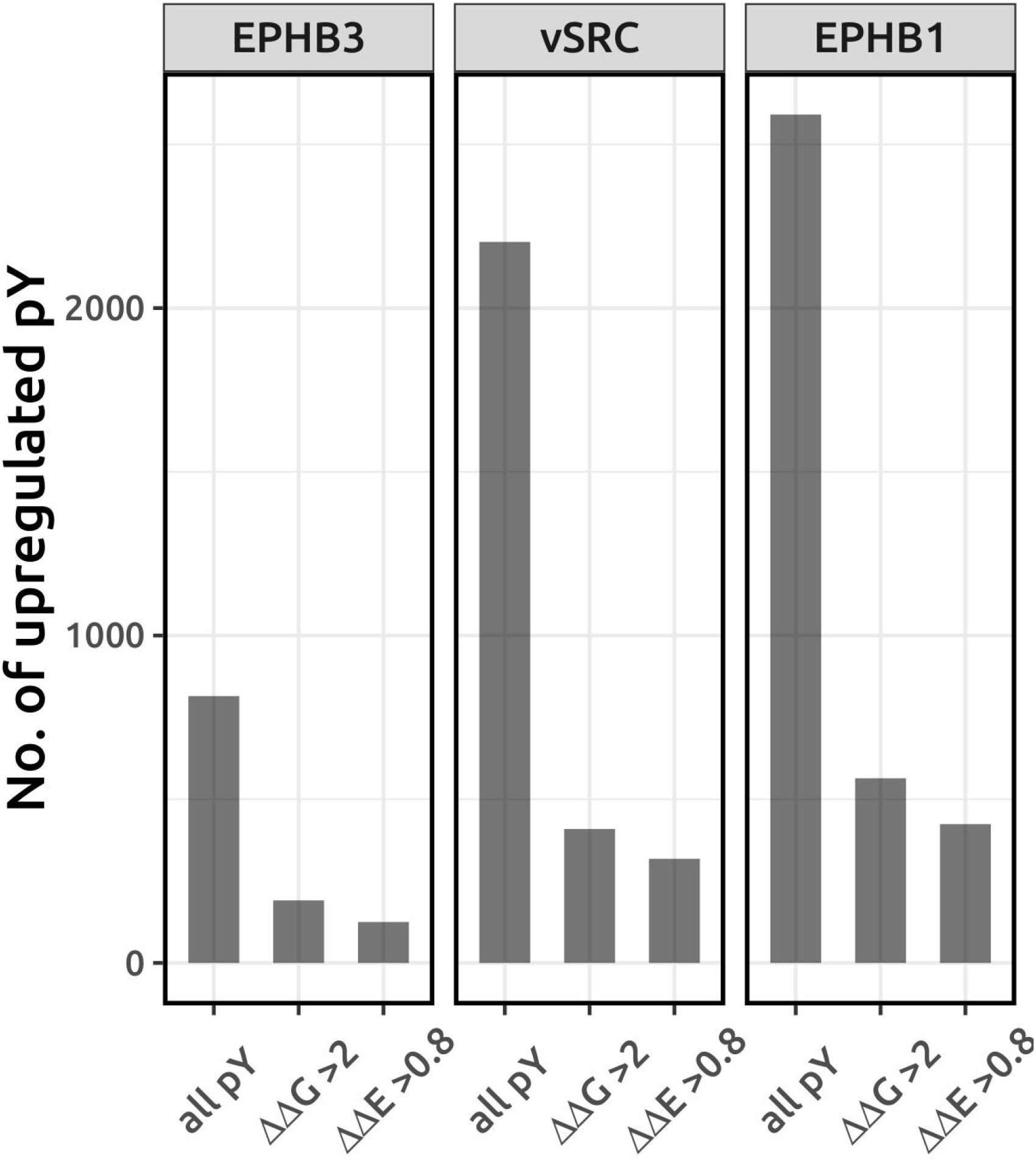
Spurious phosphorylation annotation for EPHB3, vSRC, and EPHB1. For EPHB3, vSRC, and EPHB1, the total number of upreglated pY, the number of destabilising pY (protein level, ΔΔG > 2), and the number of pY mapping to highly conserved positions (ΔΔE > 0.8). EPHB3 is an active kinase but with only a small effect on fitness (Figure 3e**, Figure S9a)**. vSRC and EPHB1 are among the kinases most deleterious for fitness, have the strongest pY profiles, and are used here to infer phosphorylation stoichiometry (Figure 3h, Figure 3i**, Figure S14, see Methods).**

**Supplementary figure 12:**
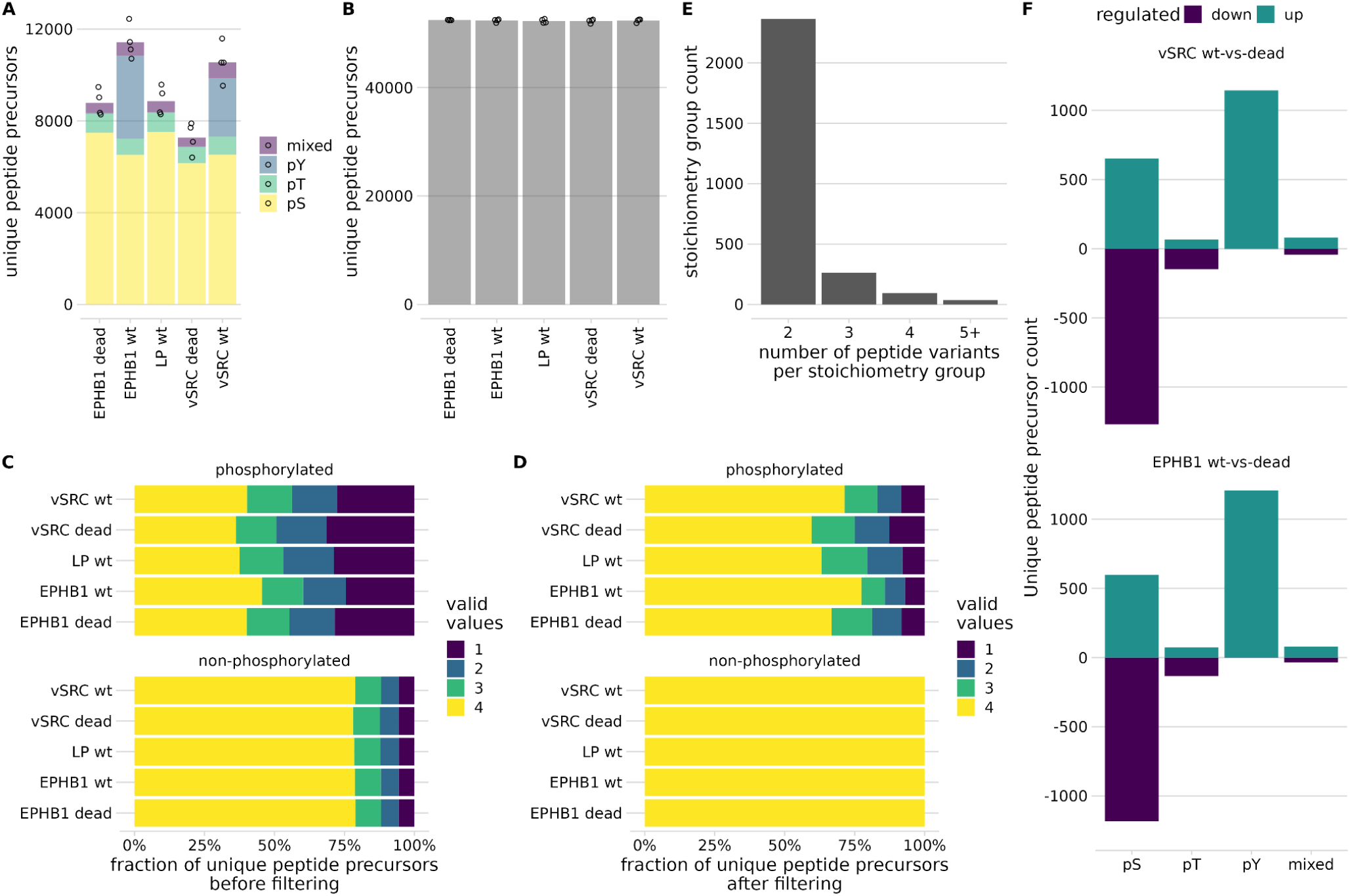
Calculation of stoichiometry values from (phospho-)proteomics data. **a-b)** Unique peptide precursor count of **a)** phosphoproteomic and **b)** proteomic data. **c-d)** The fraction of fully quantified peptide precursors per condition **c)** before and **d)** after quality control filtering. **e)** For stoichiometry groups with three or more peptide precursor variants, two or more isoforms of phosphorylated peptides were quantified. To simplify the correlation modelling, these stoichiometry groups were discarded. **f)** LIMMA analysis of phosphorylated peptide precursors reveals the same regulatory pattern of highly up-regulated phosphotyrosine containing peptide precursors, the non-phosphorylated counterparts of which were discarded for protein-level normalisation.

**Supplementary figure 13:**
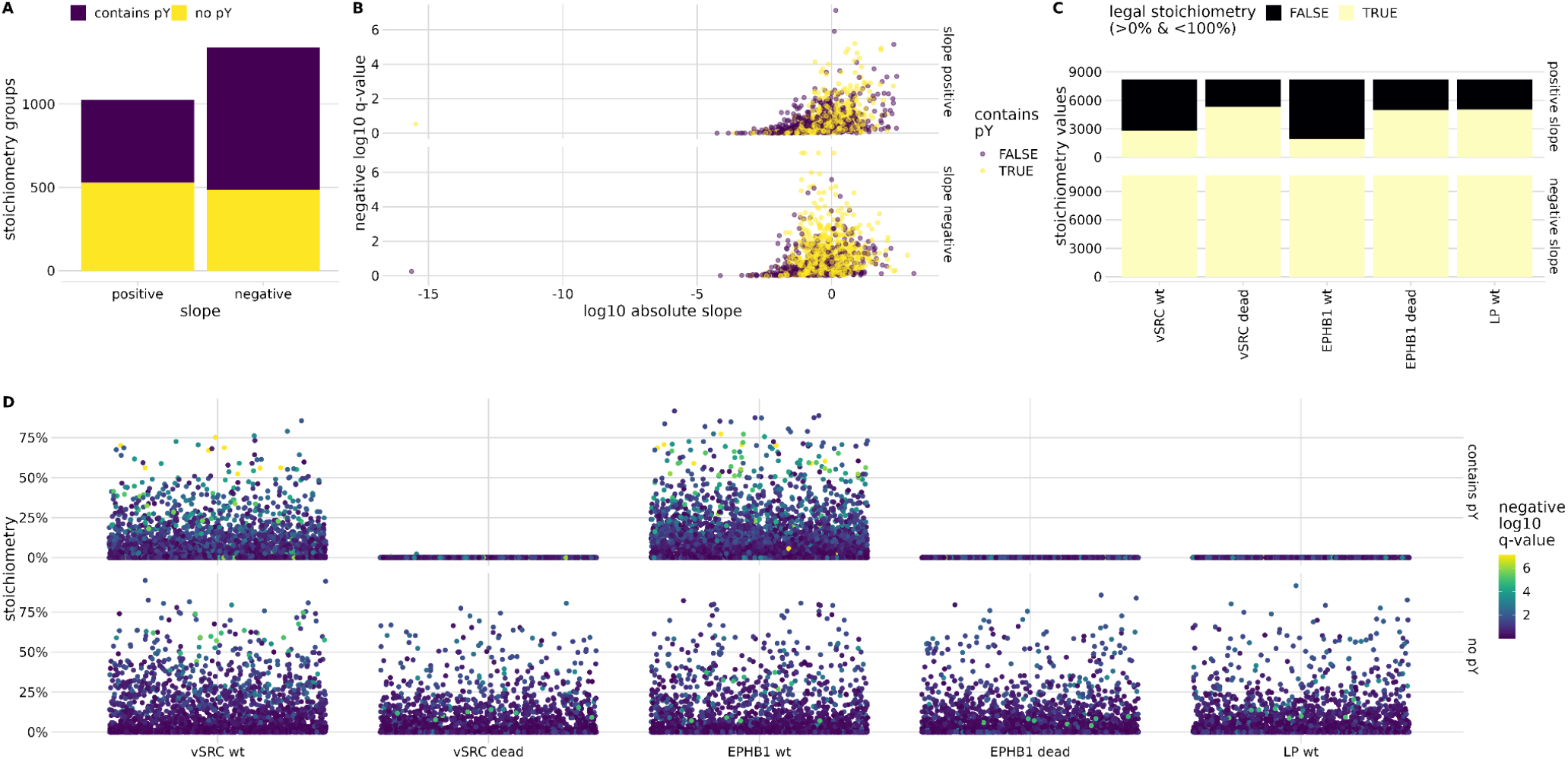
Quality control and filtering of stoichiometry values. **a)** A negative slope during stoichiometry group correlation is required for unambiguous stoichiometry value estimation. Peptide precursors containing phosphotyrosine are enriched in this fraction, most likely because they show higher absolute regulation between conditions. **b)** In the same manner, phosphotyrosine containing stoichiometry groups with negative slopes yield higher negative log 10 q-values. **c)** Stoichiometry groups with positive slopes can yield stoichiometry values below 0% or above 100% (termed “illegal”), and are therefore entirely discarded. **d)** The vast majority of peptide precursors containing phospho-tyrosine with significantly up-regulated stoichiometry values can be found in the kinase wild type conditions, as expected.

**Supplementary figure 14:**
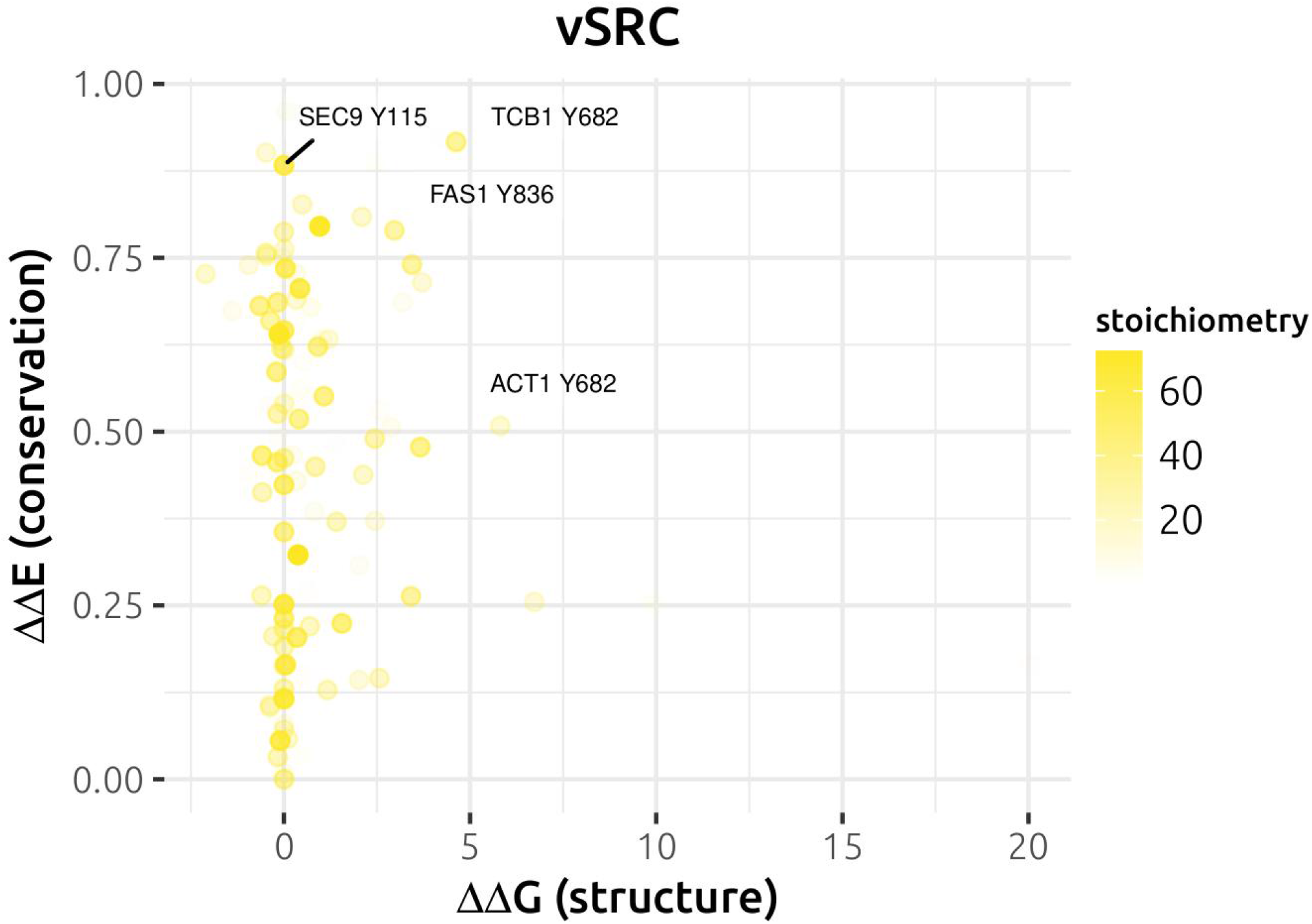
Stoichiometry of vSRC pY sites that are predicted to be deleterious for function. Higher ΔΔG values correspond to more destabilising pY (protein-level) whereas higher ΔΔE values correspond to pY mapping to more conserved Y positions. Higher inferred stoichiometries are given in dark yellow and lower inferred stoichiometries are given in light yellow.

**Supplementary figure 15:**
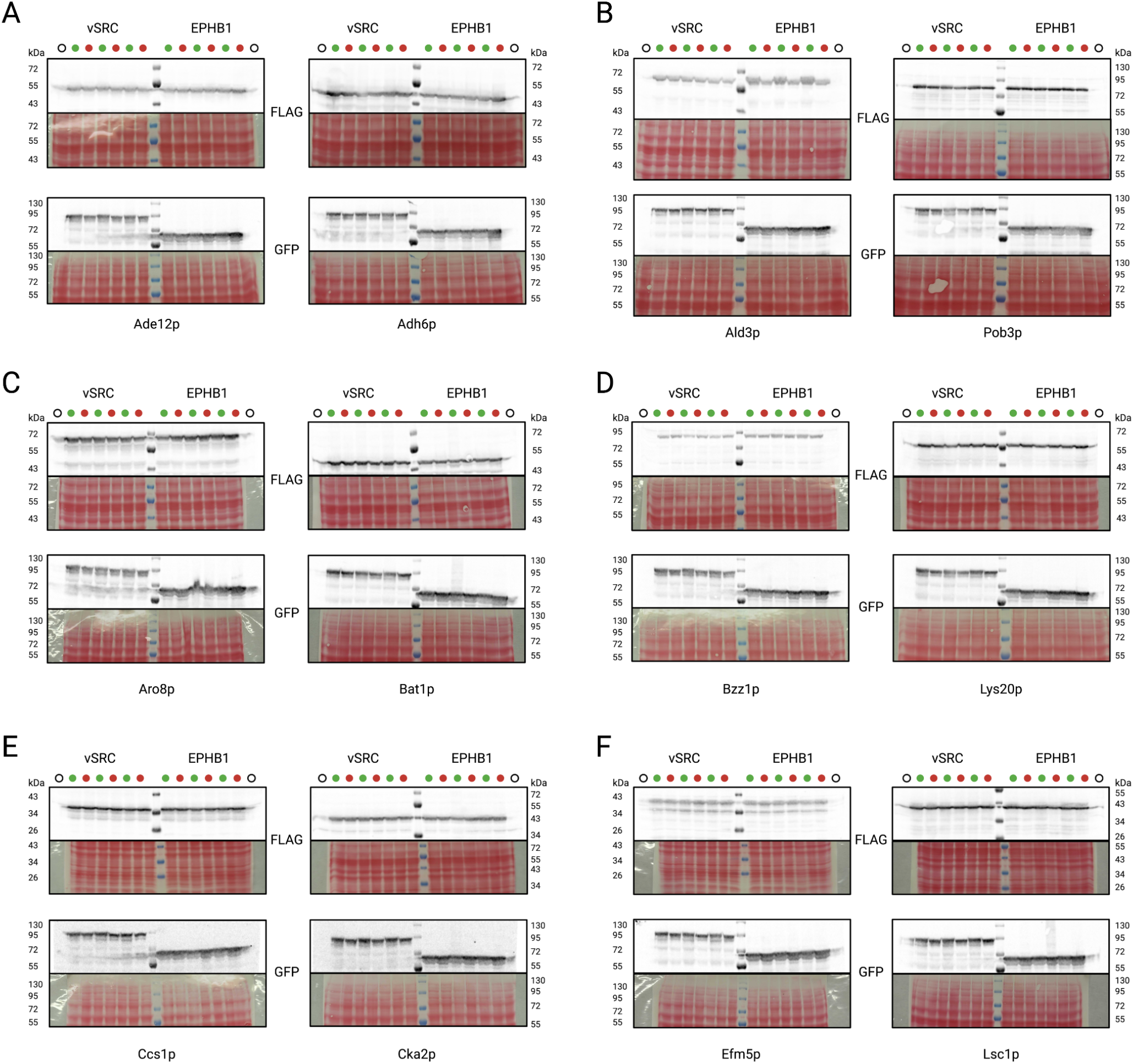
Protein with spurious phosphorylation sites analysed by Western Blot. For substrates with pY sites at intermediate-high stoichiometries and predicted to affect protein stability (high ΔΔG). Green dots show the active kinase version and red dots the kinase dead version. Empty circles show a negative control for the Western Blot. Protein (A) Ade12p, Adh6p, (B) Ald3p, Pob3p, (C) Aro8p, Bat1p, (D) Bzz1p, Lys20p, (E) Ccs1p, Cka2p, (F) Efm5p and Lsc1p were analysed. Proteins of interest were detected using an anti-Flag antibody. Kinases were detected using an anti-GFP antibody. Membranes were stained with Ponceau red to ensure proper protein transfer.

**Supplementary figure 16:**
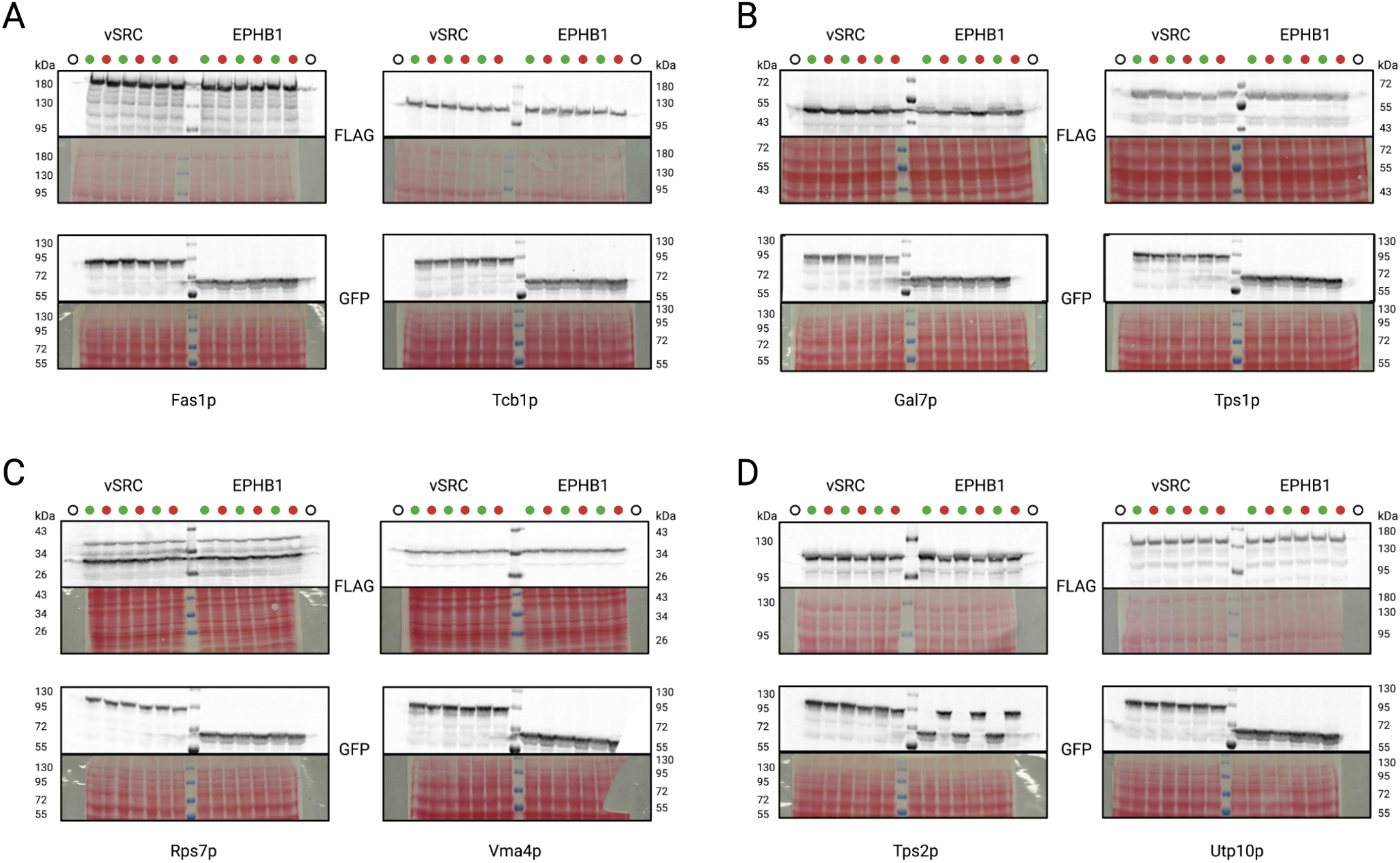
Protein with spurious phosphorylation site analysed by Western Blot. For substrates with pY sites at intermediate-high stoichiometries and predicted to affect protein stability (high ΔΔG). Green dots show the active kinase version and red dots the kinase dead version. Empty circles show a negative control for the Western Blot. Protein (A) Fas1p, Tcb1p, (B) Gal7p, Tps1p, (C) Rps7p, Vma4p, (D) Tps2p, and Utp10p were analysed. Proteins of interest were detected using an anti-Flag antibody. Kinases were detected using an anti-GFP antibody. Membranes were stained with Ponceau red to ensure proper protein transfer.

**Supplementary figure 17:**
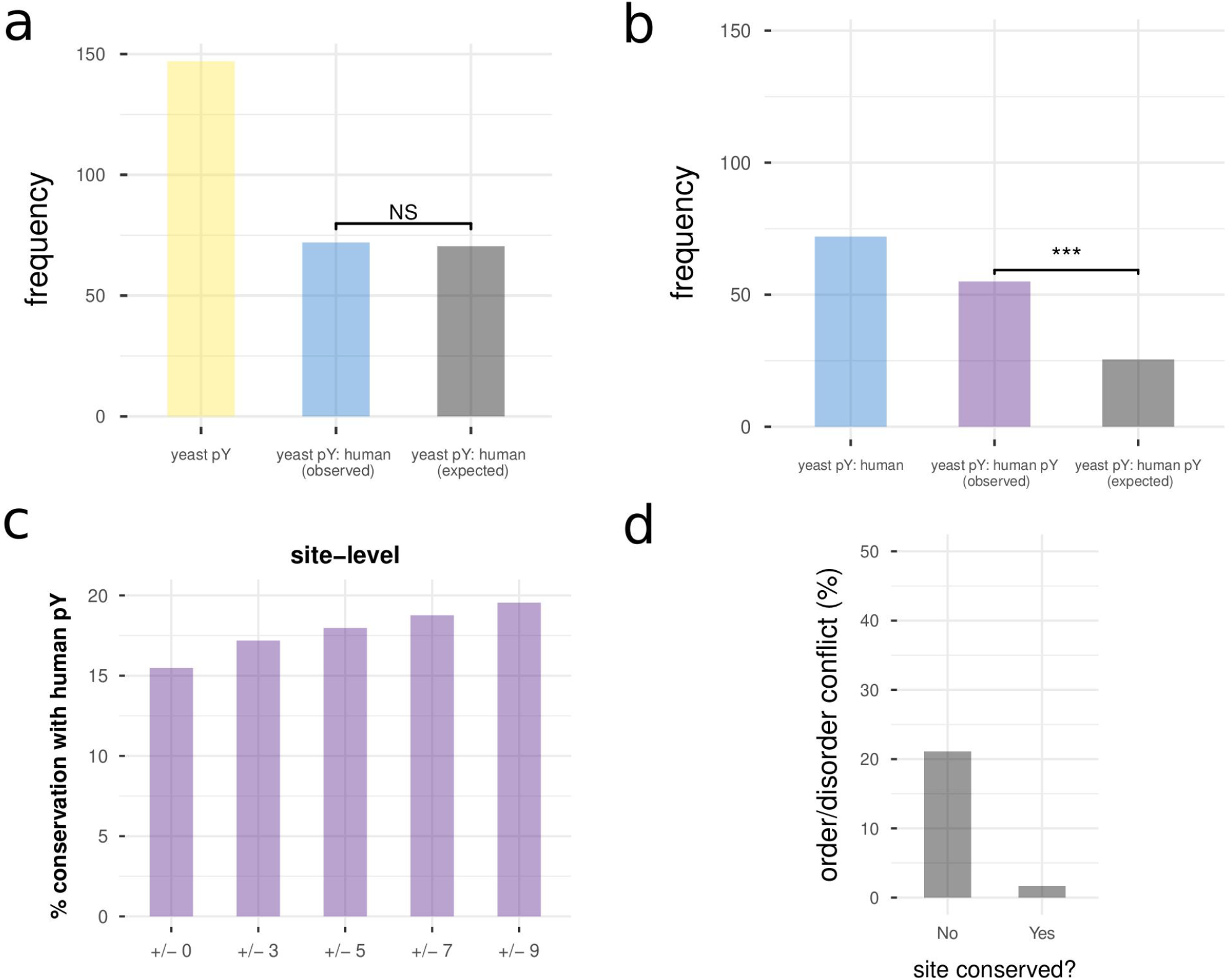
Supplementary analysis of conservation between spurious pY in yeast and native pY in humans. **a)** Human-yeast conservation at the whole protein level for native pY proteins in yeast. Yeast pY – number of unique native pY proteins in yeast. Yeast pY: human (observed) – number of *observed* native pY proteins in yeast with at least one ortholog in human. Yeast pY: human (expected) – number of *expected* unique native pY proteins in yeast with at least one ortholog in human, based upon the proportion of non-pY proteins that have at least one human ortholog. **b)** Human-yeast conservation at the level of whole protein pY phosphorylation for native pY proteins in yeast. Yeast pY: human – number of *observed* native pY proteins in yeast with at least one ortholog in human. Yeast pY: human pY (observed) – number of *observed* native pY proteins in yeast with at least one ortholog in human that is Y-phosphorylated. Yeast pY: human pY (expected) – number of *expected* native pY proteins in yeast with at least one ortholog in human that is Y-phosphorylated, based on non-pY proteins with a human ortholog and the proportion of those that are pY-phosphorylated. **c)** Relating to the site-based conservation analysis in Figure 4f, changes in the % pY conservation (for yeast spurious pY substrate with at least one pY-phosphorylated ortholog in human) with increasing sizes of an alignment window centred around the spurious pY position in yeast. **d)** For all pairs of human pY and yeast spurious pY on orthologous proteins, the percentage predicted as discordant in terms of the order/disorder prediction (order/disorder conflict).

**Supplementary figure 18:**
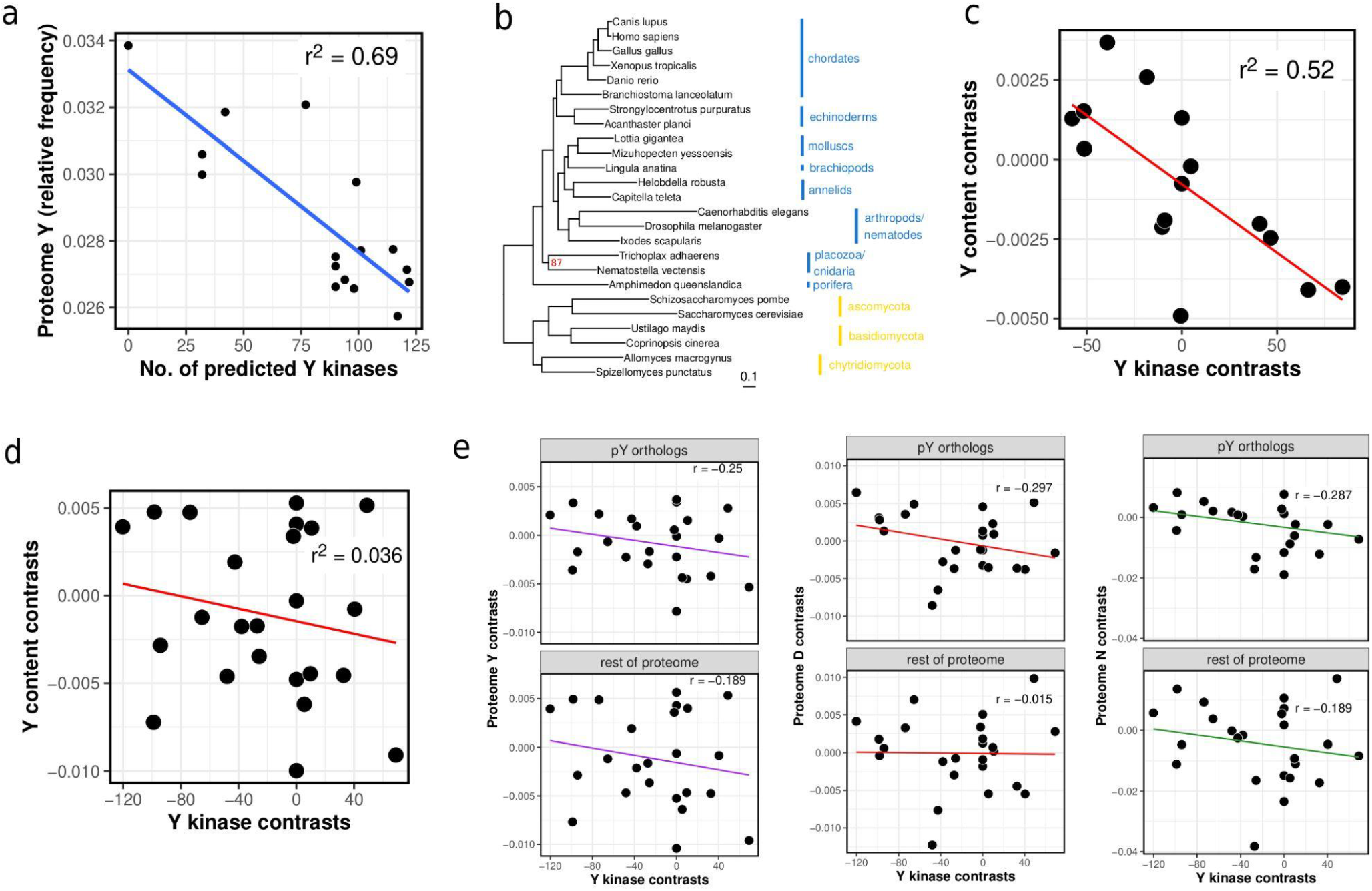
Supplementary analysis for testing metazoan counter-selection against pY at the level of the proteome. **a)** A reproduction of the (Tan *et al*, 2009) negative correlation between predicted Y kinases and proteomic Y content using modern proteome sequences and software for tyrosine kinase prediction. **b)** A species tree for all species used in the Figure 5a-d analysis (see Methods). The percentage of supporting ultrafast bootstrap replicates (/1000) was maximal (100%) except for the branch separating *N. vectensis* and *T. adhaerans* (87%). **c)** The result in panel **a** after applying a phylogenetic correction with the corresponding species tree using phylogenetic independent contrasts (Felsenstein, 1985). The species represented are the same as in (Tan *et al*, 2009). **d)** The result in Figure 5a after applying a phylogenetic correction with the corresponding species tree using phylogenetic independent contrasts (Felsenstein, 1985). The species used for this analysis are represented in panel **b. e)** Phylogenetically corrected correlation between the number of predicted tyrosine kinases (x-axis) and the proteome Y frequency (y-axis, left panel), D frequency (y-axis, middle panel), and N frequency (y-axis, right panel). In each case, the top panel represents the amino acid content only for orthologs of spurious pY substrates whereas the bottom panel represents the rest of the proteome (i.e. non-orthologs). The species used for this analysis are represented in panel **b.**

**Supplementary figure 19:**
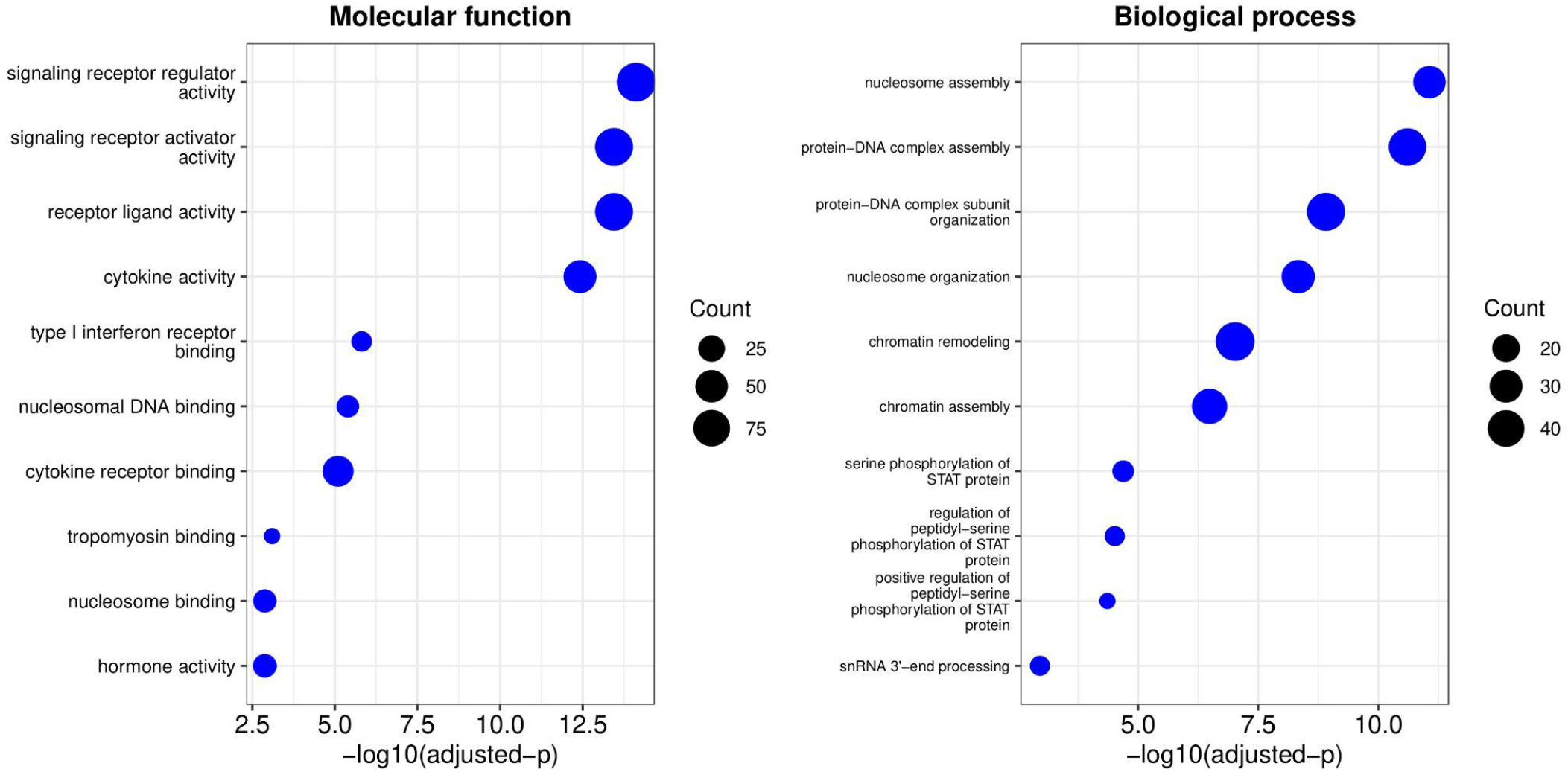
Gene ontology (GO) enrichment on human Y deserts. GO enrichment performed with respect to the molecular function and biological process terms. A tyrosine desert is defined as a protein where more than 50% of the protein length is missing tyrosine. Short proteins (length < 150 amino acids) were excluded from the analysis. ‘Count’ represents the number of unique proteins matching each GO term.

**Supplementary figure 20:**
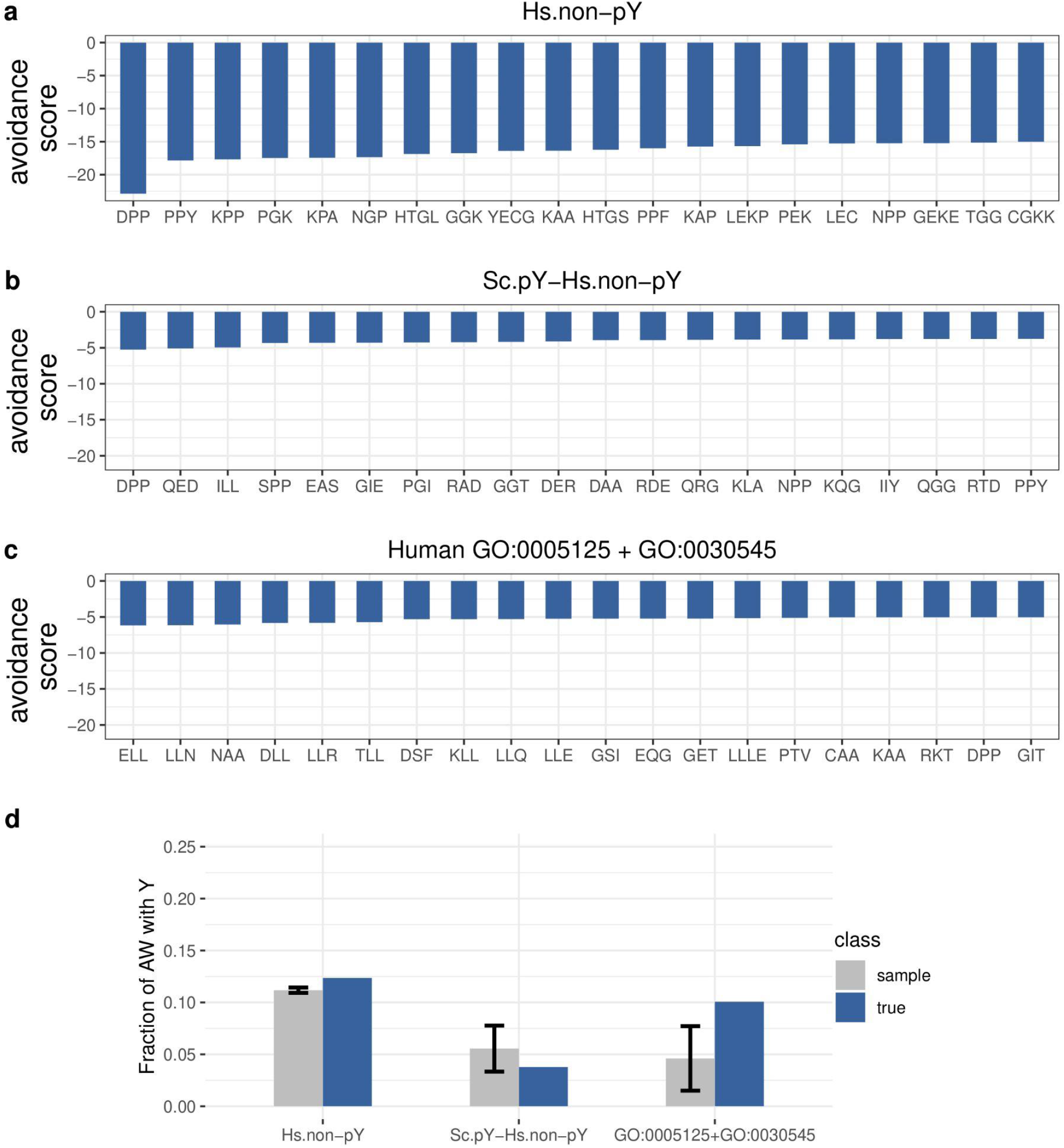
Analysis of ‘avoided words’ among protein sets of interest. **a)** Top 20 avoided words for human proteins not phosphorylated on tyrosine (Y). **b)** Top 20 avoided words for orthologs of spurious pY substrates (detected here) in *S. cerevisiae* that are not phosphorylated in humans. **c)** Top 20 avoided words for the gene ontology (GO) terms GO: 0005125 (cytokine activity) and GO: 0030545 (signalling receptor regulator activity) in humans that are enriched in tyrosine deserts **(Figure S19)**. **d)** For each of the three protein sets described above, comparison of the number of avoided amino acid words containing tyrosine (Y) for the real data compared with a random sample from the human proteome generated 500 times with matched sample size. Grey bars and error bars represent the median and standard deviation, respectively.

**Supplementary figure 21:**
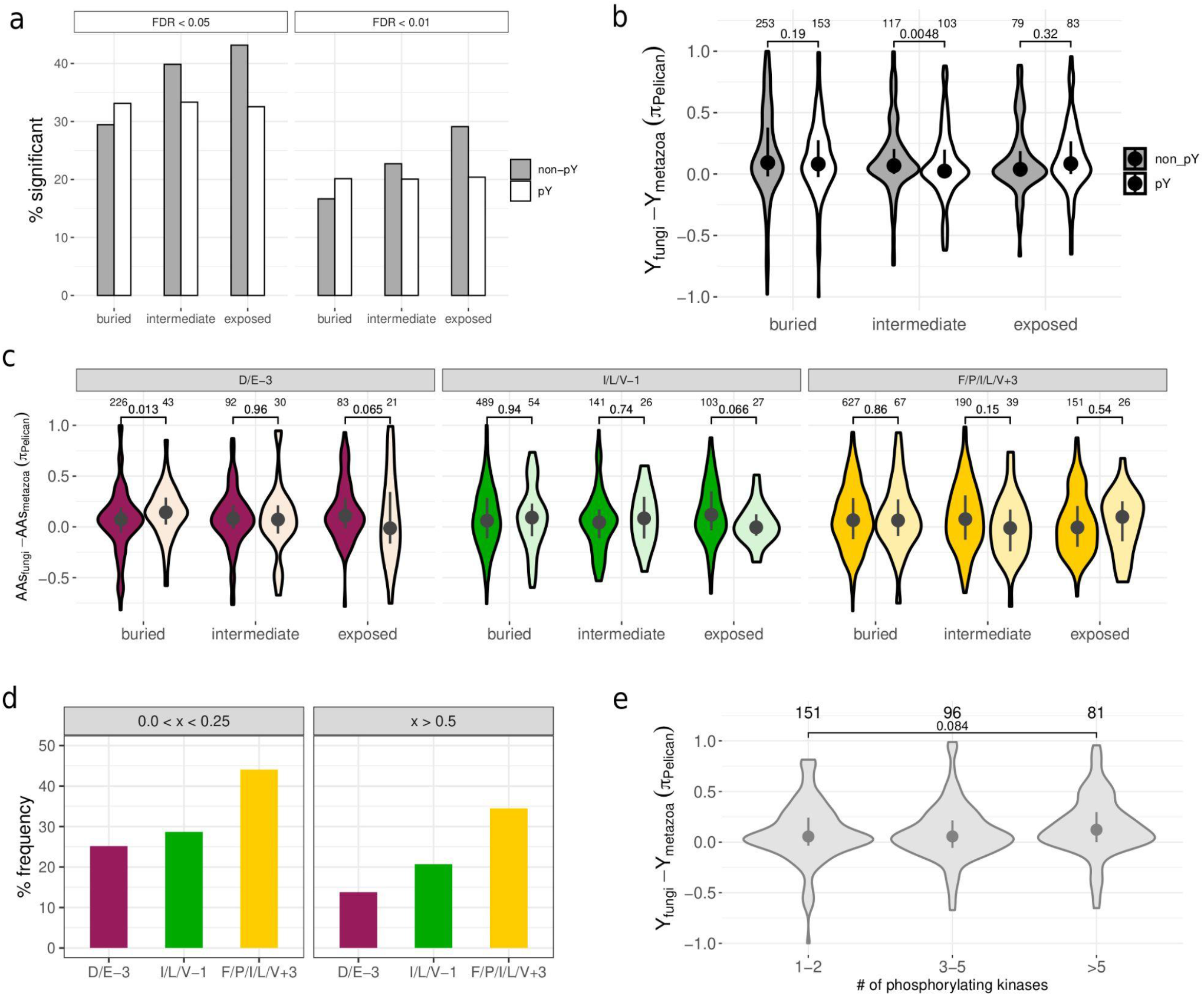
Supplementary results for the site-based analysis of pY counter-selection in metazoan species. **a)** For spuriously phosphorylated tyrosines (pY) and non-phosphorylated tyrosines (non-pY), the percentage with a significant shift in amino acid profile between animal and fungal species, as inferred by Pelican software (Duchemin *et al*, 2023)**. b)** The same analysis as in Figure 5f but after excluding non-pY sites that are poor candidates for Y phosphorylation on the basis of their sequence motif. Scores with a normalised motif score (0-1) maximum (across Y kinases) lower than 0.7 were excluded. Y kinase specificity models were constructed from data presented in (Sugiyama *et al*, 2019). **c)** Testing for counter selection against residues that are often found in Y kinase phosphorylation motifs: D/E-3, I/L/V-1, and F/P/I/L/V +3 (Deng *et al*, 2014; Li *et al*, 2023; Sugiyama *et al*, 2019). The analysis was performed and presented the same way as it is described in Figure 5f for the central Y residue. The non-phosphorylated sites are in the darker colour for all three panels. **d)** Determining whether these motif residues are more likely to be found in strongly counter-selected Ys (right) compared to weakly counter-selected Ys (left). The ‘x’ parameters refers to the difference in inferred Y preference between fungal and animal species (***π****_fungi_ **- π**_metazoa_*), with values closer to 1 being stronger candidates for Y counter-selection in animal species. **e)** Test to determine if sites that are phosphorylated by many Y kinases in this dataset are more strongly counter-selected than sites that are phosphorylated by only a small number of kinases. The analysis was performed and presented the same way as it is described in Figure 5f. This analysis excludes the vSRC mutants, which overlap strongly in terms of their substrate profiles.

## Methods

### Strain construction

Out of a total of 31 kinases, coding sequences (CDS) of 17 kinases were obtained from the Human Kinase ORF Kit (Yang *et al*, 2011), 3 from the Corwin *et al* study (Corwin *et al*, 2017), vSRC was kindly gifted by Dr. Josée Lavoie and the tyrosine kinase domains were gifted by Dr. Nicolas Bisson. Kinase-dead mutants for all kinases were generated by mutating the catalytic aspartate residue (D167 in human PKA) in the kinase active site to asparagine (Kung & Jura, 2016; Reinhardt & Leonard, 2023).

Following a methodology described in Ryan et al (Ryan *et al*, 2016), yeast (AKD0643, *ura3 his3 met15 leu2::GEM gal4::NAT gal1::stuffer-linker-GFP-HIS3*) competent cells were co-transformed with a pCAS plasmid (Addgene plasmid 6084747) expressing both the gRNA for the stuffer (Dionne et al 2021) and *Streptococcus pyogenes* Cas9, and a donor DNA. For the construction of all the strains, a single gRNA was used that targeted a genomic location (*GAL1* replaced by a stuffer sequence) flanked by *Gal1p* and C-terminal GFP tag. The donor DNA with flanking homology arms was generated by amplifying the CDS sequence using oligos containing homology with the target sequence (Oligo sequences in **Table S2**). As a linker between the CDS and the C-terminal tag, a small flexible protein sequence (GGGGSGGGGS) that has minimal effect on the structure of the linked proteins was used. Transformed cells were selected on YPD plates containing 200 μg/mL G418 (plasmid selection, Bioshop Canada) and 100 μg/mL NAT (*gal4*::*NAT* deletion, Cedarlane Labs). To allow plasmid loss, randomly selected colonies were grown in YPD media without G418 for 2 days. Subsequently, the plasmid loss was confirmed by no growth on YPD plates containing 200 μg/mL G418. The genomic integrations of the CDSs were validated by PCR and their sequences were confirmed by Sanger sequencing (Oligo sequences in **Table S2**).

To construct the mutants of the kinases the same method was followed, except that as donor DNA, either PCR product from the amplification of the mutated CDS from a plasmid or overlapping PCR with the mutagenesis oligos, was used. All oligo sequences are included in **Table S2**.

### Cell pellet preparation

Pre-cultures of yeast cells were prepared by overnight growth at 30 °C with agitation, in synthetic complete (SC) medium (1.74 g/L Yeast nitrogen base without amino acid without ammonium sulfate, 1 g/L Monosodium Glutamate salt, 20g/L Glucose, 1.34 g/L complete drop out, see **Table S15** for the drop out mixture). Pre-cultures were inoculated at an O.D. of 0.05 in a fresh SC media and grown for 12 hrs at 30 °C with agitation. Then, the cells were pelleted. The pellet was washed with PBS, and frozen at −80 °C. In the ‘induced’ condition, the fresh medium contained 1 μM beta-estradiol inducer, whereas in the ‘not induced’ condition, the fresh media contained the same volume of DMSO as the inducer.

### Activity assay by Western Blotting

The cell pellets for the activity assays were prepared as described in the *Cell pellet preparation* section above. For cell lysis, the pellets were resuspended in Laemmli buffer (16,25 mM Tris-HCl pH 6,8, 10% glycerol, 2% SDS, 0,01% bromophenol blue and 5% β-mercaptoethanol) and heated at 95°C for 10 mins. The protein extracts were then spun down and the supernatants were migrated on 10% polyacrylamide gels (volumes were normalised based on the final OD of the different yeast strains). Once separated, the proteins were transferred to nitrocellulose membranes for Western blotting and correct protein loading was validated with ponceau staining. All membranes were blocked in a 5% milk solution and then incubated for 12h at 4°C with either the mouse anti-phosphotyrosine 4G10 (05321, Millipore) or the rabbit anti-GFP (Thermo A11122) primary antibodies. Signals following a 1h incubation at room temperature with the appropriate HRP-conjugated secondary antibodies (anti-mouse NEB 7076S or anti-rabbit NEB 7074S) were acquired using the Clarity Western ECL Substrate (Bio-Rad) and an Amersham Imager 600RGB (GE Healthcare).

### Protein abundance assay by Western Blotting

For this validation, 20 proteins containing pY sites at intermediate-high stoichiometries and predicted to affect protein stability were selected and tagged with a 3X-FLAG epitope. Each protein was tagged in four different backgrounds (vSRC, vSRC kinase dead, EPHB1 and EPHB1 kinase dead) and tagging was validated by Western blots. To check the effect of phosphorylation on the protein of interest we grew each strain in triplicates with 100 nM beta-estradiol for 6 hours at 30°C (from 0.15 OD/ml to ∼ 1 OD/ml). Following the growth, the cells were spun down and washed 1 time with sterile water. The pellet was then frozen for further use. The pellet was resuspended at 0.05 U OD/µL in lysis buffer (Complete Mini, Roche and PhosStop, Roche) and 250 µl was processed for cell lysis. Cell lysis was performed by adding glass beads to the cell suspension and then vortexed on a Turbomix for 5 min. Then, 25 µL SDS 10% was added to the mixture and boiled for 10 min. Finally, the tube was spun at 16,000*g* for 5 min. Samples for migration were prepared by mixing 17.5 µL clear supernatant, 2.5 µL DTT 1M and 5 µL LB5X (250mM Tris-Cl pH 6.8, 10% SDS, 0.5% Bromophenol Blue, 50% glycerol). The percentage of acrylamide gel was determined based on theoretical protein size, as small proteins will have better separation on 15% gel than large proteins on 8% gel. Following protein separation on SDS-PAGE, they were transferred onto a nitrocellulose membrane at 0.8 mA/cm^2^ for 1h15. Each membrane was stained with Ponceau rouge to confirm proper loading and appropriate transfer. The membrane was blocked overnight in a blocking buffer (Intercept® (PBS) Blocking Buffer, 927-70003, Mandel Scientific). The next morning, Anti-FLAG M2 antibody (Anti-Flag M2, F3165-1MG, Millipore-Sigma) was applied to the membrane for 30 min to the protein of interest and with Anti-GFP (Millipore-Sigma, 11814460001) for kinase expression. Secondary antibodies (Anti-Mouse 800, LIC-926-32210, Mandel Scientific) were applied for 30 min and then the membrane was imaged with an Odyssey Fc instrument (Licor, Mandel Scientific) in 700 and 800 channels.

### Flow cytometry competition assay

For this experiment, kinase and kinase-dead versions were competed against a WT parental strain expressing mCherry (*PDC1-mCherry*). The two competing strains were mixed at a ratio 1:1 in 200 µL of SC complete media. Each competition was performed with and without beta-estradiol. The competition was performed over 3 days (∼30 generations). Each day, for the next round of the competition, cultures were diluted 10 µL in 190 µL of sterile fresh media. Samples were analysed using a flow cytometer (Guava EasyCyte instrument, Cytek Bio) for 5000 events each. The bimodal distributions of the fluorescence intensities were fitted to 2-component Gaussian mixture models, to obtain thresholds to classify the events with signal and those without signal. The threshold for each sample was calculated as the intensity level at the intersection of the fitted curves, lying between the distributions of the events with and without signal. Because the cells are expected to have either GFP or mCherry fluorescence, but not both, the events classified as not having signal in channels corresponding to GFP and mCherry fluorescence and the ones with signal in both the channels (doublets) were discarded. Among the remaining events, the relative frequency of a given strain harbouring a kinase in a sample was calculated as the proportion of events without mCherry signal.

### Fitness measurements across conditions

In total, 77 strains (31 wt kinases, 31 kinase-dead mutants, 13 mutants of vSRC, 1 strain without the CDS (empty landing pad) and the background strain BY4741) were rearrayed onto four 384 OmniTray plates using a robotically manipulated pin tool (S&P robotics). After growth for 2 days at 30 °C, the plates were then used for condensation onto 1,536 OmniTray plates containing SC complete media (1.74 g/L Yeast nitrogen base without amino acid without ammonium sulfate, 1 g/L Monosodium Glutamate salt, 20g/L Glucose, 1.34 g/L complete drop, 20 g/L agar), using the robotically manipulated pin tool. On these plates, the positions of the strains and their 16 replicates were randomised in order to account for any location-specific undue growth effects. Also, the background strain BY4741 was positioned at the border of the plates (two rows and two columns) in order to reduce any border effects. These plates were incubated for two days at 30 °C. Next, these plates were used as source plates to replicate onto the 41 different growth conditions with and without beta-estradiol (0 and 1000 nM in DMSO), referred to as destination plates. The destination plates were incubated at 37 °C in a SpImager custom platform. The images of the plates were acquired every 1h30 hours for 3 days. The conditions selected are known to trigger stress responses and were used previously in (Dionne *et al*, 2021).

### Fitness screen: analysis

Images obtained from the SpImager custom platform were used to obtain growth parameters using python based pyphe software (Kamrad *et al*, 2020). To generate the input for the pyphe software, the python based scikit-image tool (van der Walt *et al*, 2014) was used. Using scikit-image, images were first preprocessed to retain only the area containing the colonies in the images. This was done by removing the borders of the plates from the images. The images were then converted to grayscale and their intensity range was inverted. The resulting images were provided as input to the `pyphe-quantify` tool using these parameters: ‘`batch --grid auto_1536 --s 0.1`’. The measurement of the circularity of the colonies was used to filter the colonies with abnormal shapes. Next, the `pyphe-quantify` tool was used with the following parameters: ‘`timecourse --grid auto_1536 --s 0.1`’, to obtain the growth curves. For each growth curve, the colony size at the initial time point was subtracted from the colony sizes at every time point. This correction helped in reducing the effect of possible unequal replication of colonies by the pin tool. Next, the corrected growth curves were provided as input to the `pyphe-growthcurves` with default parameters, to obtain the growth parameters. Additionally, the growth curves were used to measure the Area Under the Curve (AUC) according to composite Simpson’s rule using python based scipy package’s (Jones *et al*, 2016) `integrate.simps` module, with default parameters.

Growth conditions in which CuSO_4_ was added to the media produced red coloration of the colonies. These plates were preprocessed to only include intensity from the red channel only. The following steps of the analysis were the same as other plates.

The values of the growth parameters were rescaled by those of the reference strain without the CDS. This within-plate rescaling allowed carrying out comparisons between different plates. The fitness score for a wild type kinase was calculated as a difference between its rescaled growth parameters and that of the corresponding kinase-dead mutant. The statistical significance of the difference was tested using Mann–Whitney U test corrected for multiple tests.

### (Phospho-)proteomics: sample processing and measurement

Cell pellets were thawed and mixed with 750 μl of freshly prepared urea lysis buffer (8 M urea, 75 mM NaCl, 100 mM Tris pH 8.5) and tubes were filled up with 0.5 mm Zirconia/Silica beads. Cells were lysed via 4 cycles of bead beating (1 min each with 30 sec cooling on ice in between). After removing any water sticking to the tube, a hole was poked using a needle, and the lysate was transferred to a new tube by centrifugation at 420 g for 1 min. Cell debris was discarded and lysate transferred to a new tube after 10 min of 21k g centrifugation at 4 °C. Lysates were sonicated for 20 sec each and BCA assay was performed to determine protein concentration. Lysates were reduced with 5 mM DTT (30 min, 55 °C), alkylated with 20 mM CAA (30 min, room temperature in the dark) and quenched with another 5 mM DTT (15 min, room temperature).

For (phospho-)proteome preparation, 250 μg of protein lysate were processed using an adjusted R2-P2 protocol (Leutert *et al*, 2019). In brief, 250 μg protein was diluted to 1 μg/ul, with 2 μg beads per 1 μg protein added. 750 μl of ACN were added to precipitate proteins. Washing was performed with 1 ml each of 100% ACN, 100% MeOH, 100% ACN, 80% ACN and 80% EtOH. Digestion was performed using 2.5 μg trypsin diluted in 200 μl of 100 mM Tris pH 8.5 for 12h at 37 °C. For proteome measurement, digests were acidified by adding 20 μl 50% FA to 5% total. Supernatant was separated from bead remnants on a magnetic rack, and centrifuged for 10 min at 21k g. Supernatant was filtered through a C8 StageTip (50 ml MeOH, 50 μl ACN, sample, 50 μl 80% ACN in 0.1% FA), dried in a speed-vac and stored at −20 °C until MS measurement. For phospho-proteome measurement, digests were acidified by adding 20 μl 50% TFA to 5% total and 800 μl ACN. Supernatant after centrifugation for 10 min at 21k g was used in R2-P2 phospho-peptide enrichment. Enrichment was performed using 62.5 μl of a 5% solution Fe^3+^-NTA magnetic beads (PureCube Fe-NTA MagBeads, Cube Biotech), washed 3x with 80% ACN in 0.1% TFA and eluted into 100 μl of 2.5% NH_4_OH in 50% ACN. Phospho-peptides were acidified with 10 μl of 50% FA to 10% final, filtered through a C8 StageTip (50 ml MeOH, 50 μl ACN, sample, 50 μl 80% ACN in 0.1% FA), dried in a speed-vac and stored at −20 °C until MS measurement.

For MS measurement, peptides were re-solubilized using 4% ACN in 0.1% FA. 2.5 μl of sample was analysed by nLC-MS/MS using an Orbitrap Exploris 480 Mass Spectrometer (Thermo Fisher) equipped with an Easy1200 nanoLC system (Thermo Fisher). Peptides were loaded onto a 100 μm ID × 3 cm precolumn packed with Reprosil C18 3 μm beads (Dr. Maisch GmbH), and separated by reverse-phase chromatography on a 100 μm ID × 35 cm analytical column packed with Reprosil C18 1.9 μm beads (Dr. Maisch GmbH), housed in a column heater set at 50 °C, using two buffers: (A) 0.1% FA in water, and (B) 80% ACN in 0.1% FA in water at 450 nl/min flow rate.

For LCMS measurement, peptides were separated over a 60 or 90 min LC gradient ramping from 6% B to 32% in 43 or 69 min, respectively, and washing at 95% before re-equilibrating to 3%. Data-independent acquisition (DIA) was performed as a staggered window approach (Pino *et al*, 2020) (**Table S16**). A full MS1 scan was recorded after every DIA cycle at 60k resolution with standard AGC target and automatic maximum IT. 2 distinct DIA cycles that were shifted 12 Th covered an effective range of 438 to 1170 m/z (phospho) and 363 to 1095 (proteome). One DIA cycle consisted of 30 windows of 24 Th size at 30k resolution, 27% NCE HCD, charge state 3 and AGC target 1000%. Data-dependent acquisition (DDA) analysis was performed using a 3 second cycle approach. A full MS1 scan was recorded every 3 seconds at 120k resolution with 300% AGC target, automatic maximum IT and 450 to 1150 m/z. MS2 precursors were selected using filters MIPS peptide, 5e3 intensity threshold, charge states 2-6 and 30 sec dynamic exclusion. MS2 scans were recorded at 30k resolution, 27% NCE HCD, 1.6 Th isolation window, charge state 3 and standard AGC target. For both DDA and DIA acquisition, gas phase fractionated (GPF) measurements were performed of pooled samples. For DDA-GPF, 5 measurements covering 4x 125 Da and 1x 200 Da of the total scan range were performed. For DIA-GPF, 7 measurements covering 100 Da each with 25 windows of 4 Th isolation width were conducted.

### (Phospho-)proteomics: data processing and analysis

MS raw files were converted to mzML files using MSConvert v3.0.22248-ce619fc (Chambers *et al*, 2012). For staggered DIA files, filters peakPicking at vendor msLevel 1- and demultiplex with optimization overlap_only and massError 10 ppm were used. Database and spectral library search was performed using FragPipe v18 (Yu *et al*, 2022) with a FASTA file combining yeast and human kinases (wild-type and kinase dead) downloaded from Uniprot on 2021-11-08. DDA, DDA-GPF and DIA-GPF files were used to create a spectral library in FragPipe. Default settings for strict trypsin and high resolution MS data were combined with variable modifications 15.9949 on M, 42.0106 on protein N-terminus, 79.96633 on STY, −17.0265 on peptide N-terminal QC and −18.0106 on peptide N-terminal E. DIA files were searched using the created spectral library in FragPipe and quantified with DIA-NN (Demichev *et al*, 2019) in mode “Any LC (high accuracy)”. Phospho PTM localization was performed adding the parameter “--var-mod UniMod:21,79.966331,STY --monitor-mod UniMod:21”.

LCMS data analysis was performed using custom scripts in RStudio v2023.03.0 and R v4.2.3, with packages including data.table, ggplot2, Biostrings and cowplot. All scripts are available at https://gitlab.com/public_villenlab/spurious-phosphorylation. DIA data was filtered at ≤ 1% FDR level with Q.Value and Global.Q.Value. Phospho-peptide data was further filtered at PTM.Site.Confidence ≥ 75%. For stoichiometry calculation, the phospho-peptide data was additionally filtered at ≤ 1% FDR level with PTM.Q.Value for extra stringency. Dimensionality reduction via tSNE was performed using the R package Rtsne (Maaten & Hinton, 2008; van der Maaten, 2014) with perplexity 10, theta 0.0 and max iterations 5000. Statistically significant regulation was tested using the R packages samr (Tusher *et al*, 2001) and limma (Ritchie *et al*, 2015). For SAM, default settings were used. For LIMMA, contrast matrices were created between kinase wild type and kinase dead conditions, as well as vSRC mutants and vSRC kinase dead conditions. For each test, only peptide precursors that were quantified in at least 75% of replicates in at least one condition were used. Of the remaining PCMs, if all replicates of one condition were not quantified, intensities for this condition were imputed. Imputation was performed the same way as the Perseus computational platform (Tyanova *et al*, 2016), using a normal distribution of both conditions in the test (sd = 0.3 x sd_of_data; mean = mean_of_data - 1.8 x sd_of_data). LIMMA testing was performed using functions lmFit, constrasts.fit and robust eBayes. P-values were multiple testing corrected using Benjamini Hochberg. Motif analysis was performed using the R package dagLogo (Ou *et al*, 2020). Enriched amino acids were filtered by p-value ≤ 0.05 and visualised using the R package ggseqlogo (Wagih, 2017).

### (Phospho-)proteomics: stoichiometry

Phosphorylation site stoichiometry values were calculated by correlating intensities of phosphorylated peptide precursors and their non-phosphorylated counterparts (Presler *et al*, 2017; Hogrebe *et al*, 2018). To adjust for changes in protein abundance between wild type and kinase dead conditions, proteome maxLFQ intensities were calculated and used to normalise peptide precursor intensities (Cox *et al*, 2014). Crucially, only non-phosphorylated peptide precursors whose phosphorylated counterparts did not exhibit significant regulation as determined by LIMMA were included in the maxLFQ calculations. For LIMMA testing only, intensities were imputed if peptide precursors could only be quantified in max 25% of one condition, and min 75% in the other. maxLFQ-normalised peptide precursors were filtered: 1) to retain only the highest abundant charge state per modified peptide; 2) to have phosphorylated peptide precursors fully quantified in at least one condition; 3) to have all non-phosphorylated peptide precursors fully quantified in all conditions; 4) ro retain only peptide precursors with exactly 1 phosphorylated counterpart to the non-phosphorylated peptide precursors (called a “stoichiometry group”). Finally, the remaining phosphorylated peptide precursors were imputed to be 0 whenever not quantified. After stoichiometry calculation, stoichiometry groups yielding a positive slope were removed, as they could yield stoichiometry values below 0% or above 100% (called “illegal”).

### Structural bioinformatics

Structural models for kinase substrates were sourced from AlphaFold2 where possible (Jumper *et al*, 2021; Varadi *et al*, 2022). The solvent-accessible surface area (SASA) of tyrosine residues was calculated with DSSP v. 3.1.4 (Kabsch & Sander, 1983) using the dssp() function in the R package bio3d (Grant *et al*, 2021). The SASA was normalised by the maximum Y accessibility (255 Å^2^) to give the relative solvent accessibility (RSA) (Tien *et al*, 2013). The Y disorder state was predicted using an AF2-based approach (Akdel *et al*, 2022; Piovesan *et al*, 2022), where a smoothened RSA in a +/-12 amino acid window surrounding the central Y was used as a proxy for protein disorder prediction **(Figure 2a)**. Tyrosines with a smoothened RSA greater than or equal to 0.581 were predicted as disordered (Piovesan *et al*, 2022).

The effect of phosphorylation on the Gibbs free energy of folding (ΔΔG) was predicted using FoldX (Schymkowitz *et al*, 2005; Studer *et al*, 2016). All protein structures were first repaired using the FoldX RepairPDB command (--ionStrength=0.05, --pH=7, --water=CRYSTAL --vdwDesign=2). The ΔΔG of phosphorylation (Y to pY) was then calculated using the BuildModel command, with 5 runs per phosphosite. Finally, the effect of pY phosphorylation on protein interactions was predicted by calculating the ΔΔG for complex formation using the PSSM command, with 5 runs per phosphosite. Destabilising pYs were called using a standard ΔΔG threshold of 2 kcal/mol (Nishi *et al*, 2011; Wagih *et al*, 2018; Høie *et al*, 2022).

Structural annotations of protein-protein interfaces for *S. cerevisiae* were derived from the ‘Highest confidence interfaces’ dataset of Interactome INSIDER (Meyer *et al*, 2018). This resource includes annotations of PDB structures and homology models present in Interactome3D (Mosca *et al*, 2013). We supplement this data with a recent AF2-based screen of protein interactions in *S. cerevisiae* (Humphreys *et al*, 2021). Interface residues were determined using the approach outlined in (Meyer *et al*, 2018) i.e. residues with an RSA > 0.15 and with a change in SASA > 1.0 Å^2^ upon complex formation. Interface residues were filtered further to have an AF2 predicted aligned error (PAE) less than 8 Å. Finally, we include Interactome INSIDER’s machine learning-based predictions of interface residues for our analysis **(Figure 2e and Figure S5i)**

Calculation of spurious-to-native and native-to-native 1D and 3D distances were made with respect to the ultra-deep reference yeast phosphoproteome presented in (Leutert *et al*, 2023). Sites are considered 1D proximal if they are within 4 amino acids of each other. Sites are considered 3D proximal if they are within 8 Å of each other (Humphreys *et al*, 2021). pS/pT and pY sites mapping to low-confidence positions (pLDDT < 70) were excluded from this analysis. Phosphosites were added to the AF2 structures using the PyTMs package in PyMOL (Warnecke *et al*, 2014).

### Conservation-based variant effect prediction (VEP)

Homologs of all unique spurious substrates were retrieved using hhblits v. 3.3.0 (Remmert *et al*, 2011). Each sequence was searched with one iteration (n=1) against the UniRef30 database (2022 version) that contains non-redundant UniProtKB sequences clustered at 30% sequence identity (Suzek *et al*, 2015; Mirdita *et al*, 2022). Homologs were retrieved with an e-value threshold of e^-10^ and the resulting alignment filtered to retain only positions found in the query sequence (i.e. the *S. cerevisiae* copy). Hits with 50% or more gapped positions were then removed (Høie *et al*, 2022). Variant effect prediction (VEP) based upon the sequence alignment was performed with GEMME software for all alignments containing at least 20 hit sequences (Laine *et al*, 2019). Results were taken from the combined evolutionary model (‘evolCombi’) and then rank-normalised (within proteins) such that the most harmful mutations were assigned a score of 1 and the least harmful a score of 0.

GEMME and similar softwares perform VEP for amino acid mutations but not for phosphorylation. We therefore weight GEMME scores for the 19 amino acids by their biophysical similarity to pY. Estimates of AA-to-pY biophysical similarity were obtained by mutating a large sample of spurious pY (n = ∼2300) in Silico from Y to the 19 other amino acids and pS/pT/pY using the PSSM command in FoldX (Schymkowitz *et al*, 2005). Energy terms were taken for the backbone Hbond, VdW, electrostatics, polar solvation, hydrophobic solvation, VdW clashes, sidechain entropy, mainchain entropy, torsional clashes, backbone clashes, helix dipole and ionisation energy. Correlation matrices were generated for each energy term, and then average correlation of each amino acid to pY was used to calculate a pY-weighted mean of the GEMME scores. Annotations for essential proteins were retrieved from YeastMine (Balakrishnan *et al*, 2012).

### Evolutionary bioinformatics (human-yeast conservation)

Orthologs of the kinase substrates were retrieved from the vertebrate, fungi, and metazoa databases of Ensembl Compara using the Ensembl REST API (Vilella *et al*, 2009; Yates *et al*, 2015). Tyrosine-phosphorylated proteins in human were annotated using PhosphoSitePlus (Hornbeck *et al*, 2015), while requiring at least 5 supporting sources to ensure high-confidence annotations (Ochoa *et al*, 2020; Kalyuzhnyy *et al*, 2022). Assessing the conservation of kinase-substrate relationships **(Figure 4c)** was performed using the ProtMapper resource for human substrate annotations (Bachman *et al*, 2022), and the substrates identified here (via MS/MS) for the yeast substrate annotations.

Protein domain annotations were assigned using pfam (Mistry *et al*, 2021), and protein abundance data was retrieved from PaxDB (Wang *et al*, 2015). Paralog annotations in *S. cerevisiae* were retrieved from (Marchant *et al*, 2019) for the paralog conservation analysis in **Figure 4e**. Conservation analysis was performed on paralog-specific pY-phosphorylated peptides only. For paralog pairs with at least one paralog-specific phosphopeptide each, the paralog pairs were aligned using MAFFT L-INS-i (Katoh & Standley, 2013). Aligned Y/Y pairs were designated as ‘single’ for non-conservation (pY/Y or Y/pY) or ‘pair’ for phosphosite conservation (pY/pY).

For the generation of alignments across orthologous groups, all unique orthologs were first filtered to remove redundant orthologs at a sequence identity threshold of 95% using CD-HIT (Fu *et al*, 2012). Ortholog sequences were aligned to the *S. cerevisiae* copy using the ‘phmmer’ command from HMMER 3.3 software and then the HMM-sequence alignments were concatenated to give a single aligned sequence for every ortholog. The resulting alignments between the *S. cerevisiae* copy and its orthologs were used to determine site-level conservation between *S. cerevisiae* pY and human pY **(Figure 4f).** For further evolutionary analysis (described in the section below), potential false positive homologs were removed by excluding any predicted ortholog that aligned to less than 50% of the sequence of the *S. cerevisiae* copy. The trimAl software was then used to remove positions from the alignment with more than 20% gaps across all sequences (Capella-Gutiérrez *et al*, 2009), and an additional filter was applied to remove spurious homologs in the MSA also using trimAl (parameters: -resoverlap 0.75, -seqoverlap 80) (Capella-Gutiérrez *et al*, 2009).

The structural analysis of pY accessibility (RSA), order/disorder, and effect on stability (ΔΔG > 2) was undertaken using the same approach outlined above (‘Structural bioinformatics’ section). Human proteins were considered to be pY-phosphorylated if supported by at least five sources from PhosphoSitePlus (Hornbeck *et al*, 2015). For native pY sites we use a high-confidence group found to be endogenously phosphorylated in both (Lanz *et al*, 2021) and (Leutert *et al*, 2023). For the spurious pY group, we conservatively exclude any pY site found to be endogenously phosphorylated either in (Lanz *et al*, 2021) or (Leutert *et al*, 2023).

### Evolutionary bioinformatics (testing for pY counter-selection)

Non-redundant reference proteomes were taken from UniProt for each of the species analysed (UniProt Consortium, 2023). Kinases were predicted and classified (e.g. as tyrosine kinases) using the Kinannote software (Goldberg *et al*, 2013).

The species tree in **Figure S18b** was constructed using orthologs from 19 metazoan species and 6 fungal species selected to give broad taxonomic coverage, with each phylum represented by at least one species if a reference proteome was available. Orthologous groups were used if they contained at least one ortholog from each of the 25 species. Orthologous sequences were aligned using MAFFT L-INS-i (Katoh & Standley, 2013), and alignment positions with more than 20% gaps were removed using trimAl (Capella-Gutiérrez *et al*, 2009). Orthologs aligning to less than 50% of the *S. cerevisiae* sequence (i.e. the query sequence) were considered as spurious homologs and removed. For species containing more than one ortholog in an alignment (i.e. one-to-many or many-to-many copies), the copy with the highest sequence identity to the *S. cerevisiae* copy (i.e. the query sequence) was retained and the others removed.

The resulting set of alignments was then used to construct a species tree with IQ-TREE2 using the ‘concatenation/supermatrix’ approach (Chernomor *et al*, 2016; Minh *et al*, 2020). Optimal substitution models for each alignment were determined automatically within IQ-TREE2 using ModelFinder (Kalyaanamoorthy *et al*, 2017), and branch confidence was determined with 1000 ultrafast bootstrap replicates (Hoang *et al*, 2018).

For the correlation of proteome Y content against the number of Y kinases, the phylogenetic non-independence of data points was corrected using phylogenetic independent contrasts (**Figure 5b**, **Figure S18c-e)** (Felsenstein, 1985), as it is implemented in the R package ape (Paradis & Schliep, 2018). Data for *Hydra vulgaris* was excluded from this analysis as Ensembl Compara lacked the *H. vulgaris* orthology relationships required to generate the species tree that this method uses as an input. In **Figure 5b** proteome-wide disorder and accessibility predictions were performed for all species using the approach outlined above (section: structural bioinformatics) with scripts from: https://github.com/BioComputingUP/AlphaFold-disorder (Piovesan *et al*, 2022). Sites with an RSA < 0.2 were considered to be buried and sites with an RSA > 0.4 were considered to be exposed. *Acanthaster planci* was excluded from this analysis as there were too few AF2 structures for a reliable estimate of Y content in buried and exposed regions.

For the analysis of PTM deserts **(Figure 5c-d)**, proteome-wide disorder and RSA prediction was performed as described directly above (Piovesan *et al*, 2022). Each proteome was then simulated 100 times while accounting for the order:disorder content, the amino acid composition of ordered and disordered regions, and the length of each protein in the proteome. A protein was considered to have a Y desert if a contiguous sequence with length 50% or more of the whole protein was missing tyrosine (Szulc *et al*, 2023). Short proteins (length < 150 amino acids) were excluded from the analysis. Gene ontology (GO) enrichment of tyrosine deserts was performed with the clusterProfiler package using all proteins with disorder predictions and length > 150 amino acids as the background set of proteins (Yu *et al*, 2012).

The ‘avoided word’ **(Figure S20)** analysis was performed with AW software (https://github.com/solonas13/aw) that aims to detect DNA or amino acid strings that appear at a significantly lower (or higher) frequency than the chance expectation (Almirantis *et al*, 2019). The AW algorithm was executed to find avoided words (-w 0) of variable length at a score threshold of −3 (-t −3) for the three protein sets described in **Figure S20**.

Testing for pY counter-selection at the site-specific level was performed using Pelican software (Duchemin *et al*, 2023). This software takes as an input a multiple sequence alignment (MSA) and a phylogenetic tree with trait values annotated to each tip and ancestral node. Processed MSAs across all orthologs were taken from the phmmer-derived alignments described in the section above (Evolutionary bioinformatics: human-yeast conservation). Phylogenies for each MSA were generated with FastTree using default parameters (Price *et al*, 2010). We required that the fungal and metazoan sequences were successfully resolved in the phylogenetic tree by excluding trees where the fungal clade contained >2.5% metazoan sequences or *vice versa*. The annotation of binary ancestral traits (‘fungi’ or ‘metazoa’) was performed using the parsimony method within Pelican (Duchemin *et al*, 2023). Pelican was executed with the ‘pelican scan discrete’ command using an amino acid alphabet. Pelican p-values (‘aagtr’) were retrieved for all Y amino acids (pY and non-pY) in the *S. cerevisiae c*opies and then the Benjamini-Hochberg method was applied across all orthologous groups to correct for multiple-hypothesis testing (Benjamini & Hochberg, 1995). For the analysis of these results we exclude as potential confounders any native pY detected in (Lanz *et al*, 2021) or (Leutert *et al*, 2023), and any spurious pY aligning to a native human pY **(Figure 4f).**

### Code and data availability

The mass spectrometry proteomics data (**Table S17**) have been deposited to the ProteomeXchange Consortium via the PRIDE (Perez-Riverol *et al*, 2022) partner repository with the dataset identifier PXD045466 (https://www.ebi.ac.uk/pride/archive/projects/PXD045466).

**Reviewer account details (to access PXD045466 before final submission): Username: Password: J9cNJkOx Log in an access data via “Review Submission” panel on left side**

All R code relating to mass spectrometry data analysis is publicly available under https://gitlab.com/public_villenlab/spurious-phosphorylation. All R code relating to the analysis of protein structure, fitness, and evolution is publicly available under: https://github.com/Landrylab/Bradley_2023_spurious_pY.

## Acknowledgements

We would like to thank members of the Villen and Landry labs for providing feedback on the science. We would like to thank Ricard A. Rodriguez-Mias in the Villen lab for assistance with mass spectrometry. We would like to thank Angel F. Cisneros for assistance with the FoldX analysis. We would also like to thank Louis Duchemin, Philippe Verber, and Bastien Boussau for help using the Pelican software, and Solon P. Pissis for help using the Avoided Word (AW) software. This work was primarily funded by the Human Frontier Science Program research grant RGP34/2018 (to JV and CRL). CRL was also funded by the Canadian Institutes of Health Research Foundation grant number 387697. JV was also supported by NIH grants R35GM119536 and R01AG056359. DB was funded by the EMBO Long-Term Fellowship (LTF) (ALTF 1069-2019). AH was supported by DFF International Postdoctoral Grant (0131-00031B) and EMBO non-stipendiary Postdoctoral Fellowship (ALTF481–2020). RD was funded by the FRQS postdoctoral fellowship. ML was supported by the Swiss National Science Foundation grant P5R5PB_211122. UD was funded by an NSERC graduate fellowship. AC is supported by NIH training grant T32HG000035.

