## Supplementary_figures for "The fitness cost of spurious phosphorylation"

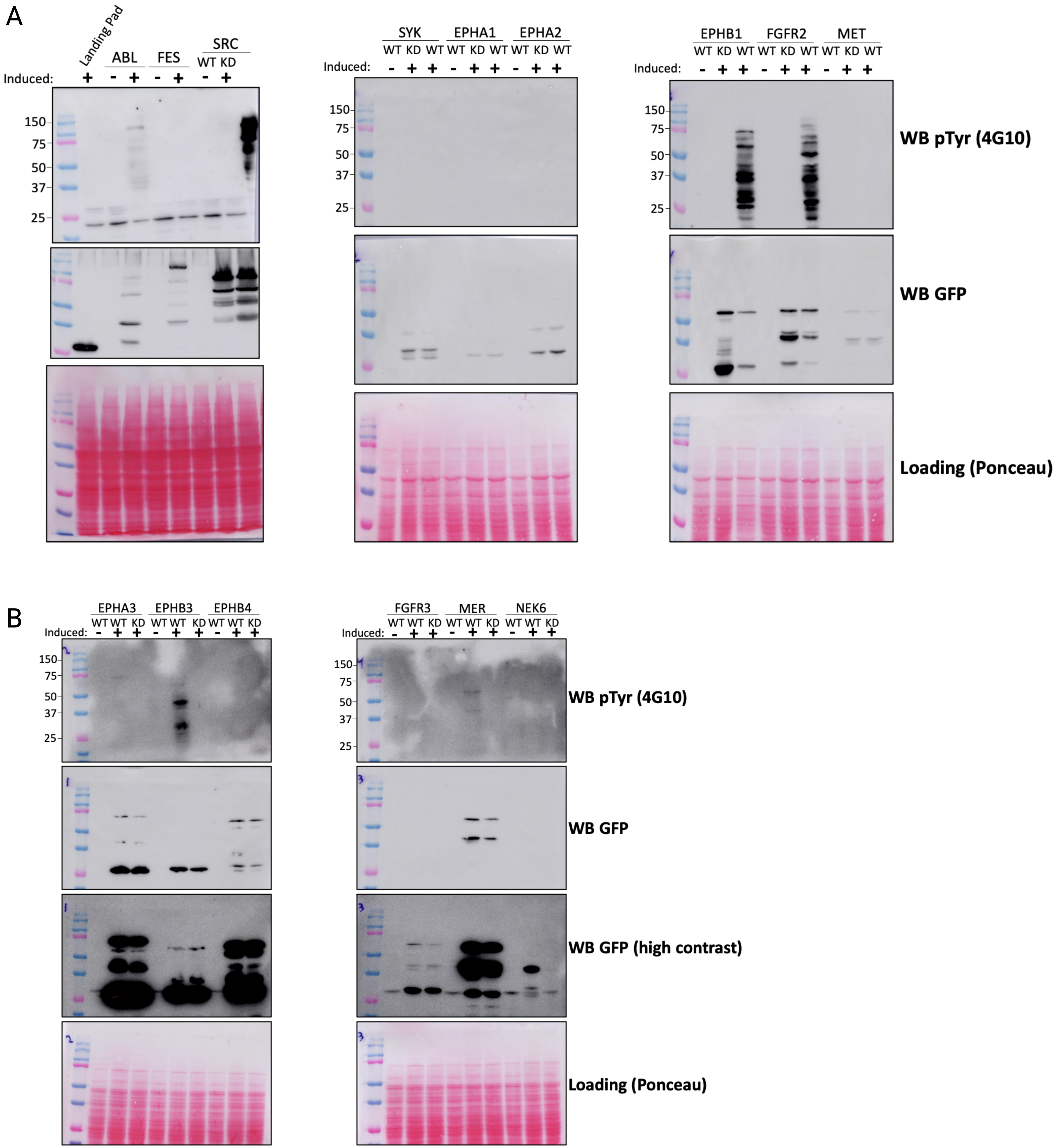
A
Landing Pad
ABL
FES
SRC
WT KD
Induced:
150
75
50
37
25
SYK
EPHA1
EPHA2
WT KD WT WT KD WT WT KD WT
EPHB1
FGFR2
MET
WT KD WT WT KD WT WT KD WT
WB pTyr (4G10)
WB GFP
Loading (Ponceau)
B
EPHA3
EPHB3
EPHB4
WT WT KD WT WT KD WT WT KD
FGFR3
MER
NEK6
WT WT KD WT WT KD WT WT KD
WB pTyr (4G10)
WB GFP
WB GFP (high contrast)
Loading (Ponceau)

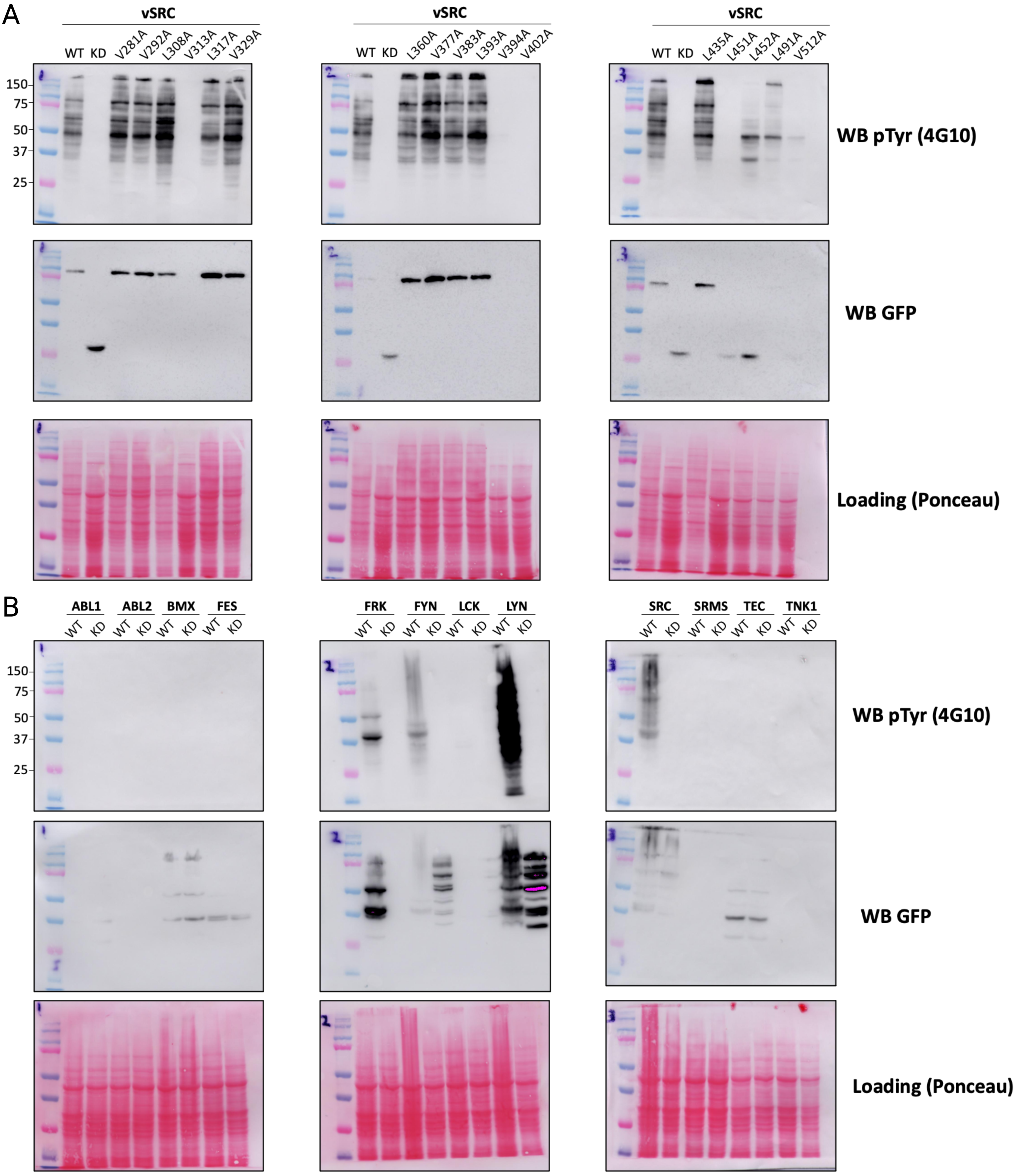
A
vSRC
WT KD V281A V292A L308A V313A L317A V329A
vSRC
WT KD L360A V377A V383A L393A V394A V402A
vSRC
WT KD L435A L451A L452A L491A V512A
150
75
50
37
25
WB pTyr (4G10)
WB GFP
Loading (Ponceau)
B
ABL1 ABL2 BMX FES
WT KD WT KD WT KD WT KD
FRK FYN LCK LYN
WT KD WT KD WT KD WT KD
SRC SRMS TEC TNK1
WT KD WT KD WT KD WT KD
150
75
50
37
25
WB pTyr (4G10)
WB GFP
Loading (Ponceau)

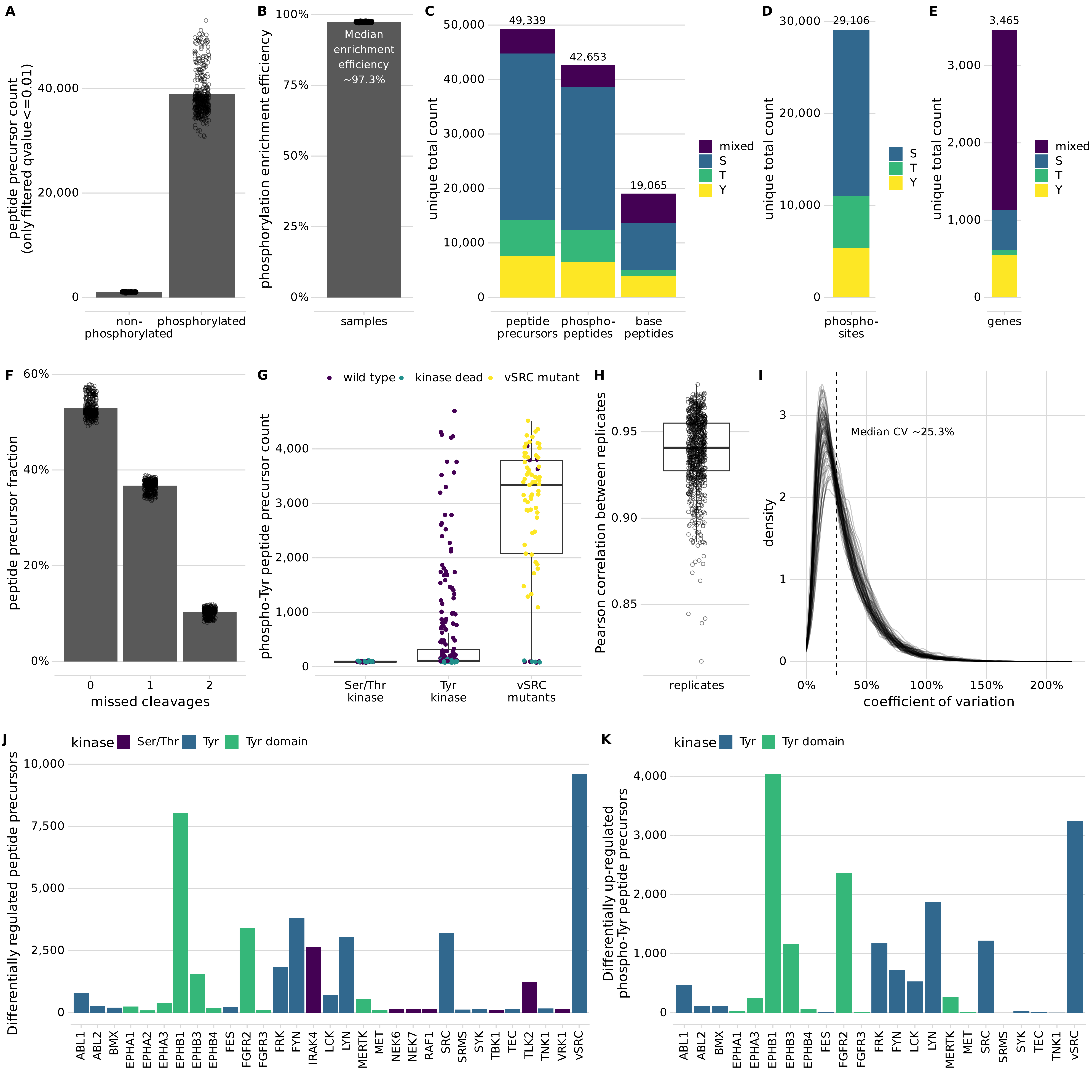
A
peptide precursor count (only filtered qvalue<=0.01)
0
20,000
40,000
non-phosphorylated
phosphorylated
B
phosphorylation enrichment efficiency
0%
25%
50%
75%
100%
Median enrichment efficiency ~97.3%
samples
C
unique total count
0
10,000
20,000
30,000
40,000
50,000
49,339
42,653
19,065
peptide precursors
phospho-peptides
base peptides
mixed
S
T
Y
D
unique total count
0
10,000
20,000
30,000
29,106
phospho-sites
S
T
Y
E
unique total count
0
1,000
2,000
3,000
3,465
genes
mixed
S
T
Y
F
peptide precursor fraction
0%
20%
40%
60%
0
1
2
missed cleavages
G
wild type
kinase dead
vSRC mutant
phospho-Tyr peptide precursor count
0
1,000
2,000
3,000
4,000
Ser/Thr kinase
Tyr kinase
vSRC mutants
H
Pearson correlation between replicates
0.85
0.90
0.95
replicates
I
density
0
1
2
3
Median CV ~25.3%
0%
50%
100%
150%
200%
coefficient of variation
J
kinase
Ser/Thr
Tyr
Tyr domain
Differentially regulated peptide precursors
0
2,500
5,000
7,500
10,000
ABL1
ABL2
BMX
EPHA1
EPHA2
EPHA3
EPHB1
EPHB3
EPHB4
FES
FGFR2
FGFR3
FRK
FYN
IRAK4
LCK
LYN
MERTK
MET
NEK6
NEK7
RAF1
SRC
SRMS
SYK
TBK1
TEC
TLK2
TNK1
VRK1
vSRC
K
kinase
Tyr
Tyr domain
Differentially up-regulated phospho-Tyr peptide precursors
0
1,000
2,000
3,000
4,000
ABL1
ABL2
BMX
EPHA1
EPHA3
EPHB1
EPHB3
EPHB4
FES
FGFR2
FGFR3
FRK
FYN
LCK
LYN
MERTK
MET
SRC
SRMS
SYK
TEC
TNK1
vSRC

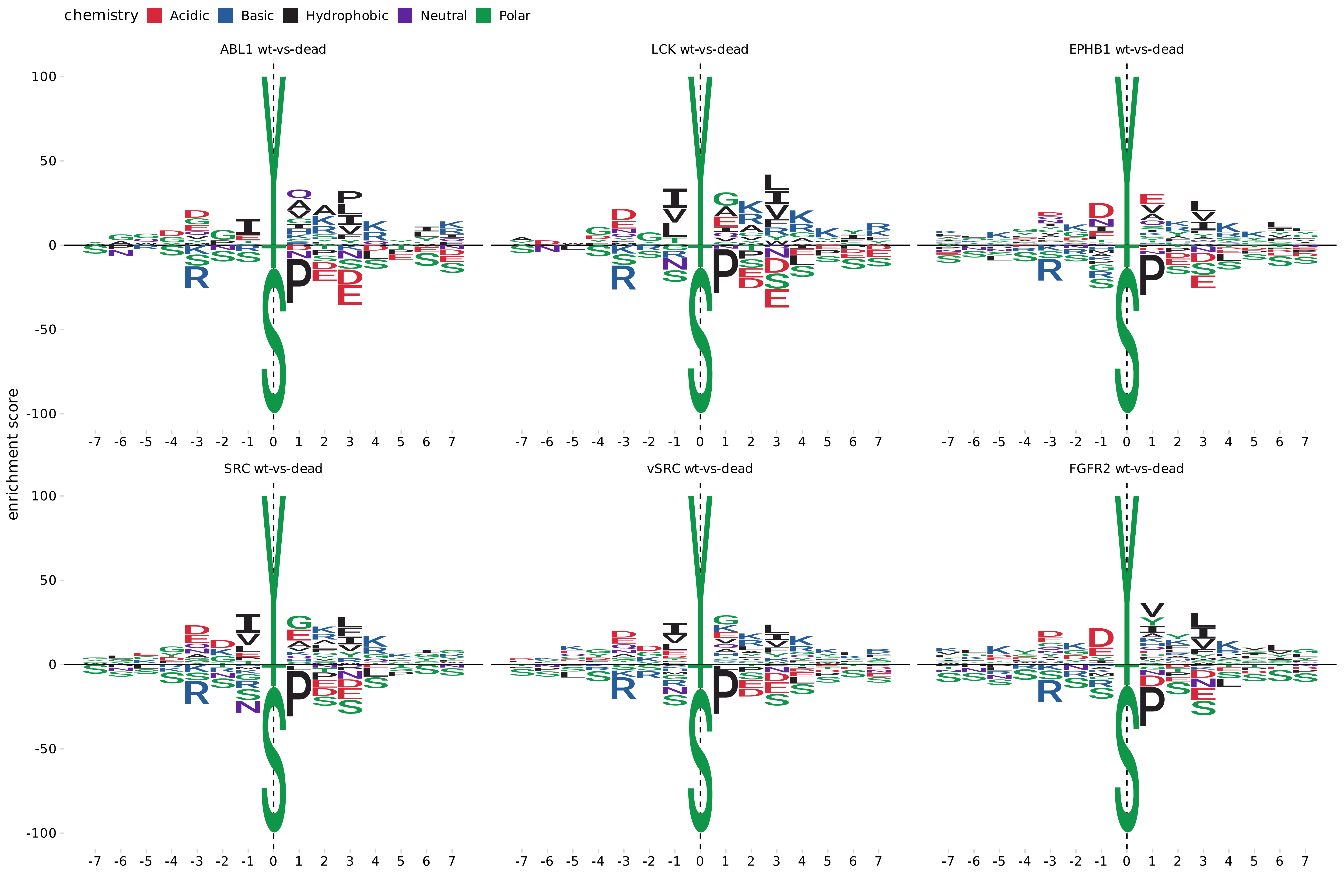

chemistry
Acidic
Basic
Hydrophobic
Neutral
Polar
ABL1 wt-vs-dead
LCK wt-vs-dead
EPHB1 wt-vs-dead
SRC wt-vs-dead
vSRC wt-vs-dead
FGFR2 wt-vs-dead
enrichment score
100
50
0
-50
-100
-7 -6 -5 -4 -3 -2 -1 0 1 2 3 4 5 6 7

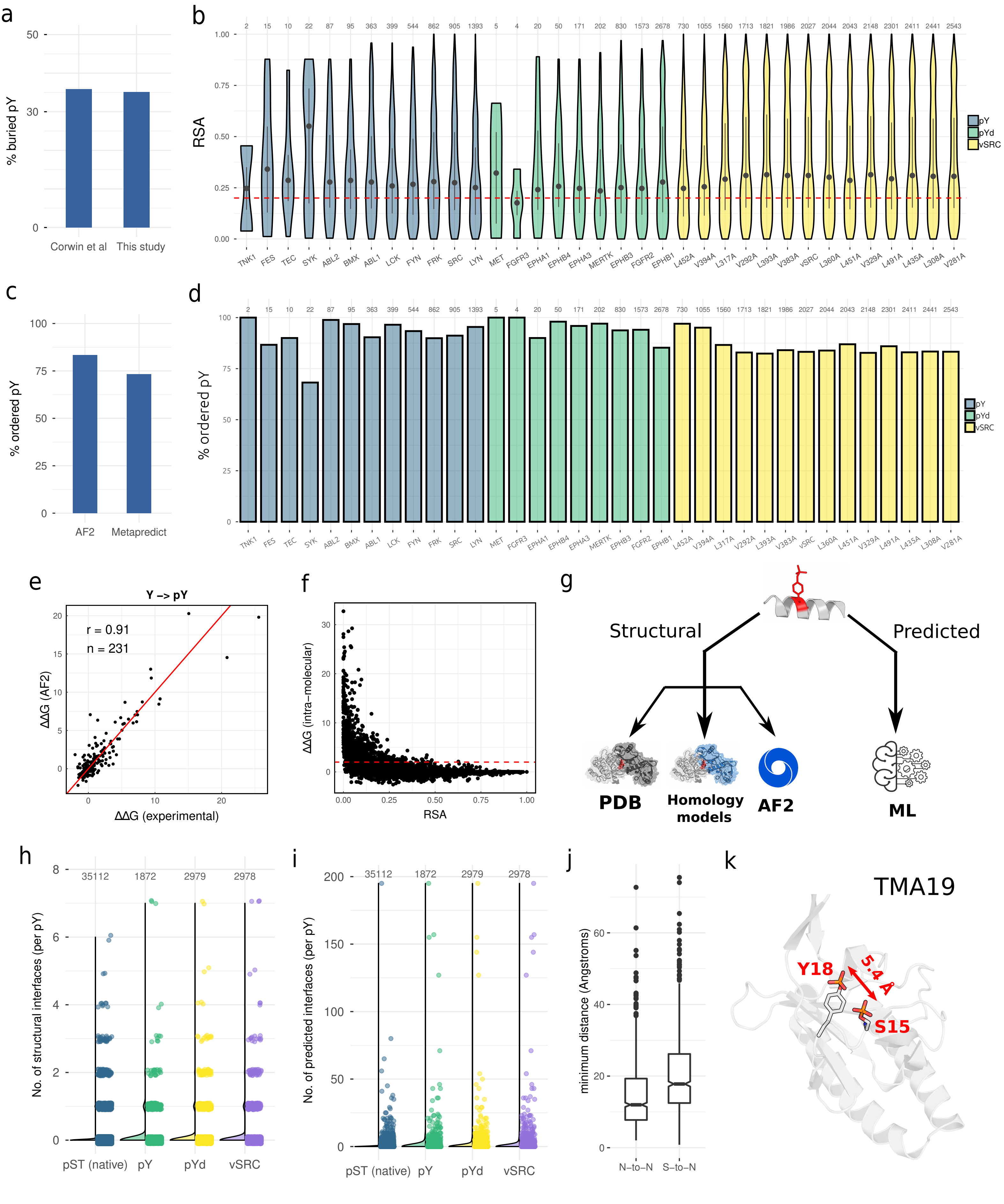

a
% buried pY
Corwin et al
This study
b
RSA
pY
pYd
vSRC
TNK1 FES TEC SYK ABL2 BMX ABL1 LCK FYN FRK SRC LYN MET FGFR3 EPHA1 EPHB4 EPHA3 MERTK EPHB3 FGFR2 EPHB1 L452A V394A L317A V292A L393A V383A vSRC L360A L451A V329A L491A L435A L308A V281A
c
% ordered pY
AF2
Metapredict
d
% ordered pY
e
Y –> pY
r = 0.91
n = 231
ΔΔG (AF2)
ΔΔG (experimental)
f
ΔΔG (intra-molecular)
RSA
g
Structural
Predicted
PDB
Homology models
AF2
ML
h
No. of structural interfaces (per pY)
pST (native)
pY
pYd
vSRC
i
No. of predicted interfaces (per pY)
j
minimum distance (Angstroms)
N–to–N
S–to–N
k
TMA19
Y18
5.4 Å
S15

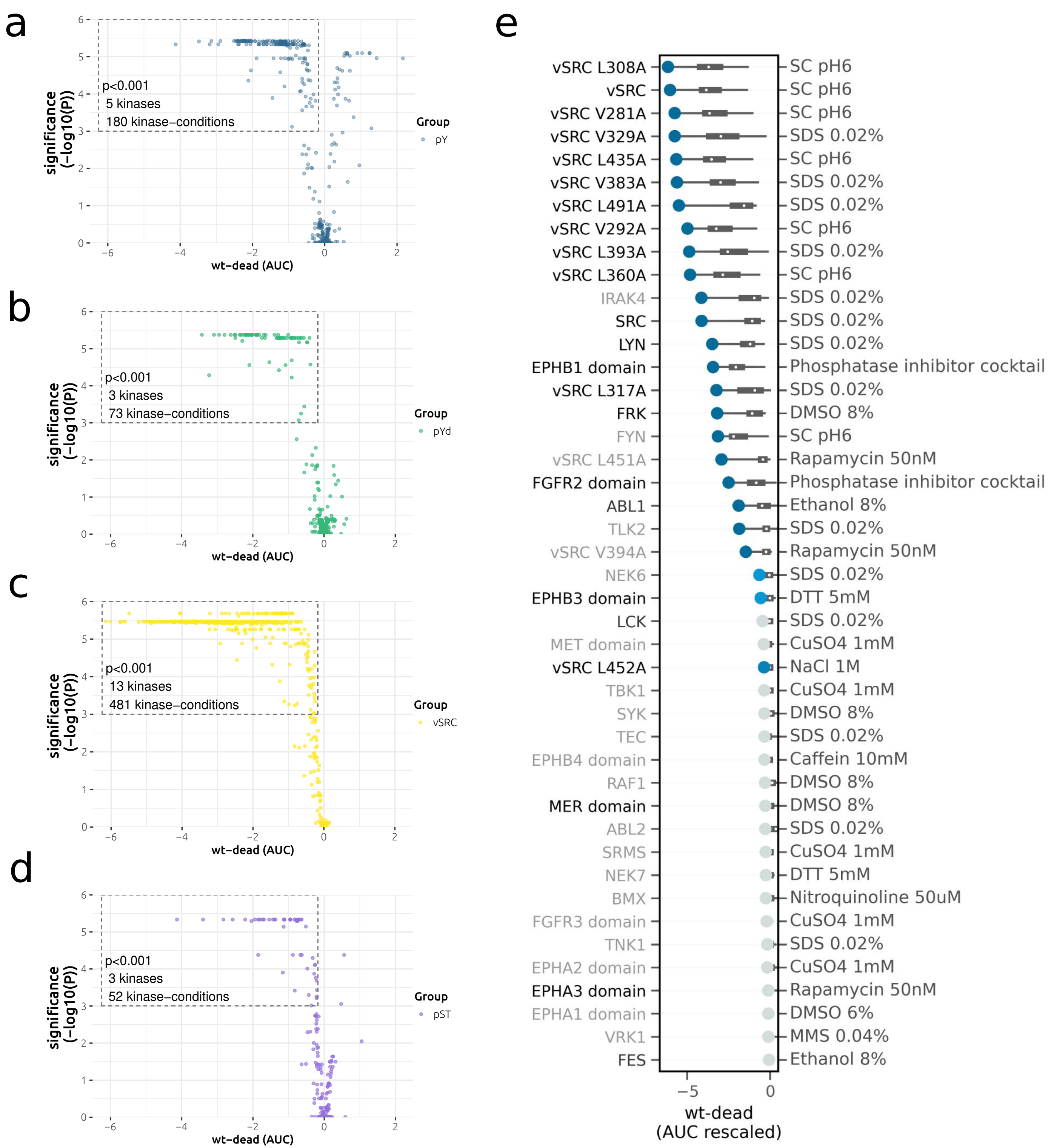

a
significance (−log10(P))
wt−dead (AUC)
p<0.001
5 kinases
180 kinase−conditions
Group
pY
b
significance (−log10(P))
wt−dead (AUC)
p<0.001
3 kinases
73 kinase−conditions
Group
pYd
c
significance (−log10(P))
wt−dead (AUC)
p<0.001
13 kinases
481 kinase−conditions
Group
vSRC
d
significance (−log10(P))
wt−dead (AUC)
p<0.001
3 kinases
52 kinase−conditions
Group
pST
e
vSRC L308A — SC pH6
vSRC — SC pH6
vSRC V281A — SC pH6
vSRC V329A — SDS 0.02%
vSRC L435A — SC pH6
vSRC V383A — SDS 0.02%
vSRC L491A — SDS 0.02%
vSRC V292A — SC pH6
vSRC L393A — SDS 0.02%
vSRC L360A — SC pH6
IRAK4 — SDS 0.02%
SRC — SDS 0.02%
LYN — SDS 0.02%
EPHB1 domain — Phosphatase inhibitor cocktail
vSRC L317A — SDS 0.02%
FRK — DMSO 8%
FYN — SC pH6
vSRC L451A — Rapamycin 50nM
FGFR2 domain — Phosphatase inhibitor cocktail
ABL1 — Ethanol 8%
TLK2 — SDS 0.02%
vSRC V394A — Rapamycin 50nM
NEK6 — SDS 0.02%
EPHB3 domain — DTT 5mM
LCK — SDS 0.02%
MET domain — CuSO4 1mM
vSRC L452A — NaCl 1M
TBK1 — CuSO4 1mM
SYK — DMSO 8%
TEC — SDS 0.02%
EPHB4 domain — Caffein 10mM
RAF1 — DMSO 8%
MER domain — DMSO 8%
ABL2 — SDS 0.02%
SRMS — CuSO4 1mM
NEK7 — DTT 5mM
BMX — Nitroquinoline 50uM
FGFR3 domain — CuSO4 1mM
TNK1 — SDS 0.02%
EPHA2 domain — CuSO4 1mM
EPHA3 domain — Rapamycin 50nM
EPHA1 domain — DMSO 6%
VRK1 — MMS 0.04%
FES — Ethanol 8%
wt-dead (AUC rescaled)

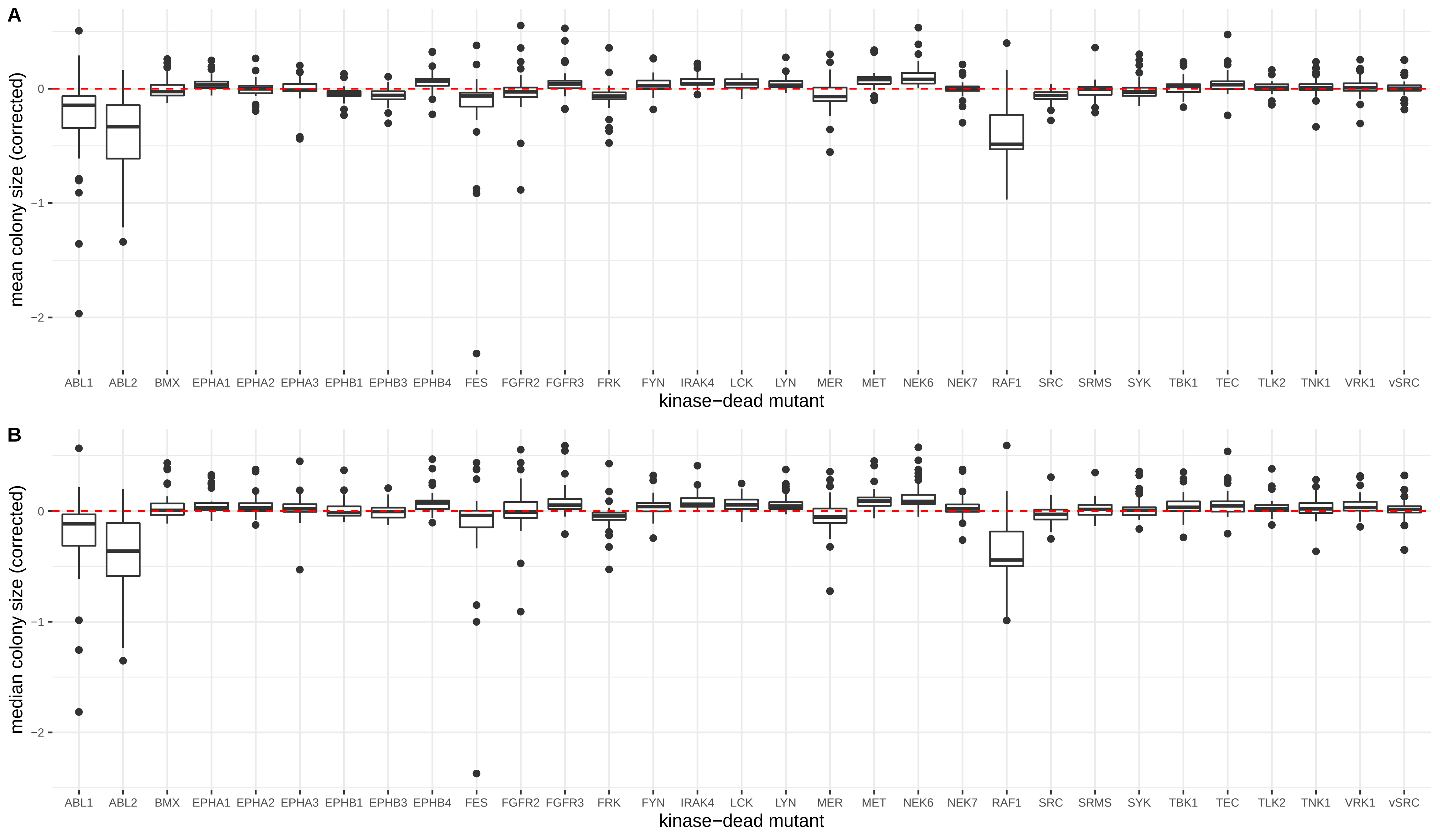
A
mean colony size (corrected)
0
−1
−2
ABL1
ABL2
BMX
EPHA1
EPHA2
EPHA3
EPHB1
EPHB3
EPHB4
FES
FGFR2
FGFR3
FRK
FYN
IRAK4
LCK
LYN
MER
MET
NEK6
NEK7
RAF1
SRC
SRMS
SYK
TBK1
TEC
TLK2
TNK1
VRK1
vSRC
kinase−dead mutant
B
median colony size (corrected)
0
−1
−2
ABL1
ABL2
BMX
EPHA1
EPHA2
EPHA3
EPHB1
EPHB3
EPHB4
FES
FGFR2
FGFR3
FRK
FYN
IRAK4
LCK
LYN
MER
MET
NEK6
NEK7
RAF1
SRC
SRMS
SYK
TBK1
TEC
TLK2
TNK1
VRK1
vSRC
kinase−dead mutant

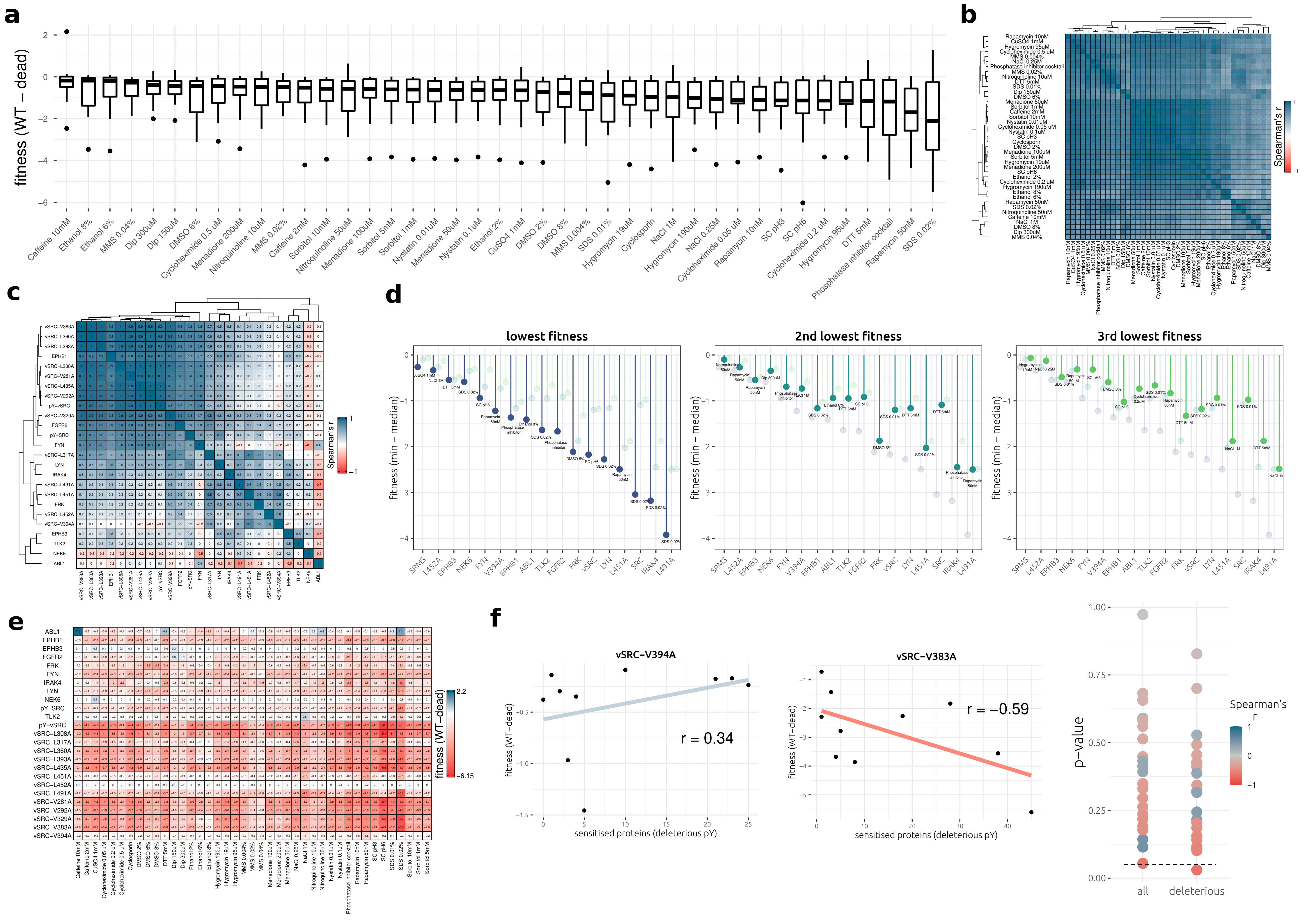
a
fitness (WT – dead)
2
0
−2
−4
−6
Caffeine 10mM
Ethanol 8%
Ethanol 6%
MMS 0.04%
Dip 300uM
Dip 150uM
DMSO 6%
Cycloheximide 0.5 uM
Menadione 200uM
Nitroquinoline 10uM
MMS 0.02%
Caffeine 2mM
Sorbitol 10mM
Nitroquinoline 50uM
Menadione 100uM
Sorbitol 5mM
Sorbitol 1mM
Nystatin 0.01uM
Menadione 50uM
Nystatin 0.1uM
Ethanol 2%
CuSO4 1mM
DMSO 2%
DMSO 8%
MMS 0.004%
SDS 0.01%
Hygromycin 19uM
Cyclosporin
NaCl 1M
Hygromycin 190uM
NaCl 0.25M
Cycloheximide 0.05 uM
Rapamycin 10nM
SC pH3
SC pH6
Cycloheximide 0.2 uM
Hygromycin 95uM
DTT 5mM
Phosphatase inhibitor cocktail
Rapamycin 50nM
SDS 0.02%
b
Spearman's r
1
−1
c
Spearman's r
1
−1
vSRC−V383A
vSRC−L360A
vSRC−L393A
EPHB1
vSRC−L308A
vSRC−V281A
vSRC−L435A
vSRC−V292A
pY−vSRC
vSRC−V329A
FGFR2
pY−SRC
FYN
vSRC−L317A
LYN
IRAK4
vSRC−L491A
vSRC−L451A
FRK
vSRC−L452A
vSRC−V394A
EPHB3
TLK2
NEK6
ABL1
d
lowest fitness
2nd lowest fitness
3rd lowest fitness
fitness (min – median)
0
−1
−2
−3
−4
SRMS
L452A
EPHB3
NEK6
FYN
V394A
EPHB1
ABL1
TLK2
FGFR2
FRK
vSRC
LYN
L451A
SRC
IRAK4
L491A
e
fitness (WT−dead)
2.2
−6.15
ABL1
EPHB1
EPHB3
FGFR2
FRK
FYN
IRAK4
LYN
NEK6
pY−SRC
TLK2
pY−vSRC
vSRC−L308A
vSRC−L317A
vSRC−L360A
vSRC−L393A
vSRC−L435A
vSRC−L451A
vSRC−L452A
vSRC−L491A
vSRC−V281A
vSRC−V292A
vSRC−V329A
vSRC−V383A
vSRC−V394A
f
vSRC−V394A
r = 0.34
fitness (WT−dead)
sensitised proteins (deleterious pY)
vSRC−V383A
r = −0.59
fitness (WT−dead)
sensitised proteins (deleterious pY)
p−value
1.00
0.75
0.50
0.25
0.00
all
deleterious
Spearman's r
1
0
−1

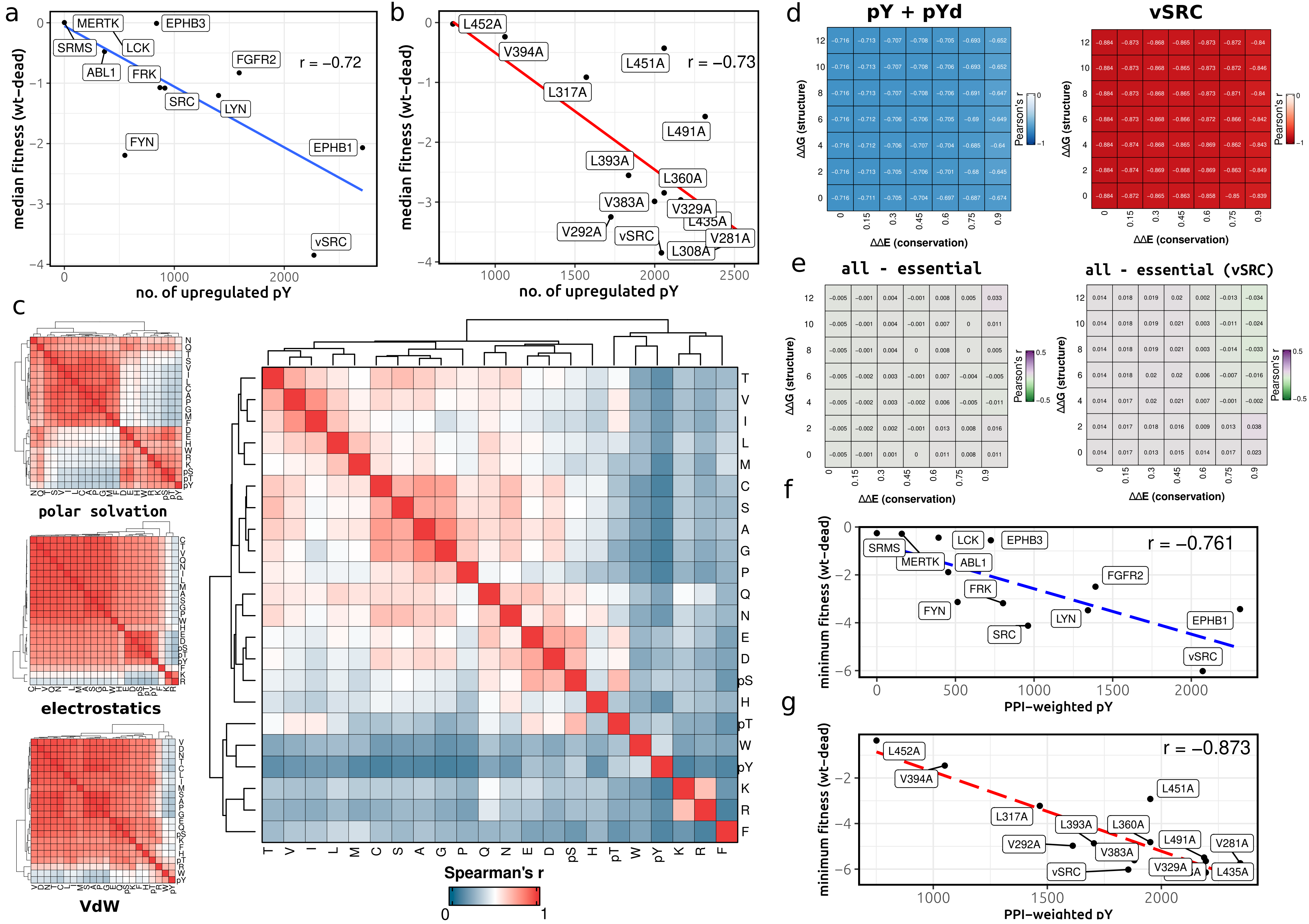
a
MERTK
EPHB3
SRMS
LCK
ABL1
FRK
FGFR2
r = −0.72
SRC
LYN
FYN
EPHB1
vSRC
median fitness (wt−dead)
no. of upregulated pY
b
L452A
V394A
L451A
r = −0.73
L317A
L491A
L393A
L360A
V383A
V329A
L435A
V292A
vSRC
V281A
L308A
median fitness (wt−dead)
no. of upregulated pY
c
polar solvation
electrostatics
VdW
Spearman's r
d
pY + pYd
vSRC
ΔΔG (structure)
ΔΔE (conservation)
Pearson's r
e
all - essential
all - essential (vSRC)
ΔΔG (structure)
ΔΔE (conservation)
Pearson's r
f
SRMS
MERTK
LCK
EPHB3
ABL1
FGFR2
FRK
FYN
LYN
EPHB1
SRC
vSRC
r = −0.761
minimum fitness (wt−dead)
PPI−weighted pY
g
L452A
V394A
L451A
L317A
L393A
L360A
L491A
V292A
V383A
V281A
vSRC
V329A
L435A
r = −0.873
minimum fitness (wt−dead)
PPI−weighted pY

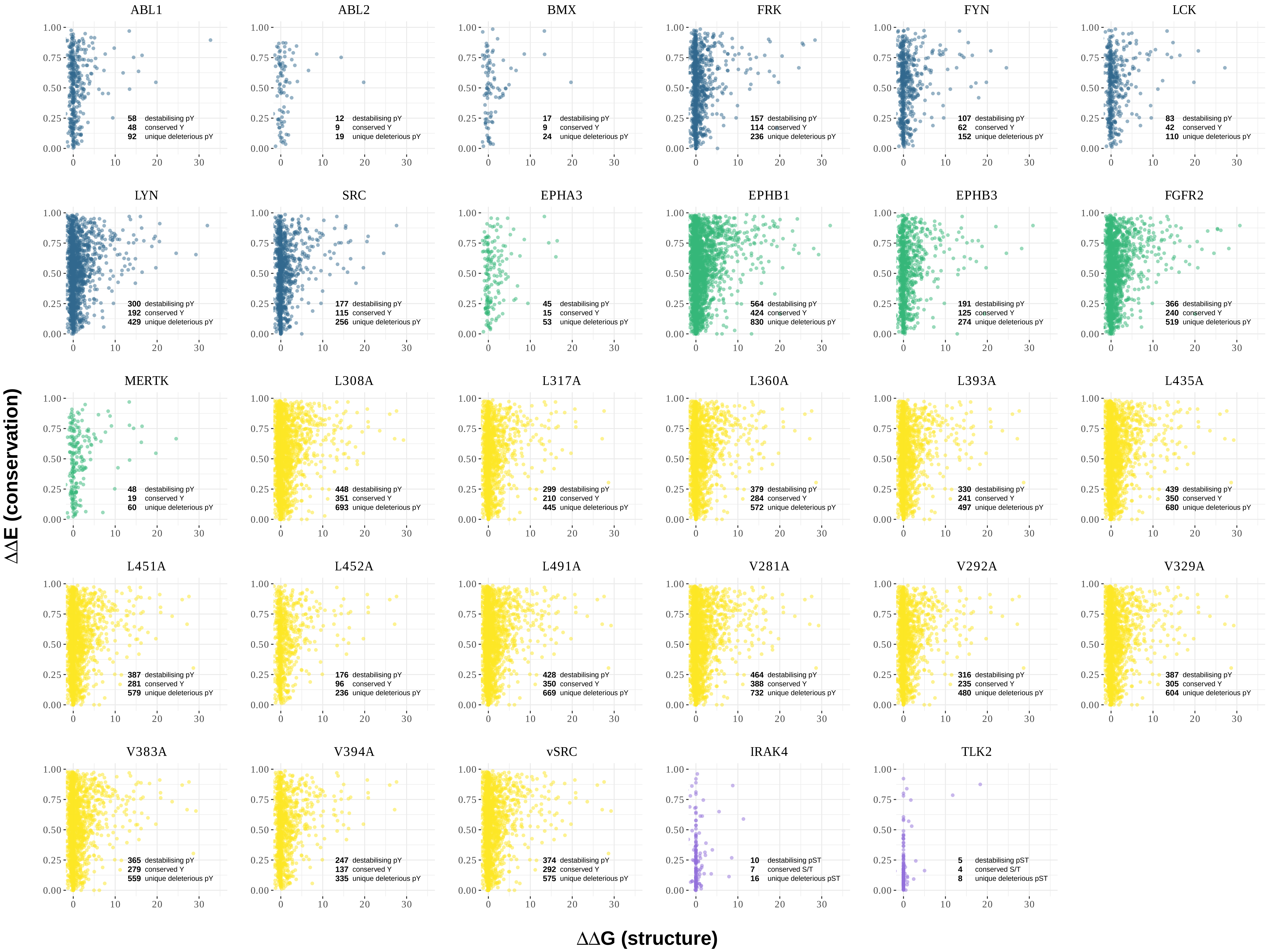

ΔΔE (conservation)
ΔΔG (structure)
ABL1
58 destabilising pY
48 conserved Y
92 unique deleterious pY
ABL2
12 destabilising pY
9 conserved Y
19 unique deleterious pY
BMX
17 destabilising pY
9 conserved Y
24 unique deleterious pY
FRK
157 destabilising pY
114 conserved Y
236 unique deleterious pY
FYN
107 destabilising pY
62 conserved Y
152 unique deleterious pY
LCK
83 destabilising pY
42 conserved Y
110 unique deleterious pY
LYN
300 destabilising pY
192 conserved Y
429 unique deleterious pY
SRC
177 destabilising pY
115 conserved Y
256 unique deleterious pY
EPHA3
45 destabilising pY
15 conserved Y
53 unique deleterious pY
EPHB1
564 destabilising pY
424 conserved Y
830 unique deleterious pY
EPHB3
191 destabilising pY
125 conserved Y
274 unique deleterious pY
FGFR2
366 destabilising pY
240 conserved Y
519 unique deleterious pY
MERTK
48 destabilising pY
19 conserved Y
60 unique deleterious pY
L308A
448 destabilising pY
351 conserved Y
693 unique deleterious pY
L317A
299 destabilising pY
210 conserved Y
445 unique deleterious pY
L360A
379 destabilising pY
284 conserved Y
572 unique deleterious pY
L393A
330 destabilising pY
241 conserved Y
497 unique deleterious pY
L435A
439 destabilising pY
350 conserved Y
680 unique deleterious pY
L451A
387 destabilising pY
281 conserved Y
579 unique deleterious pY
L452A
176 destabilising pY
96 conserved Y
236 unique deleterious pY
L491A
428 destabilising pY
350 conserved Y
669 unique deleterious pY
V281A
464 destabilising pY
388 conserved Y
732 unique deleterious pY
V292A
316 destabilising pY
235 conserved Y
480 unique deleterious pY
V329A
387 destabilising pY
305 conserved Y
604 unique deleterious pY
V383A
365 destabilising pY
279 conserved Y
559 unique deleterious pY
V394A
247 destabilising pY
137 conserved Y
335 unique deleterious pY
vSRC
374 destabilising pY
292 conserved Y
575 unique deleterious pY
IRAK4
10 destabilising pST
7 conserved S/T
16 unique deleterious pST
TLK2
5 destabilising pST
4 conserved S/T
8 unique deleterious pST

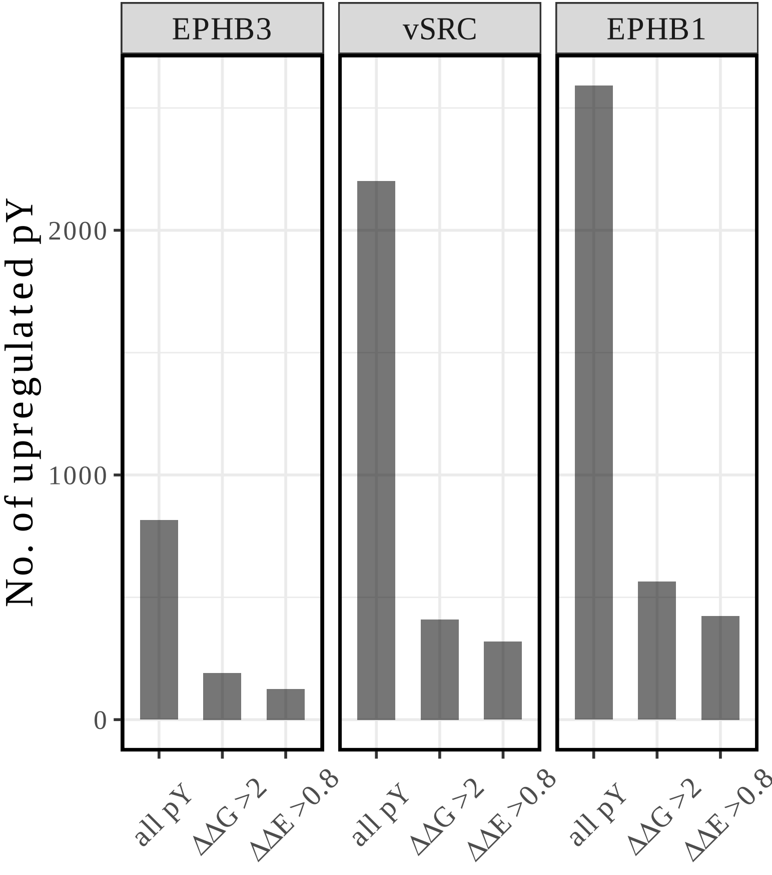

EPHB3
vSRC
EPHB1
No. of upregulated pY
2000
1000
0
all pY
ΔΔG >2
ΔΔE >0.8
all pY
ΔΔG >2
ΔΔE >0.8
all pY
ΔΔG >2
ΔΔE >0.8

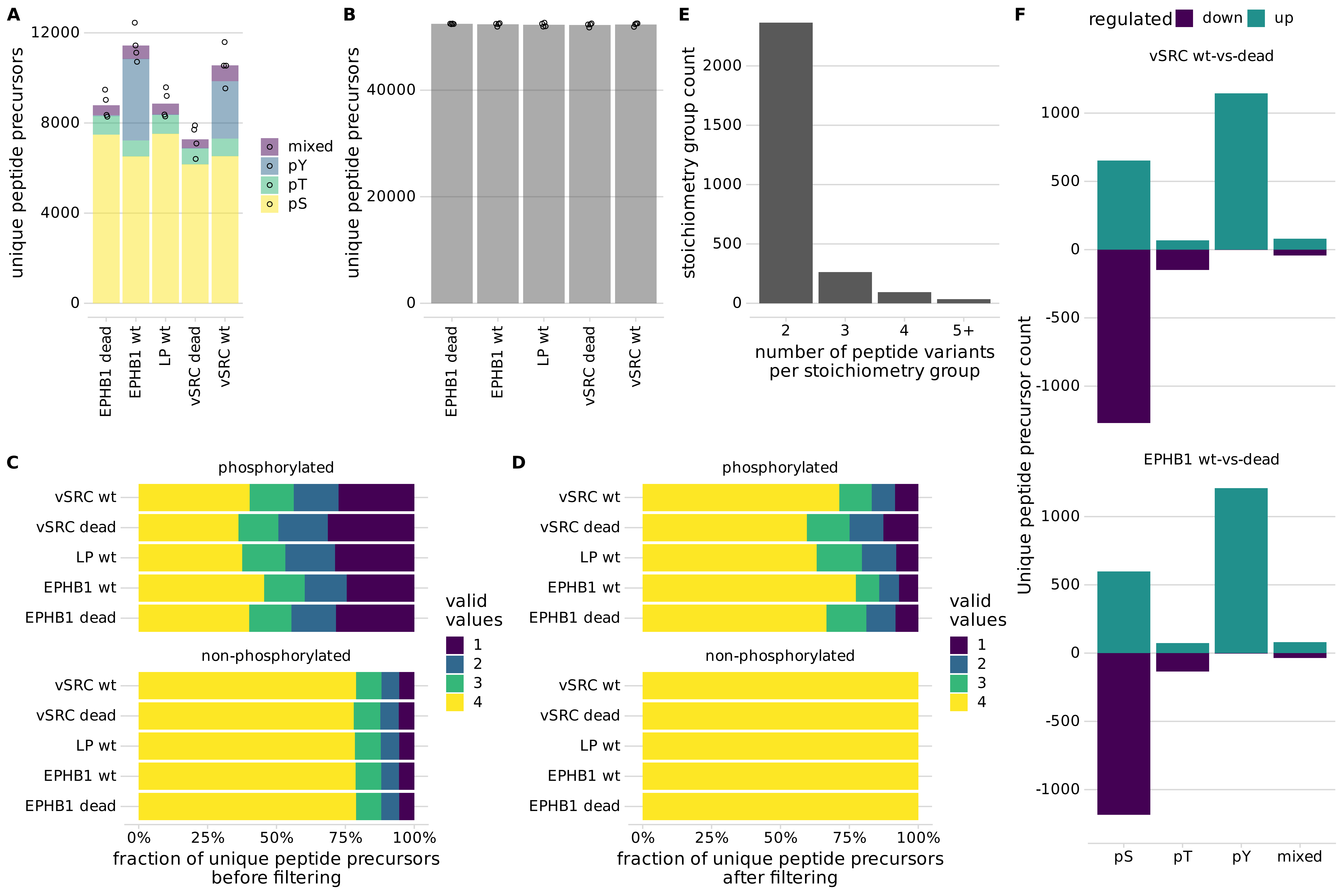

A
unique peptide precursors
12000
8000
4000
0
mixed
pY
pT
pS
EPHB1 dead
EPHB1 wt
LP wt
vSRC dead
vSRC wt
B
unique peptide precursors
40000
20000
0
EPHB1 dead
EPHB1 wt
LP wt
vSRC dead
vSRC wt
E
stoichiometry group count
2000
1500
1000
500
0
2
3
4
5+
number of peptide variants per stoichiometry group
F
regulated
down
up
vSRC wt-vs-dead
EPHB1 wt-vs-dead
Unique peptide precursor count
1000
500
0
-500
-1000
pS
pT
pY
mixed
C
phosphorylated
non-phosphorylated
vSRC wt
vSRC dead
LP wt
EPHB1 wt
EPHB1 dead
valid values
1
2
3
4
0%
25%
50%
75%
100%
fraction of unique peptide precursors before filtering
D
phosphorylated
non-phosphorylated
fraction of unique peptide precursors after filtering

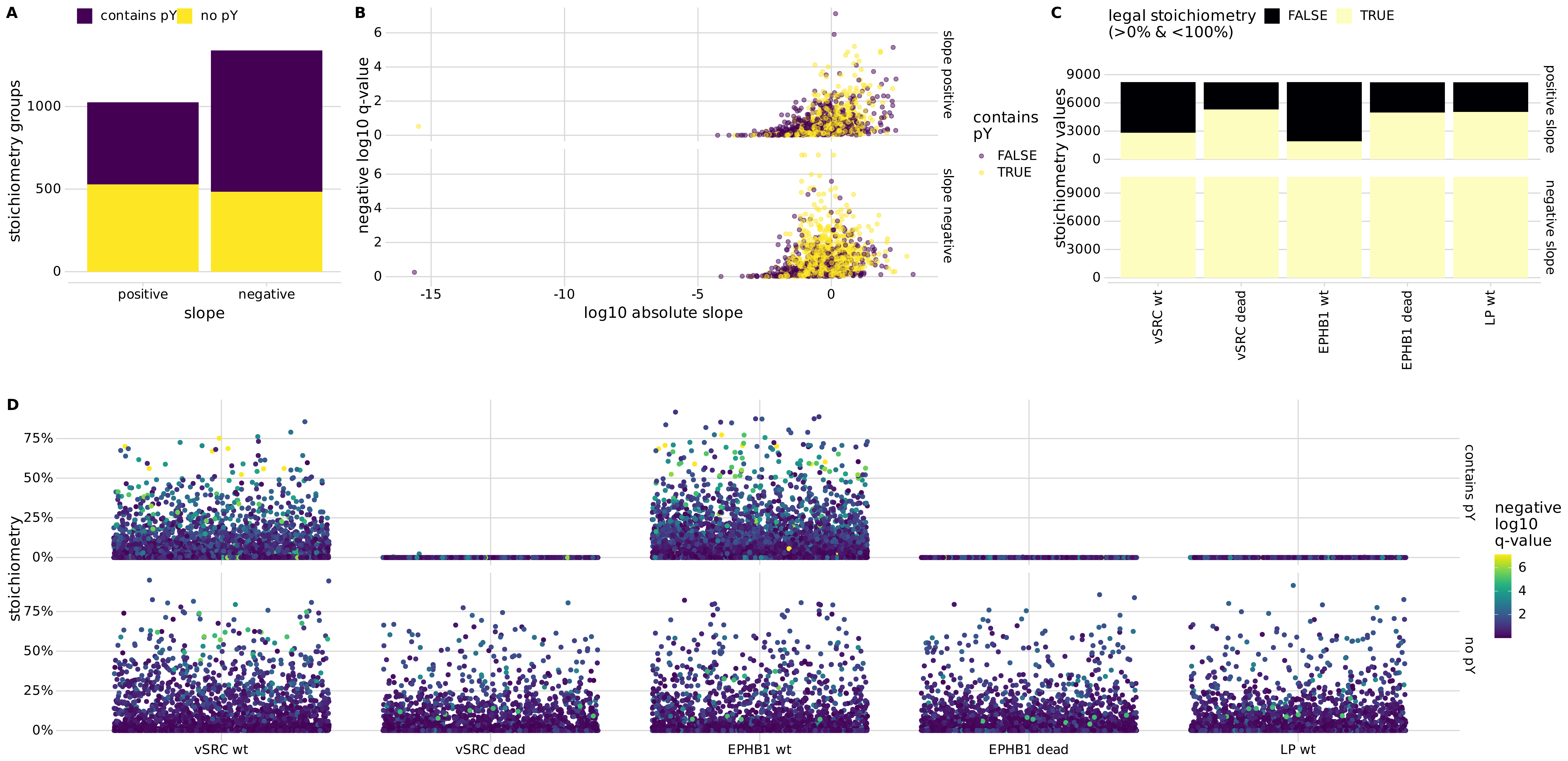
A
contains pY
no pY
stoichiometry groups
1000
500
0
positive
negative
slope
B
negative log10 q-value
6
4
2
0
-15
-10
-5
0
log10 absolute slope
slope positive
slope negative
contains pY
FALSE
TRUE
C
legal stoichiometry (>0% & <100%)
FALSE
TRUE
stoichiometry values
9000
6000
3000
0
positive slope
negative slope
vSRC wt
vSRC dead
EPHB1 wt
EPHB1 dead
LP wt
D
stoichiometry
75%
50%
25%
0%
contains pY
no pY
negative log10 q-value
6
4
2
vSRC wt
vSRC dead
EPHB1 wt
EPHB1 dead
LP wt

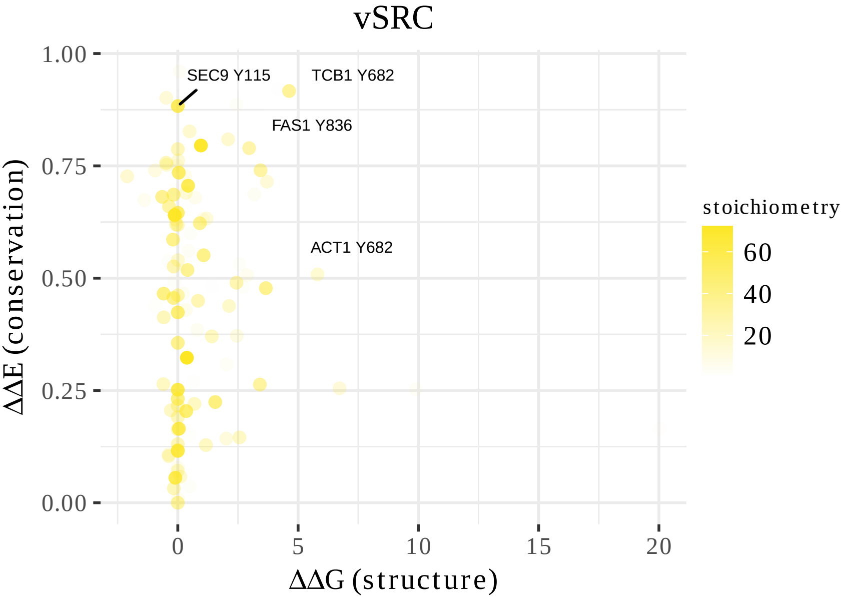

vSRC
SEC9 Y115
TCB1 Y682
FAS1 Y836
ACT1 Y682
1.00
0.75
0.50
0.25
0.00
ΔΔE (conservation)
0
5
10
15
20
ΔΔG (structure)
stoichiometry
60
40
20

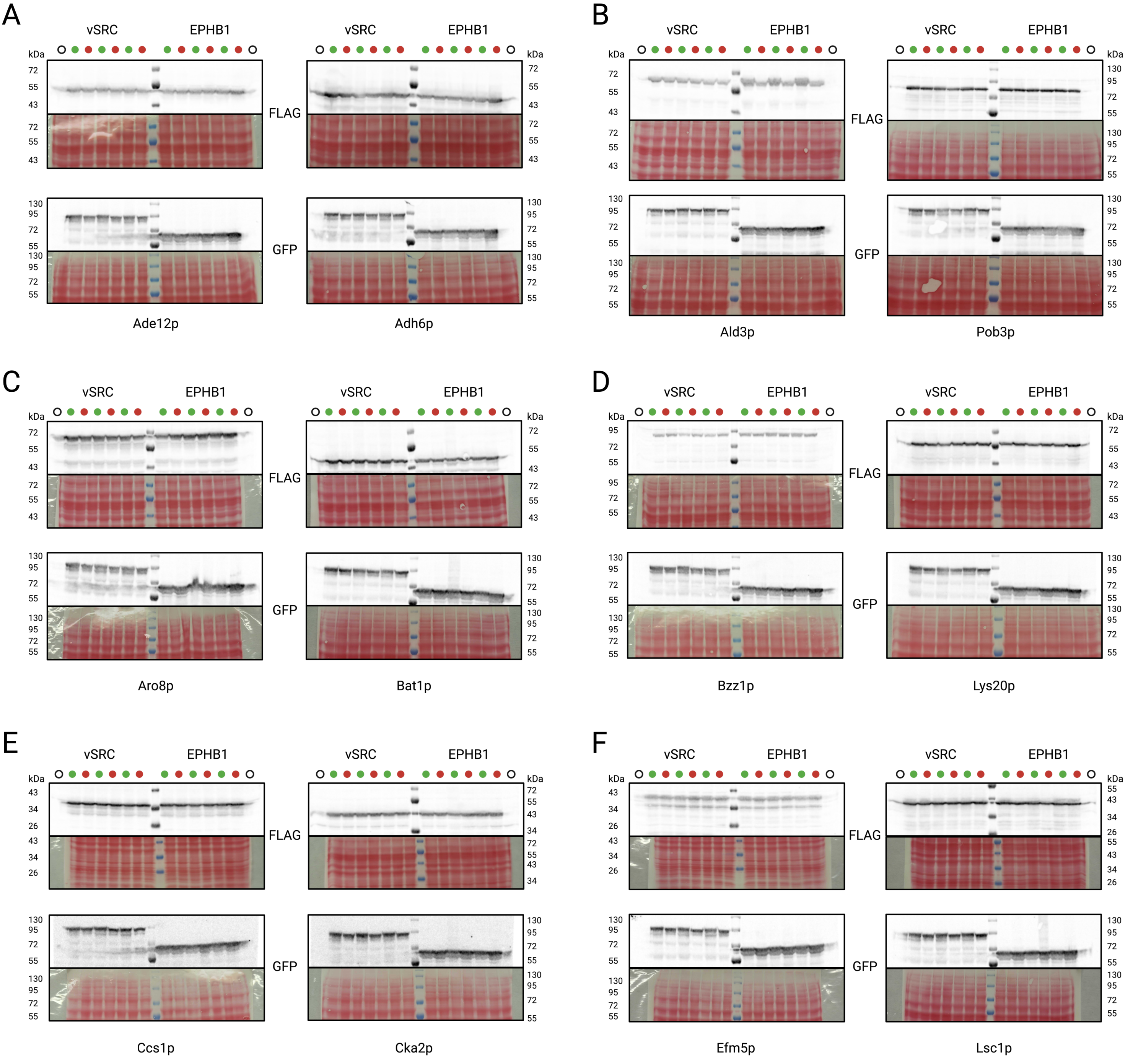
A
vSRC
EPHB1
kDa
FLAG
GFP
Ade12p
Adh6p
B
Ald3p
Pob3p
C
Aro8p
Bat1p
D
Bzz1p
Lys20p
E
Ccs1p
Cka2p
F
Efm5p
Lsc1p

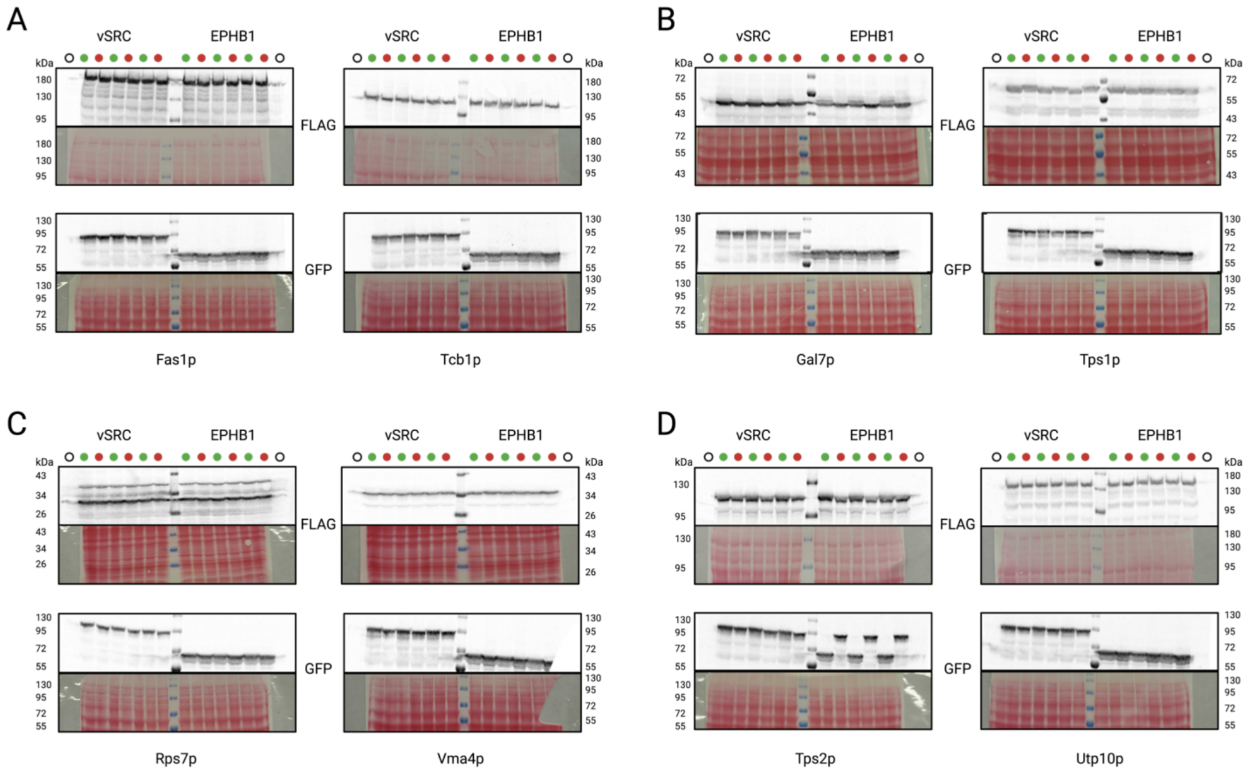

A
B
C
D
vSRC
EPHB1
kDa
FLAG
GFP
Fas1p
Tcb1p
Gal7p
Tps1p
Rps7p
Vma4p
Tps2p
Utp10p

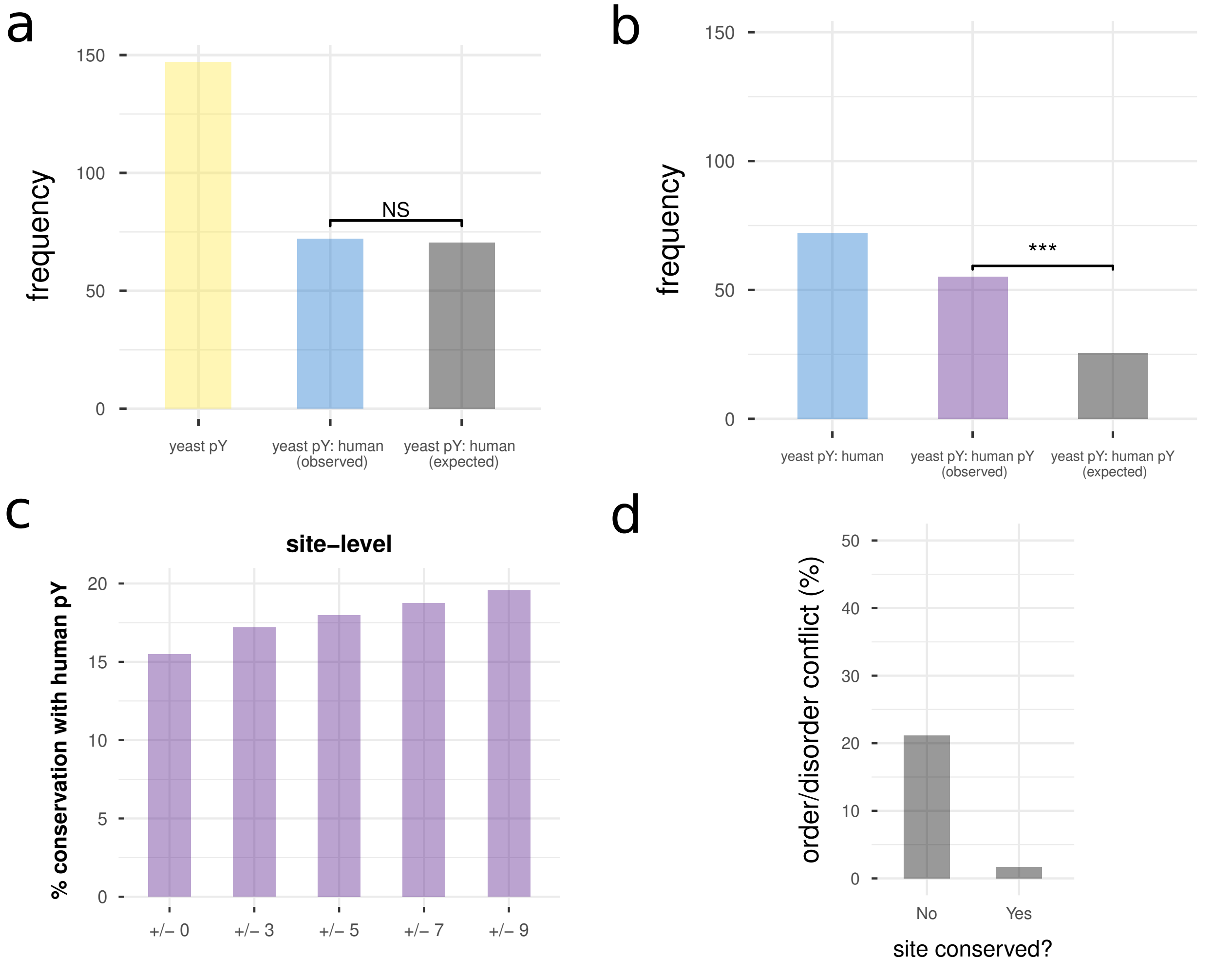
a
frequency
150
100
50
0
NS
yeast pY
yeast pY: human (observed)
yeast pY: human (expected)
b
frequency
150
100
50
0
***
yeast pY: human
yeast pY: human pY (observed)
yeast pY: human pY (expected)
c
site−level
% conservation with human pY
20
15
10
5
0
+/− 0
+/− 3
+/− 5
+/− 7
+/− 9
d
order/disorder conflict (%)
50
40
30
20
10
0
No
Yes
site conserved?

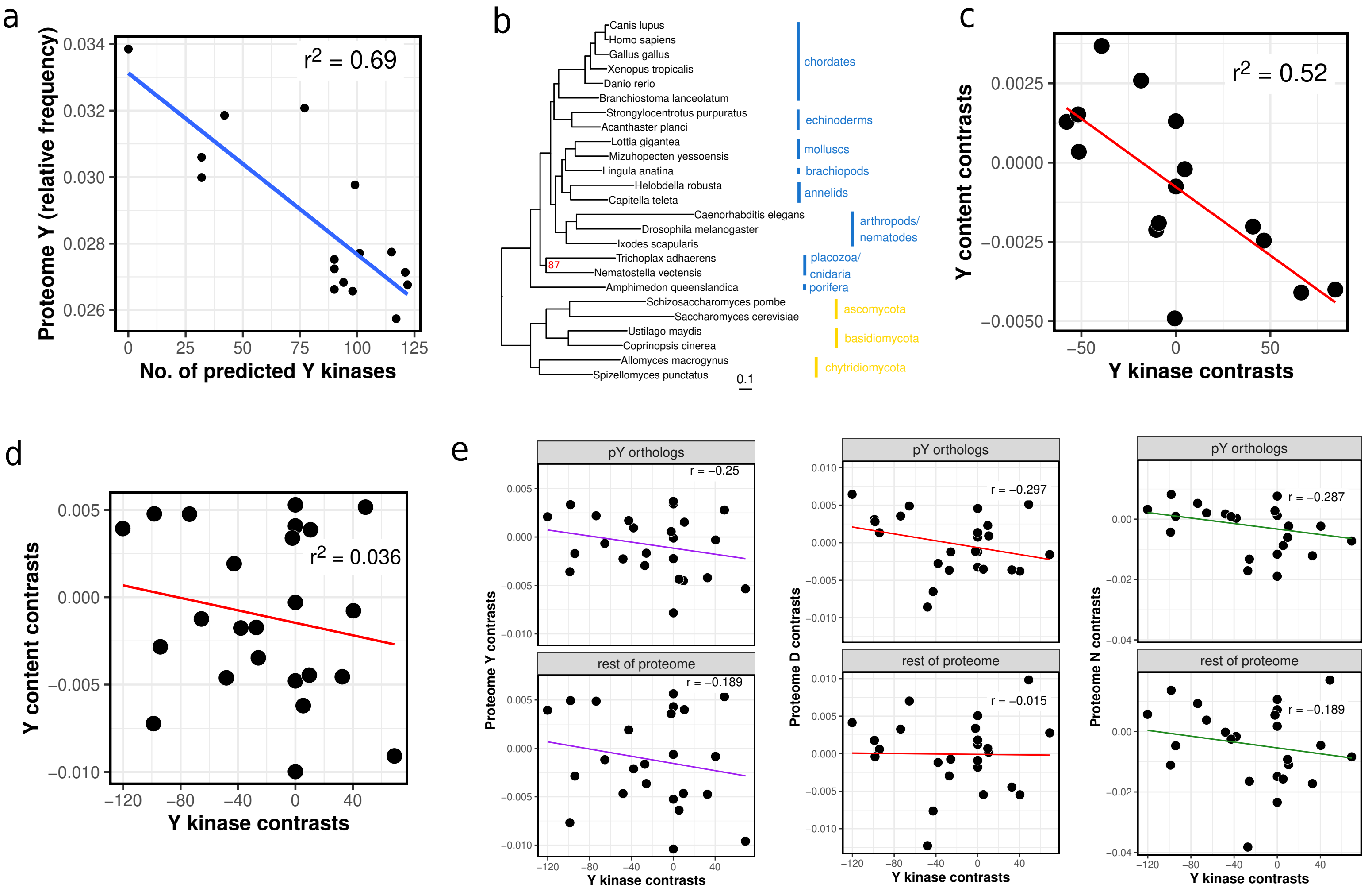

a
r2 = 0.69
Proteome Y (relative frequency)
No. of predicted Y kinases
b
Canis lupus
Homo sapiens
Gallus gallus
Xenopus tropicalis
Danio rerio
Branchiostoma lanceolatum
Strongylocentrotus purpuratus
Acanthaster planci
Lottia gigantea
Mizuhopecten yessoensis
Lingula anatina
Helobdella robusta
Capitella teleta
Caenorhabditis elegans
Drosophila melanogaster
Ixodes scapularis
Trichoplax adhaerens
Nematostella vectensis
Amphimedon queenslandica
Schizosaccharomyces pombe
Saccharomyces cerevisiae
Ustilago maydis
Coprinopsis cinerea
Allomyces macrogynus
Spizellomyces punctatus
87
0.1
chordates
echinoderms
molluscs
brachiopods
annelids
arthropods/ nematodes
placozoa/ cnidaria
porifera
ascomycota
basidiomycota
chytridiomycota
c
r2 = 0.52
Y content contrasts
Y kinase contrasts
d
r2 = 0.036
Y content contrasts
Y kinase contrasts
e
pY orthologs
rest of proteome
r = −0.25
r = −0.189
Proteome Y contrasts
r = −0.297
r = −0.015
Proteome D contrasts
r = −0.287
r = −0.189
Proteome N contrasts
Y kinase contrasts

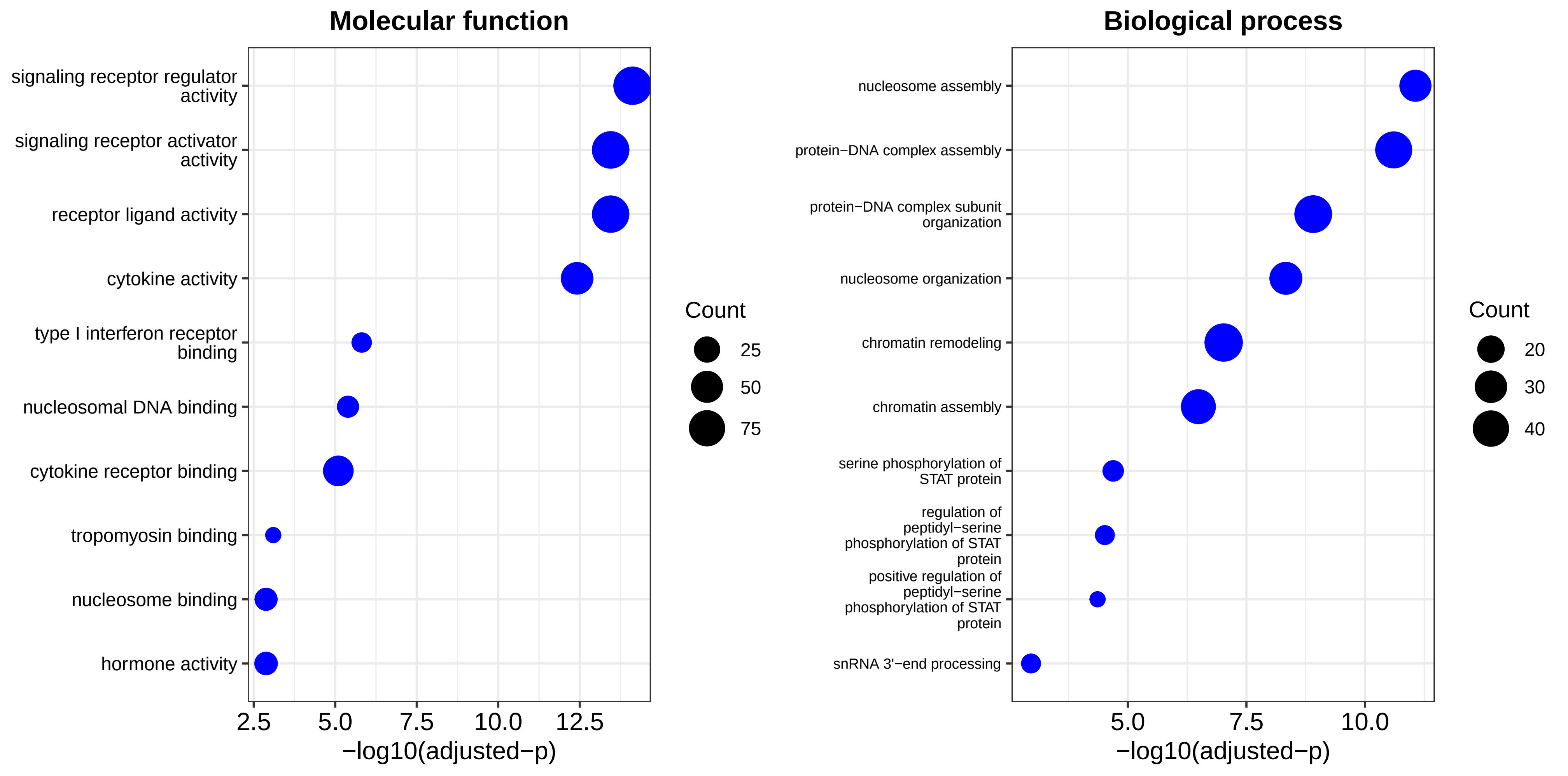

Molecular function
signaling receptor regulator activity
signaling receptor activator activity
receptor ligand activity
cytokine activity
type I interferon receptor binding
nucleosomal DNA binding
cytokine receptor binding
tropomyosin binding
nucleosome binding
hormone activity
2.5
5.0
7.5
10.0
12.5
−log10(adjusted−p)
Count
25
50
75
Biological process
nucleosome assembly
protein−DNA complex assembly
protein−DNA complex subunit organization
nucleosome organization
chromatin remodeling
chromatin assembly
serine phosphorylation of STAT protein
regulation of peptidyl−serine phosphorylation of STAT protein
positive regulation of peptidyl−serine phosphorylation of STAT protein
snRNA 3'−end processing
5.0
7.5
10.0
−log10(adjusted−p)
Count
20
30
40

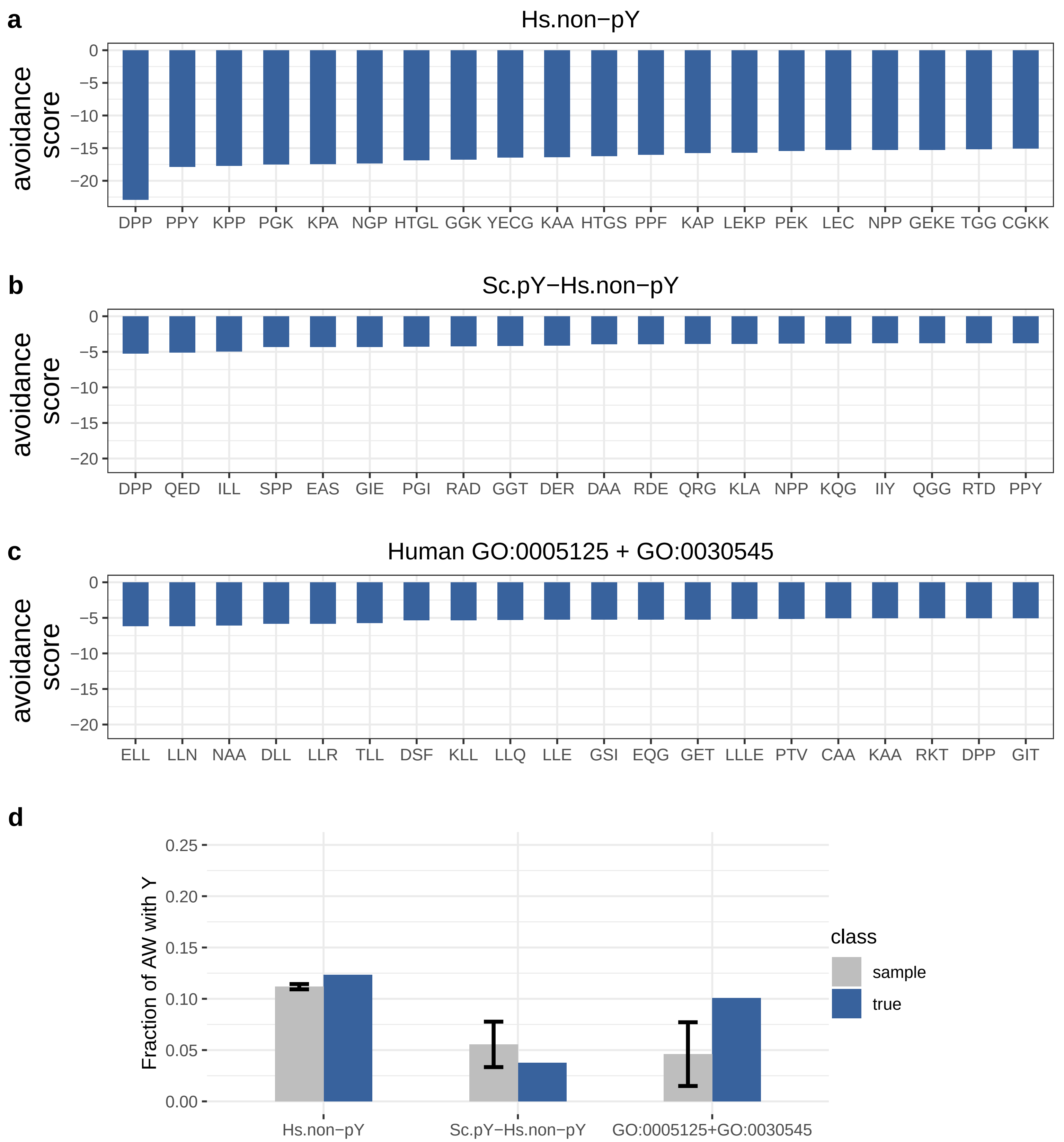
a
Hs.non−pY
avoidance score
0
−5
−10
−15
−20
DPP PPY KPP PGK KPA NGP HTGL GGK YECG KAA HTGS PPF KAP LEKP PEK LEC NPP GEKE TGG CGKK
b
Sc.pY−Hs.non−pY
avoidance score
DPP QED ILL SPP EAS GIE PGI RAD GGT DER DAA RDE QRG KLA NPP KQG IIY QGG RTD PPY
c
Human GO:0005125 + GO:0030545
avoidance score
ELL LLN NAA DLL LLR TLL DSF KLL LLQ LLE GSI EQG GET LLLE PTV CAA KAA RKT DPP GIT
d
Fraction of AW with Y
0.25
0.20
0.15
0.10
0.05
0.00
class
sample
true
Hs.non−pY
Sc.pY−Hs.non−pY
GO:0005125+GO:0030545

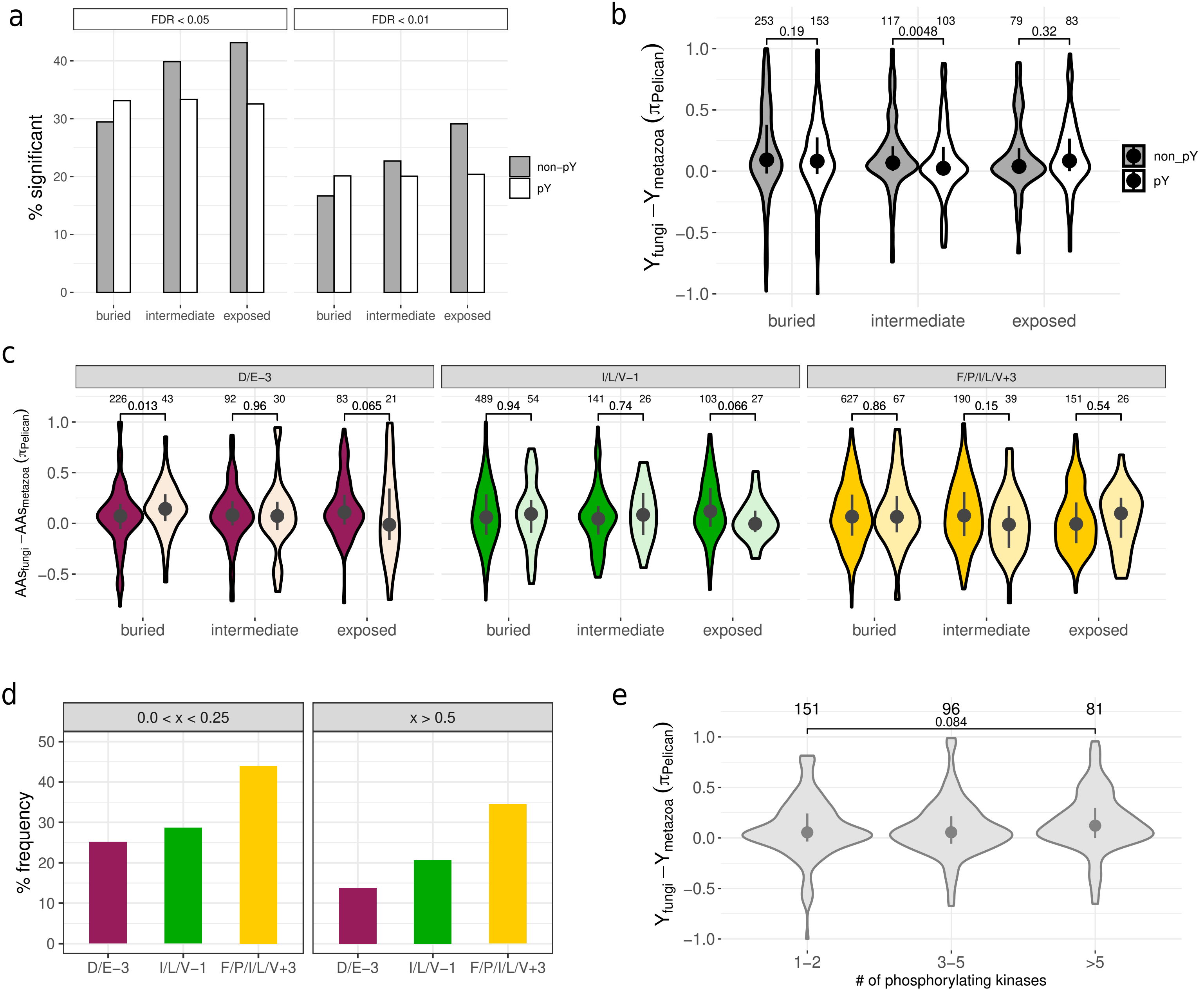

a
FDR < 0.05
FDR < 0.01
% significant
0
10
20
30
40
non−pY
pY
buried
intermediate
exposed
b
253 0.19 153
117 0.0048 103
79 0.32 83
Y_fungi − Y_metazoa (π_Pelican)
1.0
0.5
0.0
−0.5
−1.0
non_pY
pY
buried
intermediate
exposed
c
D/E−3
I/L/V−1
F/P/I/L/V+3
226 0.013 43
92 0.96 30
83 0.065 21
489 0.94 54
141 0.74 26
103 0.066 27
627 0.86 67
190 0.15 39
151 0.54 26
AAs_fungi − AAs_metazoa (π_Pelican)
1.0
0.5
0.0
−0.5
buried
intermediate
exposed
d
0.0 < x < 0.25
x > 0.5
% frequency
0
10
20
30
40
50
D/E−3
I/L/V−1
F/P/I/L/V+3
e
151
96
81
0.084
Y_fungi − Y_metazoa (π_Pelican)
1.0
0.5
0.0
−0.5
−1.0
1−2
3−5
>5
### of phosphorylating kinases
